# Discovery and description of gammanonin: a widely distributed natural product from Gammaproteobacteria

**DOI:** 10.1101/2024.08.13.607837

**Authors:** Jonas H Costa, Eva E Adams, Chad W Johnston

## Abstract

Antibiotics are essential for modern medicine, but their use drives the evolution of antimicrobial resistance (AMR) that limits the long-term efficacy of any one drug. To keep pace with AMR and preserve our ability to treat bacterial infections, it is essential that we identify antibiotics with new structures and targets that are not affected by clinical resistance. Historically, most developmental candidates for antibiotics have come from microbial natural products, as they feature chemical structures and biological activities that have been honed over millions of years of evolution. Unfortunately, as classical bioactivity screens for natural product discovery are blind to the pharmacological properties of their hits, they often identify molecules with functional groups that limit their utility as drugs. One prominent example is actinonin, an inhibitor of bacterial peptide deformylase (PDF) whose activity is dependent on a hydroxamate moiety associated with toxicity *in vivo*. The abundance of bacterial genomes now presents an opportunity for target-based natural product discovery, where biosynthetic pathways can be mined for molecules that possess desired activities but lack toxic moieties. Here, we use bioinformatics to lead a chemotype-sensitive, target-based search for natural product inhibitors of bacterial PDF that lacks the conserved and problematic metal chelating group. We describe the discovery, heterologous expression, biosynthesis, total synthesis, and activity of the molecule gammanonin: an apparent actinonin homologue from Gammaproteobacteria. Moving forward, we hope this chemotype and target-driven methodology will help to expedite the discovery of new leads for antibiotic development.

## Introduction

Antibiotics are keystone drugs that provide us an ability to fight or prevent bacterial infections, thus enabling invasive surgeries, immunosuppressive therapies, and other hallmarks of modern medicine. Unfortunately, the use of antibiotics also drives the development and dissemination of antimicrobial resistance (AMR) that limits their long-term viability^1^. As many approved antibiotics reuse chemical scaffolds or affect conserved targets and binding sites, emergent resistance mechanisms can simultaneously imperil multiple generations of drugs. Multi-drug resistant Gram-negative bacteria now pose a particularly daunting threat, as their outer membrane provides an intrinsic barrier that further limits treatment options^2^. The challenges of AMR highlight the need for continued antibiotic discovery, with an emphasis on targets and chemotypes not already impacted by clinical resistance^3^.

Historically, microbial natural products have been our most important source of antibiotics^4^, as these molecules feature privileged structures and activities that have evolved over millions of years^5^. Although bioactivity screening of microbial extracts has produced many of the antibiotic scaffolds still used today, such efforts have suffered from diminishing returns associated with oversampling, and from their tendency to uncover toxic hits with limited potential for development. A classic example is chloramphenicol, an antibiotic inhibitor of bacterial ribosomes discovered during a bioactivity screen in 1947. Though effective at fighting infections in humans, the *para*-nitrobenzene and dihaloalkane moieties of chloramphenicol produce toxic metabolic byproducts following oral or intravenous administration, leading to aplastic anemia that limits the use of this drug today^6,7^. More recent examples include actinonin^8^ and matlystatin^9^, hydroxamate inhibitors of bacterial peptide deformylase (PDF; **Figure 1a**): an essential enzyme responsible for removing formate from the N-formyl-methionine that initiates bacterial protein translation. Surprisingly, resistance to PDF-inhibitors is not achieved by mutation of this target, but through inactivating mutations in methionyl-tRNA formyltransferase, yielding bacteria with significantly reduced fitness and virulence^10^. Despite the promise of PDF as an antibiotic target, the hydroxamate present on natural inhibitors is metabolically unstable and mutagenic *in vivo*^6,11^. Actinonin derivatives that have undergone clinical trials thus feature more stable reverse hydroxamates^12^, though such molecules still had safety profiles less favorable than existing treatments such as linezolid.

**Figure 1.**
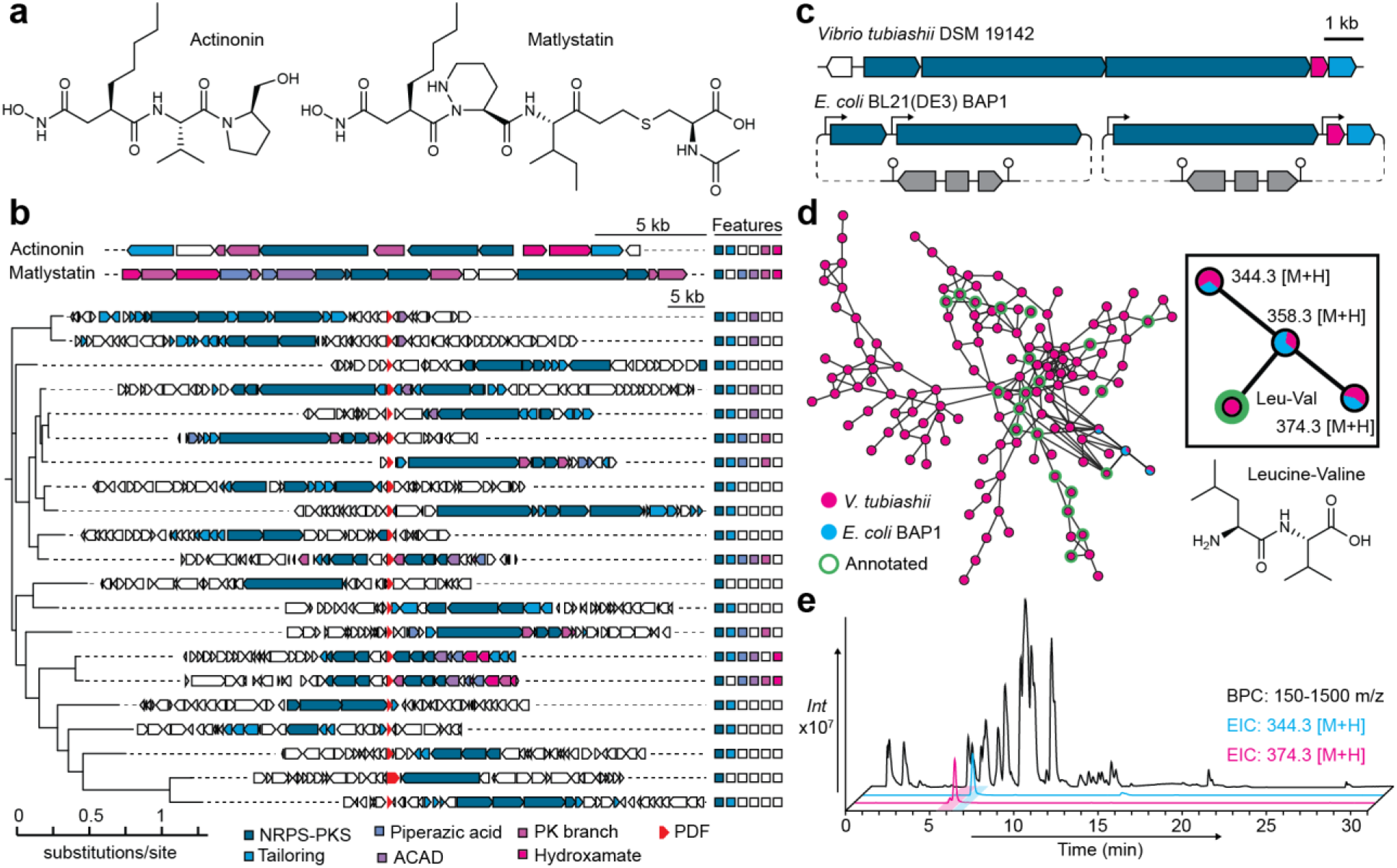
Resistance-guided discovery of novel genetic pathways for antibiotic inhibitors of peptide deformylase. (**a**) Actinonin and matlystatin are established antibiotic inhibitors of bacterial peptide deformylase (PDF). Both feature a hydroxamate that chelates a metal ion in the PDF active site. (**b**) ClusterScout was used to find biosynthetic gene clusters (BGCs) with nonribosomal peptide (NRPS) genes colocalized with an additional genomic copy of PDF, which may provide protection against nascent PDF inhibitory natural products. A BGC alignment was generated using CORASON. These BGCs have varying similarity to the pathways for actinonin and matlystatin. (**c**) A candidate BGC from Gammaproteobacteria was selected for further analysis. Plasmids were designed to enable heterologous expression in *E. coli* strain BAP1. (**d**) Molecular networking between a fractionated *V. tubiashii* extract and an extract from our heterologous expression strain identified a small series of peptides with apparent similarity to the annotated leucine-valine dipeptide. (**e**) Extracted ion chromatograms (EICs) of the two dominant ions identified in our molecular networking analysis, along with a base peak chromatogram (BPC) of the associated *E. coli* culture extract.

In the last decade, natural product discovery has experienced a renaissance driven by microbial genome sequencing^13^. Bio- and chemo-informatic methods can now use antibiotic biosynthetic gene clusters (BGCs) to guide the discovery of new molecules from diverse bacteria^14,15^. As antibiotic BGCs often contain self-protection genes that act as resistant copies of the associated target, molecules with specific activities can be mined directly from available genomes^16–18^. This methodology provides a valuable opportunity to revisit clinically untapped targets and identify developmental leads with superior pharmacological properties.

Here, we apply this target-centric mining approach to scan BGCs for hydroxamate-free inhibitors of bacterial peptide deformylase (PDF). Among other hits, this search revealed a novel BGC that is widespread in Gammaproteobacteria, encoding a molecule that resembles actinonin but lacks a clear metal chelating moiety. We report the identification, isolation, heterologous expression, total synthesis, biosynthesis, and activity of the new natural product gammanonin.

## Results

### Profiling BGCs to reveal candidate pathways for non-hydroxamate PDF inhibitors

To identify candidate BGCs for antibiotic inhibitors of bacterial PDF, we used the ClusterScout^19^ tool to search over 135,000 sequenced and annotated bacterial genomes from the Integrated Microbial Genomes (IMG) database^20^. Using nonribosomal peptide synthetase (NRPS) domains as hooks, we searched for BGCs with a co-localized PDF gene (<10kb away). This initial search returned 158 candidates, which could be further refined to 73 leads by eliminating isolated domains, BGCs for iron-binding siderophores, and pathways spuriously positioned next to the lone genomic copy of PDF. Using CORASON^21^, we grouped these leads into 21 distinct families of NRPS BGCs with a colocalized additional copy of PDF (**Figure 1b**), which could serve as a putative self-protection gene^18^. Among this series of BGCs, 18 were found in the prolific Gram positive Actinomycetes^22^, while a single candidate could be found in Gram negative Cyanobacteria, Alphaproteobacteria, and Gammaproteobacteria. Two BGCs (**Figure S1**. BGC12, 16) were closely related to matlystatin^9^, while another (BGC10) likely encodes the related molecule Sch-382583^23^, which lacks the hydroxamate moiety. Three BGCs (**Figure S1**. BGC1, 13, 21) were very closely related to the recently described pathway for lydiamycin^24^, which resembles Sch-382583 and includes a PDF gene of uninvestigated function. A full series of representative BGCs recovered from this search is available as a Supplementary Dataset, accompanied by an annotated CORASON alignment (**Figure S1**).

The most compelling lead to emerge from our analysis was the lone BGC found in Gammaproteobacteria, including several species of *Vibrio* and *Photorhabdus*. This minimal pathway features one polyketide synthase (PKS) module, three NRPS modules, a priming phosphopantetheinyl transferase (PPTase), a pathway-specific methyltransferase (MT), and critically, a PDF gene embedded in the biosynthetic operon (**Figure 1c, Table S1**). The predicted chemical product of this BGC is a polyketide-extended tripeptide – comparable to actinonin, but without a hydroxamate or other metal chelating group. This BGC is frequently flanked by transposons or phage-derived genes, suggesting it has been repeatedly mobilized and retained following horizontal gene transfer. Use of the pathway-specific MT gene in BLAST searches against the complete nucleotide collection of NCBI revealed instances of this BGC in other Gammaproteobacteria, including *Xenorhabdus* and *Brenneria*. Finally, this BGC has also been associated with intra-genus antibacterial activity in *Vibrio*^25^, suggesting its product should be an antibiotic that may be expressed in laboratory conditions. Given these promising features, we chose to investigate this BGC in the hopes of identifying a new, hydroxamate-free antibiotic inhibitor of bacterial PDF.

### Investigating of a candidate BGC from Gammaproteobacteria

We obtained *Vibrio tubiashii* DSM 19142 from the German Collection of Microorganisms and Cell Cultures (DSMZ) and attempted to repeat the culture and extraction conditions that had previously been reported to enrich for this unknown natural product^25^. Unfortunately, the reported UV signal (300 nm) for this product was absent and was furthermore inconsistent with the structure predicted from the biosynthetic machinery; a diagnostic mass was never reported. As genetic inactivation in *V. tubiashii* proved intractable, we chose to heterogously express this pathway in the *E. coli* BL21(DE3) derivative BAP1, which contains a genomically-integrated PPTase^26^. The BGC was cloned into two pET Duet plasmids, with inducible T7 promoters supporting the expression of each assembly-line gene as well as the co-transcribed PDF and MT (**Figure 1c**). BAP1 *E. coli* with this refactored pathway were grown in M9 minimal media, induced, and grown overnight at 37, 30, or 16°C. Cultures were centrifuged and the resulting cell pellets and supernatants were extracted and analyzed by high-resolution liquid chromatography-coupled mass spectrometry (HR-LCMS). To identify metabolites associated with BGC expression, we used molecular networking^27^ to compare our extracts of *V. tubiashii* and our engineered *E. coli*. A single series of peptide metabolites related to the GNPS annotated leucine-valine dipeptide was shared between *V. tubiashii* and BAP1 when induced at 16°C (**Figure 1de**). These three molecules appeared to be variably modified tripeptides present in very low concentrations in both the supernatant and cell pellet. As an alternative strategy, we developed an identical heterologous expression system using the matching BGC in *Photorhabdus asymbiotica* DSM 15149, which resulted in an identical product in a similarly low concentration (**Figure S2**). As heterologous expression of the *V. tubiashii* BGC provided better yields than the native organism and employed a minimal, chemically defined culture media, we pursued *E. coli* as a producer for all future work.

To identify which metabolite likely represented the final product of our BGC, we added stable isotopically labelled methionine and acetate, which should respectively reveal the involvement of the MT and the terminating module of the PKS (**Figure S3**). While the methionine methyl group was incorporated into all three metabolites, the PKS-associated acetate was only found in 374.3 [M+H] (**1**), suggesting the other molecules result from failed transmission between assembly-line enzymes. Adenylation domain substrate predictions^28,29^ indicated the likely incorporation of one L-leucine and two L-valine residues (**Table S2**), which could be corroborated by MS/MS of the 344.3 [M+H] tripeptide (**2**), though their order remained unclear. To resolve this, we used solid phase peptide synthesis (SPPS) to create tripeptide variants with leucine, valine, isoleucine, and N-methyl-valine at different positions. A synthetic tripeptide of N-methyl-L-valine-L-valine-L-leucine provided an identical retention time and fragmentation pattern to the natural tripeptide (**Figure S4**). A series of minor ions (358.3 [M+H]) could also be observed in extracts, appearing to correspond to valine-to-leucine/isoleucine substitutions by MS/MS. To resolve the structure of our complete natural product, we next grew, extracted, and processed 100 L of our engineered *E. coli* producer strain. Combined cell pellet and supernatant extracts were separated using size exclusion chromatography and semi-preparative LCMS to afford 7.8 mg of **2** and 1.8 mg of **1**, sufficient for structure elucidation by NMR. As expected, **2** matches our synthetic tripeptide, while **1** features a polyketide extended leucine diol reminiscent of the residue statine^30^, which is present in many natural protease inhibitors^31^. As this molecule resembles actinonin (**Figure 1a**) and is produced by Gammaproteobacteria, it was named gammanonin.

### Synthesis of gammanonin

To resolve the stereochemistry of the secondary alcohol in **1** and elucidate its absolute chemical structure, we established a total synthesis for gammanonin. As the N-methylated dipeptide could be readily obtained through SPPS using commercially available substrates, we envisioned accessing gammanonin by coupling this dipeptide to an appropriate isomer of the reduced statine residue (statinol). Synthesis of statinol followed an established route to statine^30,31^. Fmoc-L-leucine was converted to the methyl ester and reduced to the associated aldehyde, such that lithium-assisted Claisen condensation with ethyl acetate provided the *R*/*S* diastereomers of the Fmoc-L-statine ethyl ester. NMR analysis of Mosher esters^32^ was used to establish *S* as the dominant product (*ee*: 0.85), consistent with prior reports of statine synthesis^30^. Following the isolation of each diastereomer by LCMS, a second reduction and deprotection yielded pure statinols. Coupling the major isomer (L-*S*-statinol) with our dipeptide yielded a product with an identical retention time and MS/MS fragmentation pattern to gammanonin (**Figure 2a, Figure S5**). The apparent *R* isomer is observed in trace amounts in culture extracts.

**Figure 2.**
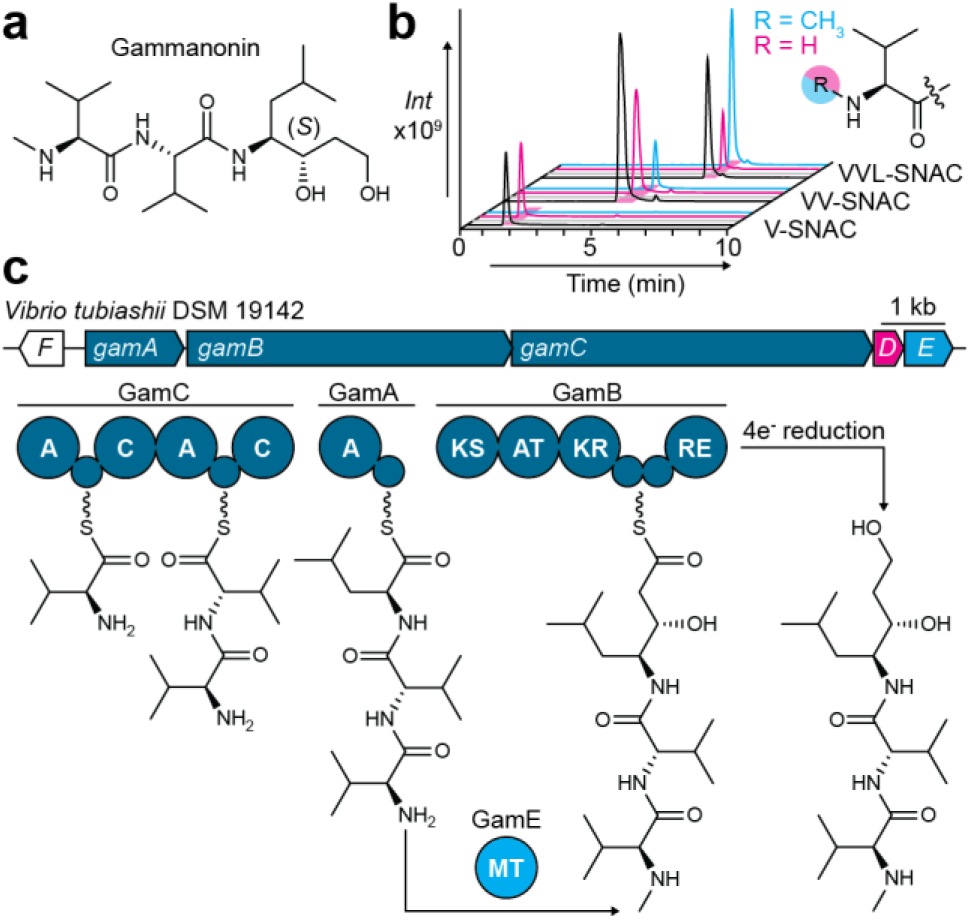
Structure and biosynthesis of gammanonin (**a**) The absolute structure of **1** was established by NMR and confirmed by total synthesis. (**b**) The substrate preference of the N-methyltransferase GamE was assessed using N-acetyl-cysteamine (SNAC) analogs of gammanonin biosynthetic intermediates. BPCs of enzymatic reactions are depicted (*black*) alongside EICs of unreacted (*magenta*) and methylated (*cyan*) SNAC peptides. GamE was found to preferentially N-methylate the tripeptide SNAC. (**c**) Proposed biosynthesis of gammanonin.

### Resolving the biosynthesis of gammanonin

The gammanonin BGC included several unusual features that we wished to resolve in order to propose an accurate biosynthetic scheme. Most notably, α-N-methylation is apparently achieved *in trans* by GamE, in contrast to nearly^33^ all bacterial nonribosomal peptides which feature *cis*-acting N-methyltransferase domains embedded in the NRPS^34^. The presence of the N-methylated tripeptide **2** indicated that GamE acts before the growing peptide is transferred to the PKS. To investigate the substrate preference of GamE, we synthesized N-acetyl-cysteamine (SNAC^35^) versions of each peptide intermediate, expressed and purified GamE, and performed methylation reactions *in vitro*. LCMS analysis of these reactions indicated that GamE cannot methylate individual amino acyl SNACs and preferentially acts on the tripeptide (**Figure 2b**). To assess the importance of this methylation to biosynthesis, we removed *gamE* from our expression plasmids, induced cultures carrying this modified system, and analyzed culture extracts by LCMS. In the absence of GamE, we did not observe **1, 2**, or unmethylated **2**, but rather only unmethylated **1** (**Figure S6**). The lack of signal for unmethylated **2** suggests this tripeptide may be degraded by the cell or that overexpression of *gamE* contributes to premature release of **2** from the assembly-line. Next, we sought to investigate the roles of the tandem acyl carrier protein (ACP) domains^36^ in the GamB PKS. As GamE works *in trans*, we suspected that one ACP may facilitate an interaction with GamE while the second may be used for polyketide extension of the tripeptide. To test this hypothesis, we replaced the active site serine residues in each of these ACPs with alanines, then induced, extracted, and analyzed cultures. Surprisingly, the inactivation of either ACP domain merely caused reduced yields of **1** and did not result in novel intermediates (**Figure S7**), suggesting these domains are essentially redundant^36^.

### Assessing the activity of gammanonin

Finally, we sought to investigate the biological activity of gammanonin and the role of the BGC-associated PDF enzyme GamD. First, we removed *gamD* from our expression plasmids and analyzed culture extracts to confirm that this gene is not involved in biosynthesis. As expected, the removal of gamD did not result in the production of new intermediates and had only a minor impact on gammanonin yields (**Figure S7**). Surprisingly, the removal of the assumed resistance gene *gamD* also did not appreciably impact the growth of *E. coli* following induction (**Figure S8**). To assess whether *E. coli* is intrinsically resistant to gammanonin, we overexpressed and purified the *E. coli* PDF enzyme (**Figure S9**) to directly test inhibition *in vitro*. This endpoint assay measures the capacity of PDF to liberate a primary amine from an N-formylated tripeptide substrate, which is then detected by the fluorogenic reporter molecule fluorescamine^37^. Using this method, we did not observe inhibition of *E. coli* PDF by gammanonin, in contrast to an actinonin positive control (**Figure 3a**). To identify organisms that may be sensitive to gammanonin, we tested this molecule in whole cell microdilution assays against a panel of Gram positive and negative bacteria (**Table 1**). As the gammanonin BGC had previously been associated with intra-genus antagonism in *Vibrio*^25^, we included a number of *Vibrio* species in this survey. Collectively, gammanonin showed limited antibacterial activity against a small set of Gram negative bacteria, including *Acinetobacter pittii, Vibrio crassotrae*, and oddly, *Photorhabdus asymbiotica*, which also has the gammanonin BGC. By contrast, actinonin displayed antibacterial activity against most tested strains. While poor whole cell antibiotic activity has been observed with other non-hydroxamate PDF inhibitors such as Sch-382583^23^, this may reflect poor uptake rather than poor target engagement. Given prior reports of this BGC being associated with anti-*Vibrio* activity^25^ and the presence of the apparent self-protection gene *gamD* in the biosynthetic operon, we next tested whether gammanonin could inhibit PDF enzymes from *Vibrio* species. We expressed and purified house-keeping PDF proteins from *V. crassotrae* and *V. tubiashii* (Ga0077872_1535), along with GamD (**Figure S9**). An additional isolated and non-conserved putative PDF from *V. tubiashii* (Ga0077872_1694) was also purified but failed to yield substantial activity in laboratory conditions (**Figure S10**). As with the *E. coli* enzyme, we failed to observe inhibition of PDF activity by gammanonin, in contrast to an actinonin positive control (**Figure 3**). That GamD is conserved, co-expressed with biosynthetic genes, does not have an apparent role in biosynthesis, and functions as a conventional peptide deformylase supports a potential role as a self-protection gene, though an appropriate context for antibacterial activity remains elusive.

**Table 1.**
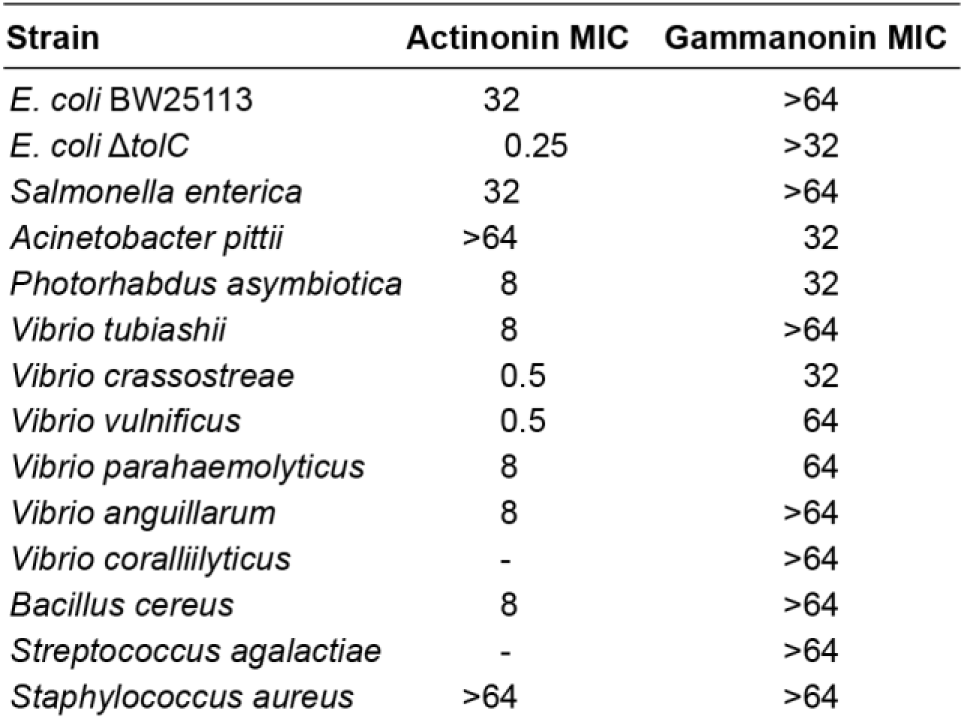
Minimum inhibitory concentrations (MICs; μg ml^-1^) of gammanonim and actinonin.

**Figure 3.**
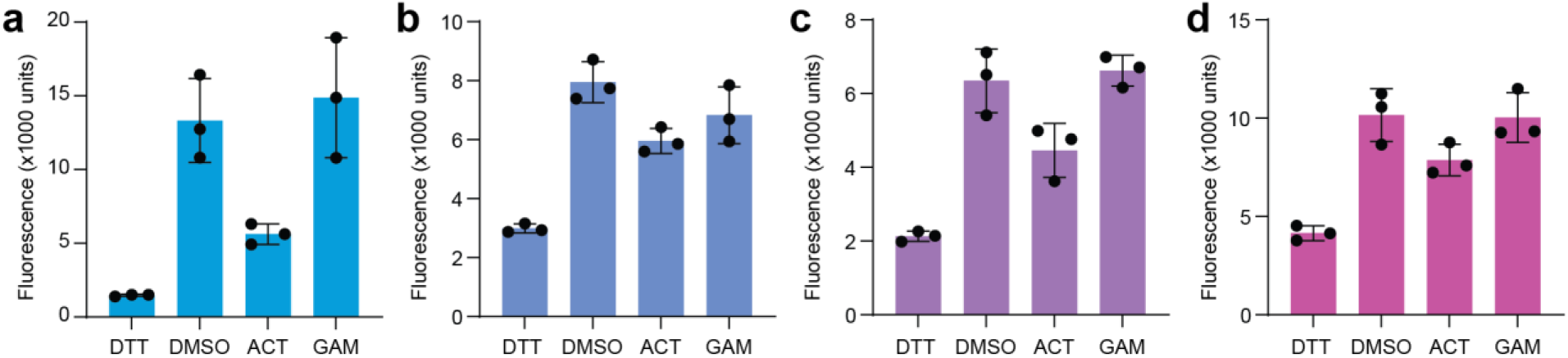
Assaying the inhibition of peptide deformylase enzymes by gammanonin and actinonin. The fluorogenic molecule fluorescamine was used to quantify the liberation of primary amines from an N-formylated peptide substrate by PDF enzymes, including those from (**a**) *E. coli*, (**b**) *V. crassotrae*, (**c**) *V. tubiashii* (locus tag: Ga0077872_1535), and (**d**) GamD. All reactions contained 25 mM HEPES (pH 7.5) with 1 mM formyl-Met-Ala-Ser substrate with 4% DMSO. Activity was assessed in the presence of 25 mM DTT (negative control), DMSO alone (uninhibited positive control), 100 µM actinonin (ACT), or 100 µM gammanonin (GAM). Reactions were performed in triplicate for 30 minutes at 37°C (*E. coli*) or 30°C (*Vibrio*) prior to addition of fluorescamine to a final concentration of 60 µg mL^-1^. Error bars represent standard deviation.

## Discussion

Microbial natural products feature chemical structures and biological activities honed through natural selection to provide a fitness advantage to their producer, yielding evolved molecules that serve as vital leads for pharmaceuticals^4^. However, natural products are not evolved for human use, and often feature chemical moieties that limit their viability as drugs. In this work, we present a chemotype-sensitive approach to natural product genome mining, with the goal of identifying antibacterial inhibitors of peptide deformylase that lack the problematic hydroxamate^11^ group found in prior developmental leads like actinonin^8,9^. Using this approach, we successfully identified a series of BGCs for suitable antibiotics (**Figure 1b**), including a reported BGC for lydiamycin^24^ and a likely pathway for the known PDF inhibitor Sch-382583^23^. The gammanonin BGC stands out among these hits, as it is found in several Gram negative Gammaproteobacteria and features a simple, highly conserved operon structure that includes a co-transcribed PDF gene. Boundaries for this BGC are also defined by its apparent acquisition through horizontal gene transfer, as evidenced by flanking genes linked to transposons and bacteriophage. Oddly, dedicated transcription factors or efflux pumps are not associated with this pathway, suggesting its phylogenetic distribution may be limited to bacteria that can complement these functions. The presence of this pathway in *Vibrio* species is also unusual, as this genus has only previously been known to produce nonribosomal peptide siderophores^38–40^ and the similarly widely distributed antibiotic andrimid^41^. This biosynthetic pathway also includes rare components, such as the *trans*-acting N-methyltransferase GamE. To our knowledge, this is the first instance of on-assembly-line α-N-methylation being achieved in *trans*, as this is typically achieved by cis-acting domains^34^ or following release from the NRPS^33^. Further profiling of substrate specificity will be useful to assess the potential of GamE to act as an adaptable biocatalyst for peptide N-methylation.

Although the gammanonin BGC encodes a functional PDF gene that presumably acts in self-protection, we were unable to observe inhibition of diverse PDF enzymes by gammanonin or substantial antibacterial activity against a diverse panel of microbes. This lack of activity is surprising, as the gammanonin BGC had previously been identified in a report from 2012 describing antagonism between *Vibrio* strains^25^. In that study, transposon mutagenesis of a *V. ordalii* isolate identified this BGC as a driver of antagonistic activity. While fractionation of whole culture extracts by HPLC identified a peak the authors claimed was responsible for the reported activity, the molecule was not isolated, antibacterial tests were not shown, and mass spectrometry data was not provided. While the described extract fractionation does enrich for gammanonin, the reported UV absorbance of 300 nm is inconsistent with our structure and biosynthetic pathway. It remains possible that this UV-active metabolite observed previously coelutes with gammanonin. Despite our initial bioactivity results, it remains possible that gammanonin functions as an antibacterial PDF inhibitor, and that an appropriate sensitive organism is yet to be identified.

## Supporting Information

The Supporting Information, including methods, supplementary figures and tables, and NMR spectra, is available as a separate file. An additional supplementary data file including BGC sequences for Figure 1b is also available.

### Notes

The authors declare no competing financial interest.

## Acknowledgements

This work was supported by seed funds from the Baylor College of Medicine and by a recruitment grant from the Cancer Prevention and Research Institute of Texas (CPRIT; RR210066). CWJ is a CPRIT Scholar in Cancer Research.

## Notes

### Competing Interest Statement

The authors have declared no competing interest.

## References

(1) Cook, M. A.; Wright, G. D. The Past, Present, and Future of Antibiotics. Sci. Transl. Med. 2022, 14 (657), eabo7793.

(2) Theuretzbacher, U.; Blasco, B.; Duffey, M.; Piddock, L. J. V. Unrealized Targets in the Discovery of Antibiotics for Gram-Negative Bacterial Infections. Nat Rev Drug Discov 2023, 22 (12), 957–975.

(3) Miethke, M.; Pieroni, M.; Weber, T.; Brönstrup, M.; Hammann, P.; Halby, L.; Arimondo, P. B.; Glaser, P.; Aigle, B.; Bode, H. B.; Moreira, R.; Li, Y.; Luzhetskyy, A.; Medema, M. H.; Pernodet, J.-L.; Stadler, M.; Tormo, J. R.; Genilloud, O.; Truman, A. W.; Weissman, K. J.; Takano, E.; Sabatini, S.; Stegmann, E.; Brötz-Oesterhelt, H.; Wohlleben, W.; Seemann, M.; Empting, M.; Hirsch, A. K. H.; Loretz, B.; Lehr, C.-M.; Titz, A.; Herrmann, J.; Jaeger, T.; Alt, S.; Hesterkamp, T.; Winterhalter, M.; Schiefer, A.; Pfarr, K.; Hoerauf, A.; Graz, H.; Graz, M.; Lindvall, M.; Ramurthy, S.; Karlén, A.; Van Dongen, M.; Petkovic, H.; Keller, A.; Peyrane, F.; Donadio, S.; Fraisse, L.; Piddock, L. J. V.; Gilbert, I. H.; Moser, H. E.; Müller, R. Towards the Sustainable Discovery and Development of New Antibiotics. Nat Rev Chem 2021, 5 (10), 726–749.

(4) Wright, G. D. Opportunities for Natural Products in 21 st Century Antibiotic Discovery. Nat. Prod. Rep. 2017, 34 (7), 694–701.

(5) Lewis, K.; Lee, R. E.; Brötz-Oesterhelt, H.; Hiller, S.; Rodnina, M. V.; Schneider, T.; Weingarth, M.; Wohlgemuth, I. Sophisticated Natural Products as Antibiotics. Nature 2024, 632 (8023), 39–49.

(6) Blagg, J. Structural Alerts for Toxicity. In Burger’s Medicinal Chemistry and Drug Discovery; 2003; pp 301–334.

(7) Stepan, A. F.; Walker, D. P.; Bauman, J.; Price, D. A.; Baillie, T. A.; Kalgutkar, A. S.; Aleo, M. D. Structural Alert/Reactive Metabolite Concept as Applied in Medicinal Chemistry to Mitigate the Risk of Idiosyncratic Drug Toxicity: A Perspective Based on the Critical Examination of Trends in the Top 200 Drugs Marketed in the United States. Chem. Res. Toxicol. 2011, 24 (9), 1345–1410.

(8) Wolf, F.; Leipoldt, F.; Kulik, A.; Wibberg, D.; Kalinowski, J.; Kaysser, L. Characterization of the Actinonin Biosynthetic Gene Cluster. ChemBioChem 2018, 19 (11), 1189–1195.

(9) Leipoldt, F.; Santos-Aberturas, J.; Stegmann, D. P.; Wolf, F.; Kulik, A.; Lacret, R.; Popadić, D.; Keinhörster, D.; Kirchner, N.; Bekiesch, P.; Gross, H.; Truman, A. W.; Kaysser, L. Warhead Biosynthesis and the Origin of Structural Diversity in Hydroxamate Metalloproteinase Inhibitors. Nat Commun 2017, 8 (1), 1965.

(10) Margolis, P. S.; Hackbarth, C. J.; Young, D. C.; Wang, W.; Chen, D.; Yuan, Z.; White, R.; Trias, J. Peptide Deformylase in Staphylococcus Aureus: Resistance to Inhibition Is Mediated by Mutations in the Formyltransferase Gene. Antimicrob Agents Chemother 2000, 44 (7), 1825–1831.

(11) Smith, G. F. Designing Drugs to Avoid Toxicity. In Progress in Medicinal Chemistry; Elsevier, 2011; Vol. 50, pp 1–47.

(12) Corey, R.; Naderer, O. J.; O’Riordan, W. D.; Dumont, E.; Jones, L. S.; Kurtinecz, M.; Zhu, J. Z. Safety, Tolerability, and Efficacy of GSK1322322 in the Treatment of Acute Bacterial Skin and Skin Structure Infections. Antimicrob Agents Chemother 2014, 58 (11), 6518–6527.

(13) Scherlach, K.; Hertweck, C. Mining and Unearthing Hidden Biosynthetic Potential. Nat Commun 2021, 12 (1), 3864.

(14) Johnston, C. W.; Skinnider, M. A.; Wyatt, M. A.; Li, X.; Ranieri, M. R. M.; Yang, L.; Zechel, D. L.; Ma, B.; Magarvey, N. A. An Automated Genomes-to-Natural Products Platform (GNP) for the Discovery of Modular Natural Products. Nat Commun 2015, 6 (1), 8421.

(15) Skinnider, M. A.; Johnston, C. W.; Edgar, R. E.; Dejong, C. A.; Merwin, N. J.; Rees, P. N.; Magarvey, N. A. Genomic Charting of Ribosomally Synthesized Natural Product Chemical Space Facilitates Targeted Mining. Proc. Natl. Acad. Sci. U.S.A. 2016, 113 (42), E6343–E6351.

(16) Culp, E. J.; Sychantha, D.; Hobson, C.; Pawlowski, A. C.; Prehna, G.; Wright, G. D. ClpP Inhibitors Are Produced by a Widespread Family of Bacterial Gene Clusters. Nat Microbiol 2022, 7 (3), 451–462.

(17) Culp, E. J.; Waglechner, N.; Wang, W.; Fiebig-Comyn, A. A.; Hsu, Y.-P.; Koteva, K.; Sychantha, D.; Coombes, B. K.; Van Nieuwenhze, M. S.; Brun, Y. V.; Wright, G. D. Evolution-Guided Discovery of Antibiotics That Inhibit Peptidoglycan Remodelling. Nature 2020, 578 (7796), 582–587.

(18) Mungan, M. D.; Alanjary, M.; Blin, K.; Weber, T.; Medema, M. H.; Ziemert, N. ARTS 2.0: Feature Updates and Expansion of the Antibiotic Resistant Target Seeker for Comparative Genome Mining. Nucleic Acids Research 2020, 48 (W1), W546–W552.

(19) Hadjithomas, M.; Chen, I.-M. A.; Chu, K.; Huang, J.; Ratner, A.; Palaniappan, K.; Andersen, E.; Markowitz, V.; Kyrpides, N. C.; Ivanova, N. N. IMG-ABC: New Features for Bacterial Secondary Metabolism Analysis and Targeted Biosynthetic Gene Cluster Discovery in Thousands of Microbial Genomes. Nucleic Acids Res 2017, 45 (D1), D560–D565.

(20) Palaniappan, K.; Chen, I.-M. A.; Chu, K.; Ratner, A.; Seshadri, R.; Kyrpides, N. C.; Ivanova, N. N.; Mouncey, N. J. IMG-ABC v.5.0: An Update to the IMG/Atlas of Biosynthetic Gene Clusters Knowledgebase. Nucleic Acids Research 2020, 48 (D1), D422–D430.

(21) Navarro-Muñoz, J. C.; Selem-Mojica, N.; Mullowney, M. W.; Kautsar, S. A.; Tryon, J. H.; Parkinson, E. I.; De Los Santos, E. L. C.; Yeong, M.; Cruz-Morales, P.; Abubucker, S.; Roeters, A.; Lokhorst, W.; Fernandez-Guerra, A.; Cappelini, L. T. D.; Goering, A. W.; Thomson, R. J.; Metcalf, W. W.; Kelleher, N. L.; Barona-Gomez, F.; Medema, M. H. A Computational Framework to Explore Large-Scale Biosynthetic Diversity. Nat Chem Biol 2020, 16 (1), 60–68.

(22) Gavriilidou, A.; Kautsar, S. A.; Zaburannyi, N.; Krug, D.; Müller, R.; Medema, M. H.; Ziemert, N. Compendium of Specialized Metabolite Biosynthetic Diversity Encoded in Bacterial Genomes. Nat Microbiol 2022, 7 (5), 726–735.

(23) Chu, M.; Mierzwa, R.; He, L.; Xu, L.; Gentile, F.; Terracciano, J.; Patel, M.; Miesel, L.; Bohanon, S.; Kravec, C.; Cramer, C.; Fischman, T. O.; Hruza, A.; Ramanathan, L.; Shipkova, P.; Chan, T.-M. Isolation and Structure Elucidation of Two Novel Deformylase Inhibitors Produced by Streptomyces Sp. Tetrahedron Letters 2001, 42 (21), 3549–3551.

(24) Libis, V.; MacIntyre, L. W.; Mehmood, R.; Guerrero, L.; Ternei, M. A.; Antonovsky, N.; Burian, J.; Wang, Z.; Brady, S. F. Multiplexed Mobilization and Expression of Biosynthetic Gene Clusters. Nat Commun 2022, 13 (1), 5256.

(25) Cordero, O. X.; Wildschutte, H.; Kirkup, B.; Proehl, S.; Ngo, L.; Hussain, F.; Le Roux, F.; Mincer, T.; Polz, M. F. Ecological Populations of Bacteria Act as Socially Cohesive Units of Antibiotic Production and Resistance. Science 2012, 337 (6099), 1228–1231.

(26) Pfeifer, B. A.; Admiraal, S. J.; Gramajo, H.; Cane, D. E.; Khosla, C. Biosynthesis of Complex Polyketides in a Metabolically Engineered Strain of E. Coli. Science 2001, 291 (5509), 1790–1792.

(27) Aron, A. T.; Gentry, E. C.; McPhail, K. L.; Nothias, L.-F.; Nothias-Esposito, M.; Bouslimani, A.; Petras, D.; Gauglitz, J. M.; Sikora, N.; Vargas, F.; Van Der Hooft, J. J. J.; Ernst, M.; Kang, K. B.; Aceves, C. M.; Caraballo-Rodríguez, A. M.; Koester, I.; Weldon, K. C.; Bertrand, S.; Roullier, C.; Sun, K.; Tehan, R. M.; Boya P. C. A.; Christian, M. H.; Gutiérrez, M.; Ulloa, A. M.; Tejeda Mora, J. A.; Mojica-Flores, R.; Lakey-Beitia, J.; Vásquez-Chaves, V.; Zhang, Y.; Calderón, A. I.; Tayler, N.; Keyzers, R. A.; Tugizimana, F.; Ndlovu, N.; Aksenov, A. A.; Jarmusch, A. K.; Schmid, R.; Truman, A. W.; Bandeira, N.; Wang, M.; Dorrestein, P. C. Reproducible Molecular Networking of Untargeted Mass Spectrometry Data Using GNPS. Nat Protoc 2020, 15 (6), 1954–1991.

(28) Blin, K.; Shaw, S.; Kloosterman, A. M.; Charlop-Powers, Z.; van Wezel, G. P.; Medema, M. H.; Weber, T. antiSMASH 6.0: Improving Cluster Detection and Comparison Capabilities. Nucleic Acids Research 2021, 49 (W1), W29–W35.

(29) Skinnider, M. A.; Johnston, C. W.; Gunabalasingam, M.; Merwin, N. J.; Kieliszek, A. M.; MacLellan, R. J.; Li, H.; Ranieri, M. R. M.; Webster, A. L. H.; Cao, M. P. T.; Pfeifle, A.; Spencer, N.; To, Q. H.; Wallace, D. P.; Dejong, C. A.; Magarvey, N. A. Comprehensive Prediction of Secondary Metabolite Structure and Biological Activity from Microbial Genome Sequences. Nat Commun 2020, 11 (1), 6058.

(30) Rittle, K. E.; Homnick, C. F.; Ponticello, G. S.; Evans, B. E. A Synthesis of Statine Utilizing an Oxidative Route to Chiral .Alpha.-Amino Aldehydes. J. Org. Chem. 1982, 47 (15), 3016–3018.

(31) Yang, S.; Zhang, W.; Ding, N.; Lo, J.; Liu, Y.; Clare-Salzler, M. J.; Luesch, H.; Li, Y. Total Synthesis of Grassystatin A, a Probe for Cathepsin E Function. Bioorganic & Medicinal Chemistry 2012, 20 (15), 4774–4780.

(32) Du, Y. E.; Cui, J.; Cho, E.; Hwang, S.; Jang, Y.-J.; Oh, K.-B.; Nam, S.-J.; Oh, D.-C. Serratiomycins D1–D3, Antibacterial Cyclic Peptides from a Serratia Sp. and Structure Revision of Serratiomycin. J. Nat. Prod. 2024, 87 (5), 1330–1337.

(33) Shi, R.; Lamb, S. S.; Zakeri, B.; Proteau, A.; Cui, Q.; Sulea, T.; Matte, A.; Wright, G. D.; Cygler, M. Structure and Function of the Glycopeptide N-Methyltransferase MtfA, a Tool for the Biosynthesis of Modified Glycopeptide Antibiotics. Chemistry & Biology 2009, 16 (4), 401–410.

(34) Süssmuth, R. D.; Mainz, A. Nonribosomal Peptide Synthesis—Principles and Prospects. Angew Chem Int Ed 2017, 56 (14), 3770–3821.

(35) Ehmann, D. E.; Trauger, J. W.; Stachelhaus, T.; Walsh, C. T. Aminoacyl-SNACs as Small-Molecule Substrates for the Condensation Domains of Nonribosomal Peptide Synthetases. Chemistry & Biology 2000, 7 (10), 765–772.

(36) Rahman, A. S.; Hothersall, J.; Crosby, J.; Simpson, T. J.; Thomas, C. M. Tandemly Duplicated Acyl Carrier Proteins, Which Increase Polyketide Antibiotic Production, Can Apparently Function Either in Parallel or in Series. Journal of Biological Chemistry 2005, 280 (8), 6399–6408.

(37) Yang, N.; Sun, C. The Inhibition and Resistance Mechanisms of Actinonin, Isolated from Marine Streptomyces Sp. NHF165, against Vibrio Anguillarum. Front. Microbiol. 2016, 7. 1467.

(38) Keating, T. A.; Marshall, C. G.; Walsh, C. T. Vibriobactin Biosynthesis in Vibrio Cholerae: VibH Is an Amide Synthase Homologous to Nonribosomal Peptide Synthetase Condensation Domains. Biochemistry 2000, 39 (50), 15513–15521.

(39) Di Lorenzo, M.; Poppelaars, S.; Stork, M.; Nagasawa, M.; Tolmasky, M. E.; Crosa, J. H. A Nonribosomal Peptide Synthetase with a Novel Domain Organization Is Essential for Siderophore Biosynthesis in Vibrio Anguillarum. J Bacteriol 2004, 186 (21), 7327–7336.

(40) Zane, H. K.; Naka, H.; Rosconi, F.; Sandy, M.; Haygood, M. G.; Butler, A. Biosynthesis of Amphi-Enterobactin Siderophores by Vibrio Harveyi BAA-1116: Identification of a Bifunctional Nonribosomal Peptide Synthetase Condensation Domain. J. Am. Chem. Soc. 2014, 136 (15), 5615–5618.

(41) Wietz, M.; Mansson, M.; Gotfredsen, C. H.; Larsen, T. O.; Gram, L. Antibacterial Compounds from Marine Vibrionaceae Isolated on a Global Expedition. Marine Drugs 2010, 8 (12), 2946–2960.

