## Supplementary Information for "Discovery and description of gammanonin: a widely distributed natural product from Gammaproteobacteria"

### Table of Contents

Materials and methods …………………………………………………………………………….….. 2-9

Supplemental tables ………………………………………………………………………….…….. 10

Supplemental figures …………………………………………………………………………….. 11-16

NMR spectra and tables ……………………………………………………………………………. 17-47

Supplemental references ………………………………………………………………………… 48-49

### Methods

#### General biological methods

Plasmid cloning was performed in NEB Turbo cells (New England Biolabs) grown in 2x YT media (VWR) with appropriate antibiotics at 37°C. Plasmids were assembled with USER cloning as previously described[1], amplifying associated sequences using Q5U Hot Start DNA polymerase (New England Biolabs). Competent bacterial cells were prepared using the TSS method as previously described[1]. Antibiotics were used at the following concentrations: chloramphenicol, 30 μg $mL^{-1}$; kanamycin, 25 μg $mL^{-1}$. Protein expression and heterologous production of natural products was performed in *E. coli* BL21(DE3) or its derivative BAP1[2] (Kerafast) as this strain features a genomically-integrated phosphopantetheinyl transferase. Bacterial culture for protein expression and purification was performed in terrific broth (VWR).

*Vibrio tubiashii* DSM 19142, *Photorhabdus asymbiotica* DSM 15149, and *Acinetobacter pittii* DSM 25618 were obtained from the German Resource Centre for Biological Material (DSMZ). *Vibrio* isolates used for antibacterial assays were obtained as a kind gift from the Seafood Safety laboratory of Prof. John Schwarz at Texas A&M Galveston. EcM2.1.dtolC was a gift from George Church (Addgene # 64053). *Streptococcus agalactiae* and *Bacillus cereus* isolates were a kind gift from Prof. Katy Patras at the Baylor College of Medicine. *Staphylococcus aureus* NCTC 8317 was obtained from the National Collection of Type Cultures (NCTC). *Salmonella enterica* ATCC BAA-2568 was obtained from the American Type Culture Collection (ATCC). *V. tubiashii* and *Vibrio* test strains were grown in Marine media (Difco) at 30°C. *P. asymbiotica, A. pittii, S. enterica*, *S. aureus*, *B. cereus*, *S. agalactiae*, *E. coli* BW25113, and EcM2.1.dtolC were cultured in cation adjusted Mueller-Hinton broth at 30°C.

Antimicrobial assays were performed in 96-well culture plates, using a serial dilution of gammanonin in DMSO from 64 μg $mL^{-1}$ to 0.125 μg $mL^{-1}$, including no drug and no bacteria controls. Bacterial inoculum was prepared from overnight cultures grown in Marine broth (for *Vibrio spp.*) or CAMHB at 30°C. 3 μL of saturated culture was diluted into 3 mL of fresh media, and 1 μL was added via multichannel pipette to each well of the serial dilution. Plates were covered with gas permeable membranes (Breathe-Easy; Diversified Biotech) and grown at 30°C with shaking in a Synergy H1 microplate reader (BioTek), recording absorbance at 600 nm every 30 minutes for 24 hours.

#### General Chemical Methods

NMR spectra were acquired on a Bruker 600 & 800 MHz NMR spectrometer using DMSO-$d_6$ as a solvent. NMR spectra were analyzed using TopSpin software. Acetonitrile for liquid chromatography was obtained from VWR (HiSolv; UPLC/HPLC grade) while HPLC-grade water was obtained from an in-house Millipore unit. Solvents for synthesis were obtained from Sigma-Aldrich. High resolution mass spectrometry coupled liquid chromatography (HR-LCMS) was performed on a Thermo Scientific Orbitrap Exploris 120 mass spectrometer coupled with a Thermo Scientific HPLC featuring a Binary Pump (VC-P10-A). Analytical scale liquid chromatography employed a Thermo Scientific Accucore C18 column (3 mm × 150 mm × 2.6 μm) heated to 40ºC, with mobile phases of 0.1% formic acid (A) and acetonitrile (B). For analysis and purification of the Fmoc-L-leucine aldehyde synthetic intermediate (**4**), 0.1% TFA was used in aqueous and organic phases. Semi-preparative compound isolation employed a Thermo Scientific HyperSil Gold (10 mm × 150 mm × 5 μm) heated to 40ºC. All the operation and spectra analysis were conducted using Thermo Chromeleon (version 7.3.1) and Free Style (version 1.8 SP2).

*Mass spectrometer conditions*: positive mode with *m*/*z* range of 150–1500, Orbitrap resolution of 120000, RF lens of 70%, spray voltage (H-ESI) at 3.5 kV, sheath gas at 30 arb, aux gas at 4 arb, sweep gas at 0 arb, ion transfer tube temperature at 320 °C, vaporizer temperature at 75ºC. *MS/MS conditions*: Orbitrap resolution of 60000, normalized collision energy type, HCD collision energy of 30, 40 and 50%, 4 scans (Top N). *Standard analytical method* (A:B) : 95:5 held for 1.2 min, gradient up to 0:100 in 16.8 min, 0:100 held for 8 min, gradient down to 95:5 in 0.2 min, 95:5 held for 8.8 min; total run time was 35 min; flow rate of 0.8 mL $min^{-1}$. Injection volume of 5 μL.

#### Bioinformatic identification and visualization of candidate BGCs for PDF-inhibitory natural products

Genome mining activities were achieved using the ClusterScout tool from Integrated Microbial Genomes (IMG) Atlas of Biosynthetic gene Clusters (IMG-ABC v5)[3,4]. A library of 409357 microbial BGCs predicted by antiSMASH v5 were scanned for colocalized protein families (Pfams) with the following parameters:

Pfam hooks: pfam00501, pfam00550, pfam00668, pfam01327
Hooks required: 4
Essential pfam: pfam01327
Maximum distance between hooks: 10000
Extend boundaries by: 5000
Minimum distance from scaffold edge: 1000

This search included Pfams associated with nonribosomal peptide synthetase (NRPS) genes (pfam00501, pfam00550, pfam00668) as well as peptide deformylase (pfam01327), returning 158 candidate BGCs as a result. Each BGC was manually investigated to verify that the observed PDF gene was not the sole genomic copy, that the BGC did not encode a siderophore (as evidenced by colocalized iron metabolism genes), and that it was not a degraded or poorly assembled BGC fragment. The 73 BGCs (**Fig S1**) that remained following this filtering represented 21 distinct pathways (**Fig 1b**). DNA sequences containing each BGC were obtained from IMG and run through antiSMASH[5] to obtain standardized GenBank file outputs and assess whether the product of a given pathway was known. GenBank files for each BGC were then processed by BigScape-CORASON[6] to generate BGC alignments. Parameters were maintained at default values, with changes for **Fig S1** being [e_value = 1E-10; cluster_radio = 100] and changes for **Fig 1b** being [e_value = 1E-4; cluster_radio = 400 e_value = 1E-4; cluster_radio = 400].

An Excel spreadsheet containing the annotated complete output of the ClusterScout search is available as a Supplementary Data File. Individual FASTA files for each BGC featured in **Fig 1b** are available as a Supplementary Data Set.

**Heterologous expression of the gammanonin BGC**

To enable heterologous expression of the gammanonin BGC, genes were cloned directly from the genomic DNA of *V. tubiashii* or *P. asymbiotica* into T7 expression cassettes of pET Duet plasmids pACYC (*gamA*, *gamB*) and pCOLA (*gamC*, *gamDE*). A single colony of *E. coli* BAP1 carrying pACYC and pCOLA plasmids was inoculated into 100 mL of M9 Minimal Medium with kanamycin and chloramphenicol and incubated at 37°C with shaking at 220 rpm for overnight growth. From this culture, 10 mL was used to inoculate 2.8 L Erlenmeyer flasks containing 1 L of M9 supplemented with antibiotics. Cultures were incubated at 37°C and 220 rpm until $OD_{600}$ 0.5-0.6, when IPTG was added. Cultures were maintained at 220 rpm overnight. Different temperatures (16°C, 24°C, 30°C, or 37°C) and IPTG concentrations (1 mM or 0.1 mM) were tested, identifying overnight incubation at 16°C and induction with 0.1 mM IPTG as optimal for gammanonin production.

Following induction and overnight growth, cultures were centrifuged at 4000 rpm, reserving both cell pellets and supernatants. Pellets were extracted with methanol, aided by submersion in an ultrasonic bath for 1 hour. Supernatants were extracted by addition of 20 g $L^{-1}$ adsorbent Amberlite XAD7 resin (Thermo Scientific), which was allowed to mix for 4 hours at 200 rpm. Resin was recovered by Buchner funnel vacuum filtration, washed with water, and eluted with methanol. Cell pellet and resin methanol extracts were dried under vacuum by rotary evaporation, resuspended in methanol, and stored at -20°C. In total, 100 L of culture was prepared and extracted.

**Purification of gammanonin**

Liquid chromatography with size exclusion resin was used as an initial means of purifying gammanonin and associated products. Crude extracts resuspended in methanol were applied to an open column containing 100 g Sephadex LH-20 (Cytiva) resin in a methanol mobile phase. A Gilson FC 204 Fraction Collector was used to fractionate the column eluate. Fractions were analyzed via HR-LCMS, and those containing the metabolites of interest were combined, dried under vacuum by rotary evaporation, and stored at -20ºC. Next, fractions were resuspended in methanol and subjected to semi-preparative HPLC.

*Chromatographic conditions*: Mobile phase was 0.1% formic acid (A) and acetonitrile (B); Eluent profile (A:B): 100:0 held for 1.2 min, gradient up to 0:100 (curve 8) in 44.8 min, 0:100 held for 6 min, gradient down to 100:0 in 0.5 min, 100:0 held for 7.5 min; total run time was 60 min; flow rate of 5 mL $min^{-1}$. Injection volume of 500 µL. Gammanonin eluted at 15 min.

**Extraction and fractionation of *Vibrio* culture**

The growth, extraction, and fractionation of cultures of *Vibrio* was performed following the protocol established by Cordero et al[7]. In short, a colony of *V. tubiashii* was inoculated into 3 mL marine broth (Difco) and grown overnight at 30°C as a small-scale overnight culture. From this, 1 mL was inoculated into 1 L of marine broth in a 2.8 L Erlenmeyer flask and incubated at 25°C and 150 rpm for 24 hours. Afterwards, 20 g $L^{-1}$ Amberlite XAD7 resin (Thermo Scientific) was added to each culture and allowed to mix for a further 24 hours. Resin was recovered by Buchner funnel vacuum filtration, washed with water, and eluted with methanol. Cell pellet and resin methanol extracts were dried under vacuum by rotary evaporation, resuspended in methanol, and stored at -20°C. This crude extract was resuspended in 20 mL of 1:1:1 mixture of trimethylpentane, ethyl acetate, and methanol, mixed with 50 g of silica gel mesh 230-400, dried under vacuum by rotary evaporation, and loaded onto a 300 mL bed volume of silica gel in a 600 mL Buchner funnel. Six fractions were collected, washing the silica with 300 mL of trimethylpentane, 1:1 trimethylpentane and ethyl acetate, ethyl acetate, 1:1 ethyl acetate and acetone, acetone, and methanol. All fractions were dried under rotary evaporation and analyzed by HR-LCMS.

**Dereplication of MS data and Molecular Networking**

Molecular networks of HR-LCMS/MS data from cell pellet and supernatant extracts were created using the online workflow on the GNPS[8] website (https://gnps.ucsd.edu). MS/MS data were converted to mzXML file format using MSConvert[9] package (Version 3.0.21015-d21ed5bee, Proteowizard Software Foundation, USA) and uploaded to GNPS platform using WinSCP (Version 6.3.3). Data was filtered to remove all MS/MS fragment ions within +/- 17 Da of the precursor m/z. MS/MS spectra were window filtered by choosing only the top 6 fragment ions in a +/- 50 Da window throughout the spectrum. The precursor ion mass tolerance was set to 0.02 Da and a MS/MS fragment ion tolerance of 0.02 Da. A network was then created where edges were filtered to have a cosine score above 0.78 and more than 5 matched peaks. Further, edges between two nodes were kept in the network if and only if each of the nodes appeared in each other's respective top 8 most similar nodes. Finally, the maximum size of a molecular family was set to 200, and the lowest scoring edges were removed from molecular families until the molecular family size was below this threshold. The spectra in the network were then searched against GNPS' spectral libraries. The library spectra were filtered in the same manner as the input data. All matches kept between network spectra and library spectra were required to have a score above 0.7 and at least 6 matched peaks.

Spectral networks were visualized with Cytoscape[10] (Version 3.9.0). A complete molecular network for **Fig 1d** is available at https://gnps.ucsd.edu/ProteoSAFe/status.jsp?task=6b9f779ba02f48fb9b90fc809c2f0cdc

**Incorporation of isotopically labelled precursors**

To assess the incorporation of isotopically labelled precursors into gammanonin metabolites, 100 mL M9 cultures of *E. coli* BAP1 with pACYC and pCOLA plasmids were grown and induced with IPTG as described. Following induction, cultures were supplemented with 2 mM L-valine, 2 mM L-leucine, and either 4 mM sodium acetate ($^{13}C_2$; Sigma-Aldrich) or 4 mM L-methionine-(methyl-$d_3$). Cultures were grown and extracted following standard methods and analyzed by HR-LCMS to assess incorporation.

**Expression and purification of gammanonin N-methyltransferase GamE**

The *gamE* gene was amplified directly from a colony of *V. tubiashii* with USER primers, purified by gel extraction, and assembled into plasmid pET-28a, which features an N-terminal $His_6$ tag for protein purification. After sequencing, this plasmid was transformed into *E. coli* BL21(DE3). A single colony was grown overnight at 37°C and 300 rpm in 3 mL 2xYT media with appropriate antibiotics before being added into 1 L terrific broth. Cultures were grown at 37°C and 220 rpm to an OD 0.6 before being induced by the addition of IPTG to final concentration of 0.5 mM. Induced cultures were incubated at 16°C and 180 rpm overnight. The following morning, cultures were centrifuged at 5000 rpm for 15 minutes at 4°C. Supernatants were decanted and cell pellets were stored at -80°C until further use. *All subsequent purification steps took place at 4°C*. Cell pellets were resuspended in 30 mL lysis buffer (50mM $K_2HPO_4$, 500mM NaCl, 5mM BME, 20mM imidazole, 10% glycerol, pH 8) with half of a SIGMAFAST protease inhibitor tablet (Sigma-Aldrich) per cell pellet and sonicated 10s-on 45s-off for a total "on" time of 10 minutes. Lysate was centrifuged at 15,000 rpm for 45 minutes and the supernatant was loaded onto a column with 2mL Ni-NTA resin pre- equilibrated with lysis buffer. After the full supernatant volume was loaded, columns were washed with 8 column volumes of wash buffer (50 mM $K_2HPO_4$, 500 mM NaCl, 5mM BME, 20 mM imidazole, pH 8). The proteins were then eluted with 15 mL elution buffer (50 mM $K_2HPO_4$, 500 mM NaCl, 5 mM BME, 300 mM imidazole, 10% glycerol, pH 8). Eluted proteins were dialyzed in 1 L phosphate buffer (50 mM $K_2HPO_4$, 500 mM NaCl, 1 mM DTT, pH 8) overnight. Dialyzed protein solutions were then applied to Amicon 10,000 MW centrifugal filters and concentrated to a volume of 1 mL. Purification was verified using SDS-PAGE. Purified GamE was stored at -80°C with 25% glycerol. A C-terminal $His_6$-tagged variant of GamE was also purified with similar expression, purification, and assay results. This purification method is adapted from Yang et al[11].

**Methyltransferase assay**

An assay to assess the substrate specificity of the GamE N-methyltransferase was derived from a previously described method[12]. In brief, this assay tests the ability of GamE to utilize S-adenosyl methionine (SAM) and methylate aminoacyl N-acetylcysteamine thioester substrates (aminoacyl SNACs[13]). A stock of purified GamE was recovered from storage at -80°C and thawed slowly on ice. Assays took place in 100 µL 100 mM Tris-HCl (pH 7.5), with 6.4 µM GamE, 0.8 mM aminoacyl-SNAC, and 2.4 mM SAM, incubated overnight at 25°C. Negative controls omitted GamE. Following overnight incubation, reactions were quenched with 50 µL methanol, vortexed, and centrifuged at 15,000 rpm for 5 minutes to remove any precipitated protein. Reactions were analyzed by HR-LCMS, identifying methylated SNACs via the associated +14 Da product.

**Expression and purification of peptide deformylase enzymes**

*E. coli* BL21 (DE3) cells harboring peptide deformylase genes assembled into the pET28a N-terminal $His_6$-tag vector were grown in 4 mL 2xYT broth and 25 µg $mL^{-1}$ kanamycin. After overnight growth, cultures were added to 1L terrific broth supplied with antibiotics and grown at 37°C with shaking at 220 rpm to an OD 600 of 0.8. Cultures were removed from the incubator and protein overexpression was induced with the addition of IPTG to a final concentration of 0.5 mM and 0.1 mM nickel chloride[14]. Cultures were placed in a 25°C incubator and allowed to incubate overnight shaking at 150 rpm. Cells were harvested by centrifugation at 5000 rpm for 15 minutes. The supernatant were discarded, and cell pellets were stored at -80°C until further use.

Pellets were thawed over ice with 30 mL lysis buffer (50 mM HEPES, pH 7.5, 0.5 M KCl, 10% glycerol, 0.1% Triton X-100, 5 mM imidazole, 1 mM TCEP)[15]. All steps going forward took place at 4°C. Cells were sonicated for 10 minutes, 10s on with 45s off time between each pulse. To remove cell debris, lysate was centrifuged for 30 minutes at 12000 rpm. The supernatant was loaded onto 2 mL Ni-NTA resin pre-equilibrated with lysis buffer. After the supernatant was fully loaded, column was washed with 100 mL lysis buffer and 50 mL TG buffer: (50 mM HEPES, pH 7.5, 0.5 M KCl, 5 mM imidazole, and 1 mM TCEP). Proteins were eluted in a 50 mL volume gradient of imidazole (20-320mM) in TG buffer. Eluted fractions were dialyzed overnight in 1 L dialysis buffer (25 mM HEPES, pH 7.5, 150 mM KCl, 1 mM TCEP). Dialyzed fractions were concentrated using Amicon 3 kDa centrifugal filter units centrifuged at 4200 rpm to a volume of ~1 mL. Successful protein purification was verified using SDS-PAGE.

**Peptide deformylase assay**

An enzymatic assay for peptide deformylase (PDF) activity followed conditions described by Yang et al.[16] Briefly, this assay measures the liberation of a primary amine from an N-formylated peptide substrate by PDF, using the fluorogenic molecule fluorescamine to quantify free amines using a microplate reader.

Reactions were performed in 100 µL total volume, with 25 mM HEPES, pH 7.5, 1 mM formyl-Met-Ala-Ser substrate (fMAS; generated in-house by SPPS and purified by HPLC), with 30 nM PDF, 100 µM inhibitor or equivalent volume DMSO. Total DMSO concentration for all conditions was 4%. *Vibrio* PDF assay reactions were supplemented with 0.4 M NaCl. Reagents were added to a black 96 well plates with clear bottoms and incubated for 30 minutes at 37°C (*E. coli* PDF) or 30°C (*Vibrio* PDFs). Fluorescamine was added to a final concentration of 60 µg $mL^{-1}$ and fluorescence was monitored in a BioTek Synergy H1 microplate reader with excitation at 390 nm and emission at 470 nm. 25 mM DTT was used to inactivate PDFs in negative control wells to assess background fluorescence generated by lysine residues in PDF.

**Mosher ester synthesis**

Synthesis of Mosher esters followed the established protocol[17]. 3 mg of the major Fmoc statine ethyl ester isomer (**5**) was transferred to each of two 20 mL scintillation vials, lyophilized overnight, then dissolved in 1 mL anhydrous pyridine. Next, 30 µL of R- or S-α-methoxy-α-(trifluoromethyl) phenylacetyl chloride (MTPA-Cl) was added to either vial and allowed to react with stirring at room temperature for 2 hours. Reactions were quenched with 100 µL methanol, dried under vacuum by rotary evaporation, resuspended in methanol, and purified by semi-preparative HPLC. The delta values ($\Delta\delta$S-R) of the signals around the stereogenic center was assigned by analyzing $^1$H NMR and $^1$H-$^1$H COSY NMR spectra. Based on this comparison, we assigned the alcohol of the major Fmoc statine ethyl ester (**5**) as the *S* isomer.

**Synthesis of N-acetyl-cysteamine (SNAC) peptide substrates**

Synthesis of aminoacyl SNAC substrates followed the standard protocol established by Ehmann DE. et al[13]. Boc-protected peptides were generated through standard solid phase peptide synthesis as described above, incorporating Boc-L-valine as the final component and eluting from resin with 20% hexafluoro-isopropanol (HFIP) in DCM. To a scintillation vial was added 1 eq. Boc-protected peptide (Boc-L-valine, Boc-L-valine-L-valine, or Boc-L-valine-L-valine-L-leucine; 0.3 mmol), 1.2 eq HOBt, and 1.2 eq DCC, along with in THF (5 mL). Following a brief incubation, 1.2 eq N-acetyl-cysteamine was added and stirred for 45 minutes at room temperature. Afterwards, 0.5 eq. of K2CO3 was added and the reaction was allowed to proceed with stirring overnight. Reactions were partitioned between ethyl acetate and water, dried with $MgSO_4$, filtered, and concentrated under vacuum by rotary evaporation. SNAC peptides were purified by preparative reverse phase HPLC.

**Gammanonin synthesis**

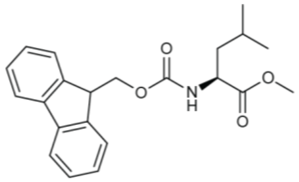

The scheme for generating the statine ethyl ester is derived from Rittle et al[18] and Yang et al[19]. In a 100 mL round bottom flask with a stir bar, Fmoc-leucine (500 mg) was resuspended in methanol (10 mL). A few drops of glacial sulfuric acid were added to this solution, which was then refluxed under $N_2$ at 80°C overnight. This reaction was dried under vacuum by rotary evaporation to yield a clear oil that was sampled for LCMS analysis, demonstrating complete conversion to the methyl ester **3**.

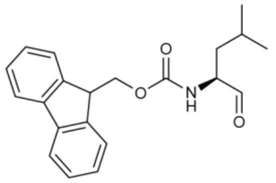

Fmoc-leucine methyl ester **3** (500 mg; 1.36 mmol) was dissolved in THF (20 mL) in a 100 mL round bottom flask with a stir bar under $N_2$ before being cooled to -78°C. To this solution, 5 eq. of diisobutylaluminum hydride (DIBALH; 1 M in cyclohexane) was added dropwise by syringe. This reaction was stirred at -78°C for 2 hours before being quenched by adding a small volume of ethyl acetate. The quenched reaction was mixed with a saturated solution of Rochelle salt (40 mL), allowed to warm to room temperature, and stirred for 2 hours. This solution was extracted twice with ethyl acetate, dried with $MgSO_4$, filtered, and concentrated under vacuum by rotary evaporation to yield 440 mg an off-yellow oil. LCMS analysis of this oil indicated a composition of Fmoc-leucine aldehyde **4** (>80%) along with a small quantity of unreacted starting material and the fully reduced alcohol. As this product will slowly isomerize at ambient temperatures[18], this oil was used directly for synthesis of the statine ethyl ester **5**.

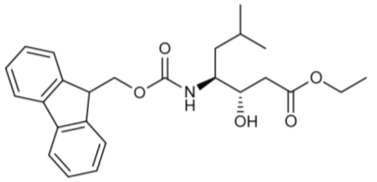

A 100 mL round bottom flask with stir bar under $N_2$ was loaded with 10 mL THF and cooled to -78°C. Next, 5 eq. of lithium bis(trimethylsilyl)amide (LiHMDS; 1 M in THF; 6.5 mL) was added and stirred to cool. 6 eq. of anhydrous ethyl acetate (8 mmol; 0.8 mL) was added dropwise and mixed for 1 hour to generate a lithio ethyl acetate reagent. Fmoc leucine aldehyde (440 mg; 1.3 mmol) was dissolved in 5 mL THF and cooled to -80°C before being added dropwise to this solution and then stirred for 3 hours at -78°C under $N_2$. The reaction was quenched by adding a small volume of ethyl acetate and then allowed to warm to room temperature. This solution was partitioned between ethyl acetate and water, recovering the organic phase and washing the aqueous phase twice with additional ethyl acetate. The combined organic phases were dried with $MgSO_4$, filtered, and concentrated under vacuum by rotary evaporation to yield an off-yellow oil. LCMS analysis revealed two isomers of Fmoc-statine ethyl ester as the major products. Two rounds of preparative reverse-phase HPLC was used to obtain the major S-isomer **5** (69 mg; 162 μmol; consistent with prior reports[18]) and the minor R-isomer **6** (6 mg; 14 μmol); (*ee*: 0.85).

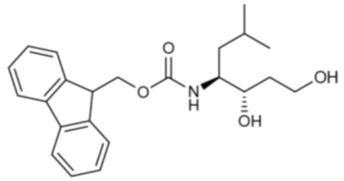

The major Fmoc-statine ethyl ester isomer (**5**) was dissolved in 10 mL THF, moved to a 100 mL round bottom flask with stir bar under $N_2$, and cooled to 0°C. 5 eq. of DIBALH (1 M in cyclohexane) was added dropwise to this solution and stirred for 1 hour before being quenched by the addition of a small volume of ethyl acetate. The quenched reaction was mixed with a saturated solution of Rochelle salt (20 mL), allowed to warm to room temperature, and stirred for 2 hours. This solution was extracted twice with ethyl acetate, dried with $MgSO_4$, filtered, and concentrated under vacuum by rotary evaporation. LCMS analysis confirmed complete conversion to the alcohol (Fmoc-*S*-L-statinol; **7**).

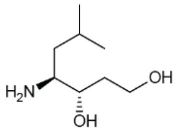

Fmoc-*S*-L-statinol was deprotected by dissolving in 5 mL DMF with 20% piperidine, shaking in a scintillation vial for 1 hour at room temperature. Removal of Fmoc was confirmed by LCMS. DMF was removed by lyophilization. Residual piperidine was removed by purifying *S*-L-statinol (**8**) by preparative reverse phase HPLC. Recovery and yield of **8** was limited, resulting in only ~1 mg of pure material for NMR analysis and peptide coupling from an initial 500 mg of Fmoc-L-leucine.

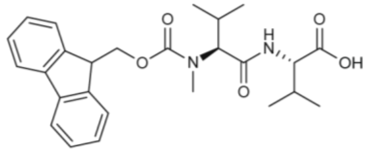

300 mg of 2-chlorotrityl resin (loading capacity calculated to 0.45 mmol $g^{-1}$) was added to a snap-cap polypropylene column (Poly-Prep Chromatography Column; Bio-Rad). The resin was swollen in 6 mL DCM, then drained. To the swollen resin was added 3 eq. of Fmoc-L-valine (0.4 mmol; 150 mg) in 5 mL DCM, with trace DMF to assist in solubilization. To this mixture was added 9 eq. of N,N-diisopropylethylamine (DIPEA; 250 µL). The vessel was capped, and loading was allowed to proceed with shaking at room temperature for 2 hours. After loading, the reaction vessel was drained, and the resin was washed with 5 mL DCM (x2) and 5 mL DMF (x2). Deprotection of the resin-bound peptide was achieved by the addition of 5 mL DMF with 20% piperidine (x2), mixing for 5 minutes before washing with 5 mL DMF (x2), 5 mL DCM (x2), and 5 mL DMF (x2). Next, 3 eq. of Fmoc-L-N-methylvaline (ChemImpex; 0.4 mmol; 95 mg) was dissolved in 5 mL DMF with 255 mg HATU and 95 µL DIPEA. This solution was allowed to mix for 5 minutes at room temperature before being added to the resin and mixed for 45 minutes at room temperature. Following incubation, the reaction was drained, and the resin washed with 5 mL DMF (x2) and 5 mL DCM (x4). The Fmoc-protected dipeptide was removed from the resin by addition of 5 mL TFA cleavage solution (95% TFA, 5% $H_2O$, 5% triisopropylsilane), shaking at room temperature for 30 minutes. The eluate was dried under nitrogen, yielding 100 mg of Fmoc-protected dipeptide (**9**).

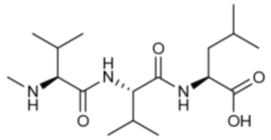

300 mg of 2-chlorotrityl resin (loading capacity calculated to 0.45 mmol $g^{-1}$) was added to a snap-cap polypropylene column (Poly-Prep Chromatography Column; Bio-Rad). The resin was swollen in 6 mL DCM, then drained. To the swollen resin was added 3 eq. of Fmoc-L-leucine (0.4 mmol; 150 mg) in 5 mL DCM, with trace DMF to assist in solubilization. To this mixture was added 9 eq. of N,N-diisopropylethylamine (DIPEA; 250 µL). The vessel was capped, and loading was allowed to proceed with shaking at room temperature for 2 hours. After loading, the reaction vessel was drained, and the resin was washed with 5 mL DCM (x2) and 5 mL DMF (x2). Deprotection of the resin-bound peptide was achieved by the addition of 5 mL DMF with 20% piperidine (x2), mixing for 5 minutes before washing with 5 mL DMF (x2), 5 mL DCM (x2), and 5 mL DMF (x2). Next, 3 eq. of Fmoc-L-valine (0.4 mmol; 95 mg) was dissolved in 5 mL DMF with 255 mg HATU and 95 µL DIPEA. This solution was allowed to mix for 5 minutes at room temperature before being added to the resin and mixed for 45 minutes at room temperature. Following incubation, the reaction was drained, and the resin washed as before. Deprotection and coupling steps were repeated with Fmoc-L-N-methylvaline (ChemImpex; 0.4 mmol; 95 mg). Prior to elution, the tripeptide was deprotected by addition of 5 mL DMF with 20% piperidine (x2), followed by washing in 5 mL DMF (x2) and 5 mL DCM (x4). Elution was achieved by addition of 5 mL TFA cleavage solution (95% TFA, 5% $H_2O$, 5% triisopropylsilane), shaking at room temperature for 30 minutes. The eluate was dried under nitrogen, yielding 100 mg of the gammanonin tripeptide (**2**).

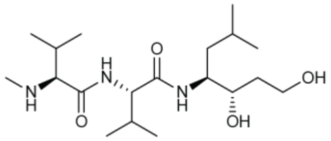


To a scintillation vial containing 1 eq. of Fmoc-dipeptide was added 1 mL DMF. To a second vial containing 1.1 eq of L-*S*-statinol (<1 mg) was added 3 mL DMF, 3 eq. HATU, and 3 eq. DIPEA. Following incubation for 5 minutes at room temperature, the solutions were combined and allowed to react with shaking for 2 hours. The reaction was monitored by LCMS. Following tripeptide formation and consumption of free dipeptide, the product was deprotected by the addition of 1 mL piperidine. Following a 1-hour incubation with shaking at room temperature, DMF was removed by lyophilization, and synthetic gammanonin (**1**) was compared to the natural molecule by analytical LCMS (**Fig S5**).

**Table S1. Components of the gam biosynthetic gene cluster.** Closest characterized homologs were identified through BLAST searches in the UniProtKB-SwissProtdatabase (June 2024).

| Protein | Size (aa) | Putative function | Closest characterized homolog (%AA ID) | UniProtKB |
|---|---|---|---|---|
| GamF | 248 | 4'-phosphopantetheinyl transferase | Demethylmenaquinone methyltransferase (36%) | B1VEN4.1 |
| GamA | 558 | NRPS (A-T) | Tyrocidine synthase 3 (32%) | O30409.1 |
| GamB | 1849 | PKS (KS-AT-KR-T-T-RE) | Beta-ketoacyl-acyl-carrier-protein synthase I (33%) | Q7TXL6.1 |
| GamC | 2051 | NRPS (A-T-C-A-T-C) | Linear gramicidin synthase subunit C (30%) | Q70LM5.1 |
| GamD | 170 | Peptide deformylase | Peptide deformylase 2 (60%) | Q87I22.1 |
| GamE | 274 | Methyltransferase | Demethylmenaquinone methyltransferase (36%) | B1VEN4.1 |

**Table S2. Substrate predictions for core biosynthetic proteins**

| Protein | A-domain residues (NRPS-PKS) | Predicted substrate antiSMASH | Predicted substrate PRISM | Observed substrate |
|---|---|---|---|---|
| GamA | **D A F A Y G C V** | L-Leucine | L-Valine | L-Leucine |
| GamB | - | Malonyl-CoA | Malonyl-CoA | Malonyl-CoA |
| GamC | **D A L W M G G T**<br>**D A L L L G G T** | L-Valine<br>L-Valine | L-Phenylalanine<br>L-Valine | L-Valine<br>L-Valine |

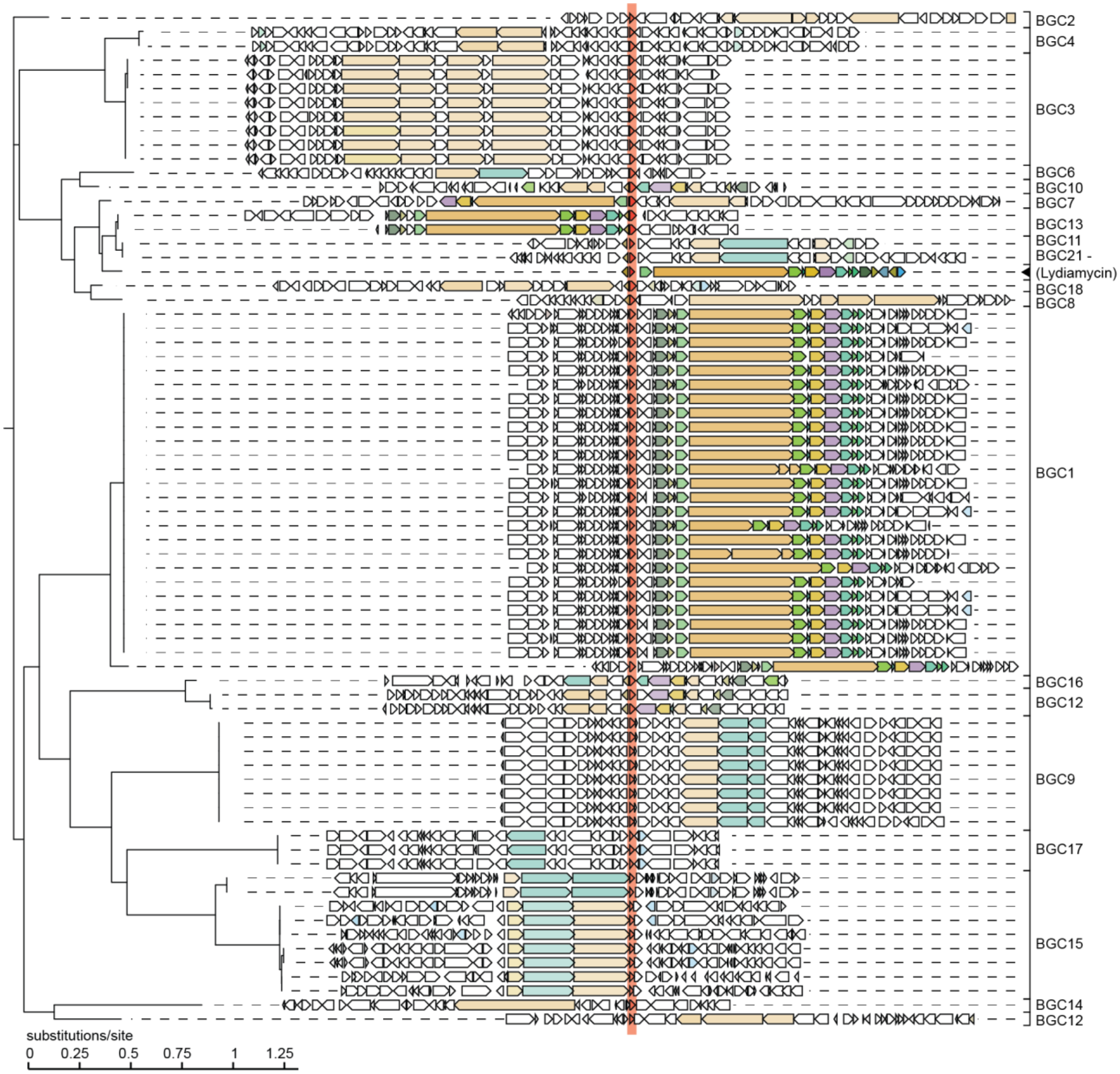


**Figure S1. Nonribosomal peptide synthetase (NRPS) pathways that feature colocalized peptide deformylase genes.** ClusterScout[3] was used to identify biosynthetic gene clusters (BGCs) in the Integrated Microbial Genomes database that featured colocalized NRPS and PDF genes. After removing spurious hits, CORASON[6] was used to align the remaining 73 BGCs into 21 distinct families. Representative BGCs for each family are available as FASTA files that can be analyzed by AntiSmash[5] or PRISM[20]. PDF genes in each BGC are highlighted in red, while the remaining genes feature default coloring from CORASON.

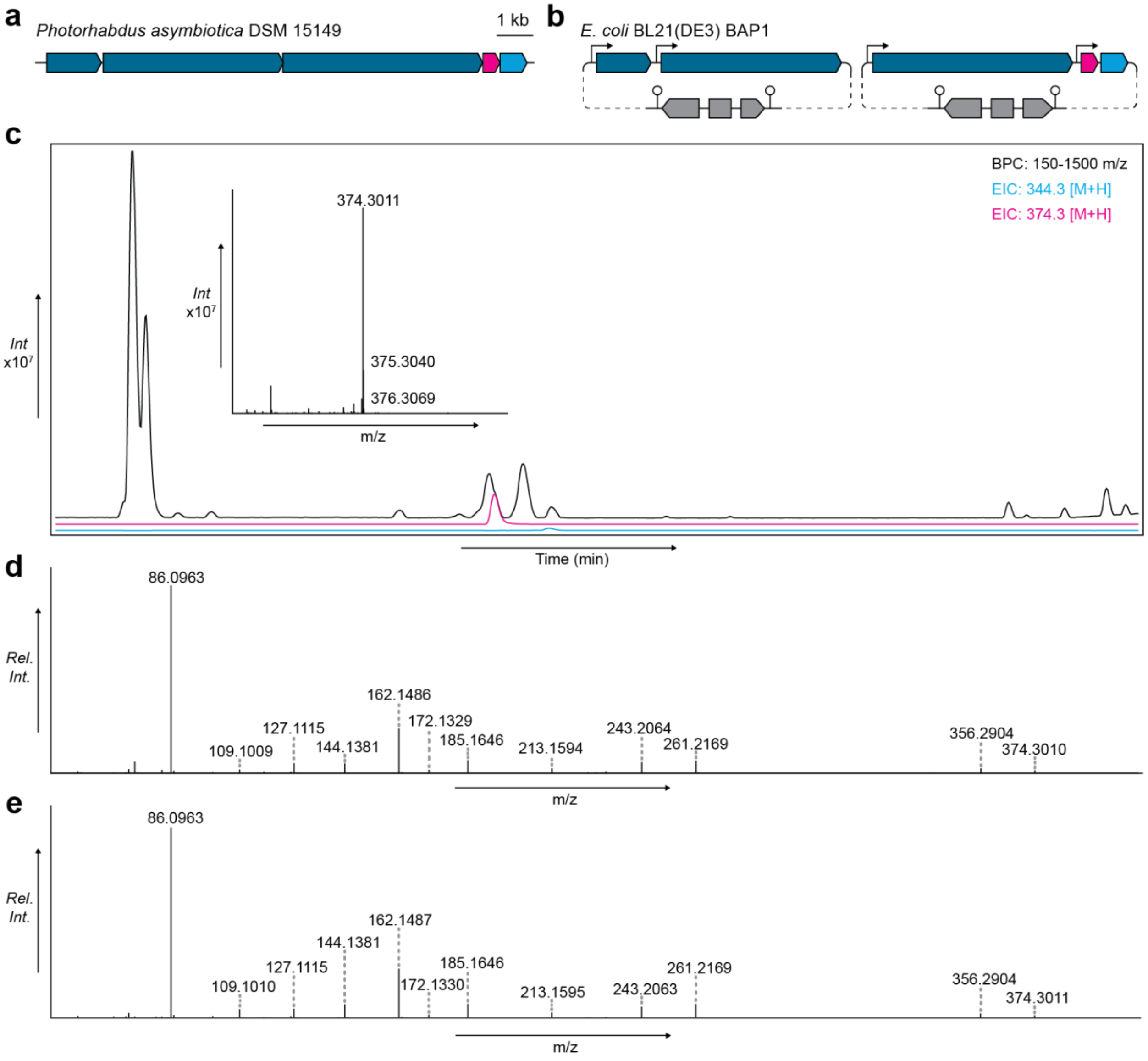


**Figure S2. Heterologous expression of a BGC from *Photorhabdus* yields an identical product. a.** A BGC from *Photorhabdus* that was also identified from our bioinformatic investigation is homologous to the BGC under investigation from *Vibrio*. This includes NRPS/PKS genes (dark blue), a peptide deformylase (magenta), and a methyltransferase (cyan). **b.** Plasmid designs for the heterologous expression of this pathway in *E. coli* BAP1. **c.** LCMS analysis of a cell pellet extract of the *E. coli* BAP1 heterologous expression strain, displaying a base peak chromatogram (BPC) and extracted ion chromatograms (EICs) for ions 374.3 [M+H] (gammanonin) and 344.3 [M+H] (gammanonin tripeptide). **d.** MS/MS fragmentation pattern for the 374.3 [M+H] ion observed during heterologous expression of the *Photorhabdus* pathway. **e.** MS/MS fragmentation pattern for the 374.3 [M+H] ion observed during heterologous expression of the *Vibrio* pathway.

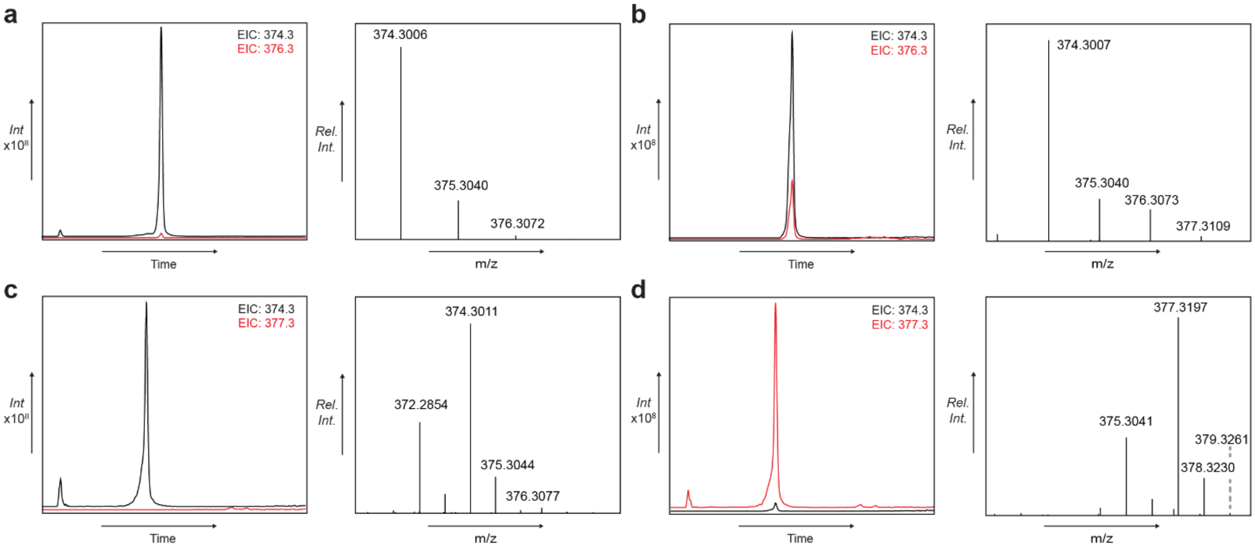


**Figure S3. Incorporation of isotopically labelled precursors into gammanonin. a.** Extracted ion chromatogram (EIC) for ions 374.3 [M+H] (gammanonin) and 376.3 [M+H] from cultures without label. Isotopic distribution of gammanonin is provided to the right. **b.** EIC for ions 374.3 [M+H] (gammanonin) and 376.3 [M+H] from cultures with $^{13}C_2$ acetate. Isotopic distribution of gammanonin is provided to the right. **c.** EIC for ions 374.3 [M+H] (gammanonin) and 377.3 [M+H] from cultures without label. Isotopic distribution of gammanonin is provided to the right. **b.** EIC for ions 374.3 [M+H] (gammanonin) and 377.3 [M+H] from cultures with L-methionine-(methyl-$d_3$). Isotopic distribution of gammanonin is provided to the right

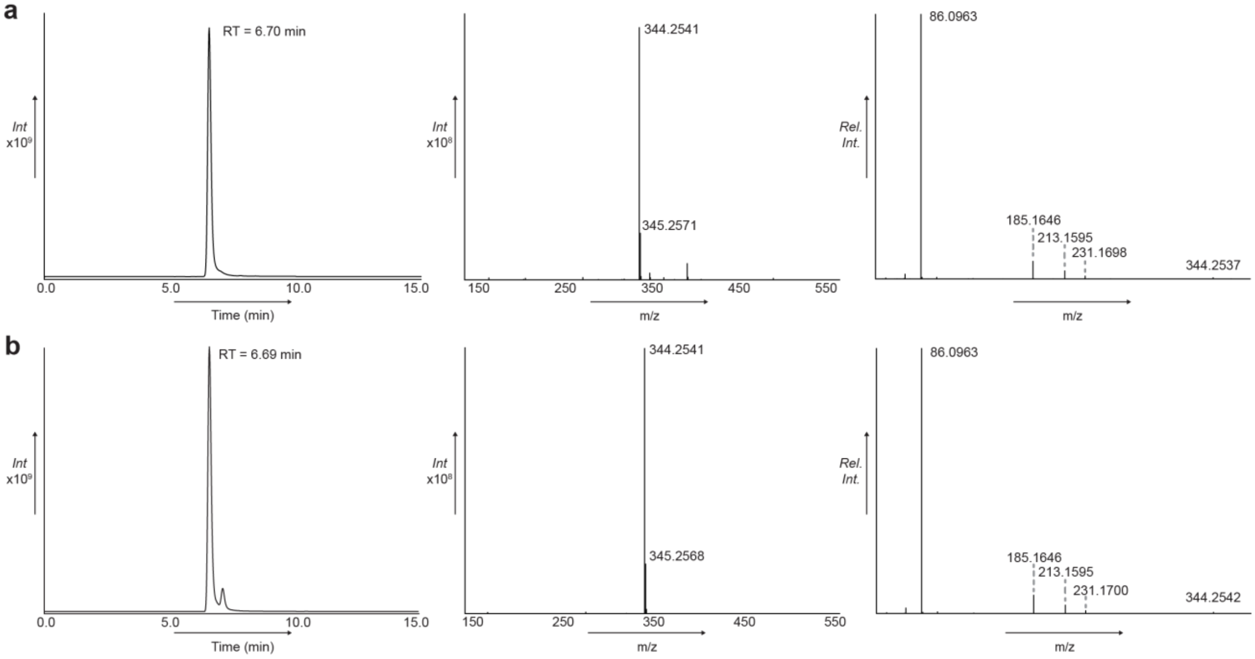


**Figure S4. Comparison of natural and synthetic gammanonin tripeptide. a.** Extracted ion chromatogram (EIC) of the natural gammanonin tripeptide (344.25 [M+H]), along with the parent ion, and MS/MS spectra. **b.** Extracted ion chromatogram (EIC) of synthetic L-N-methyl valine-L-valine-L-leucine (344.25 [M+H]), along with the parent ion, and MS/MS spectra

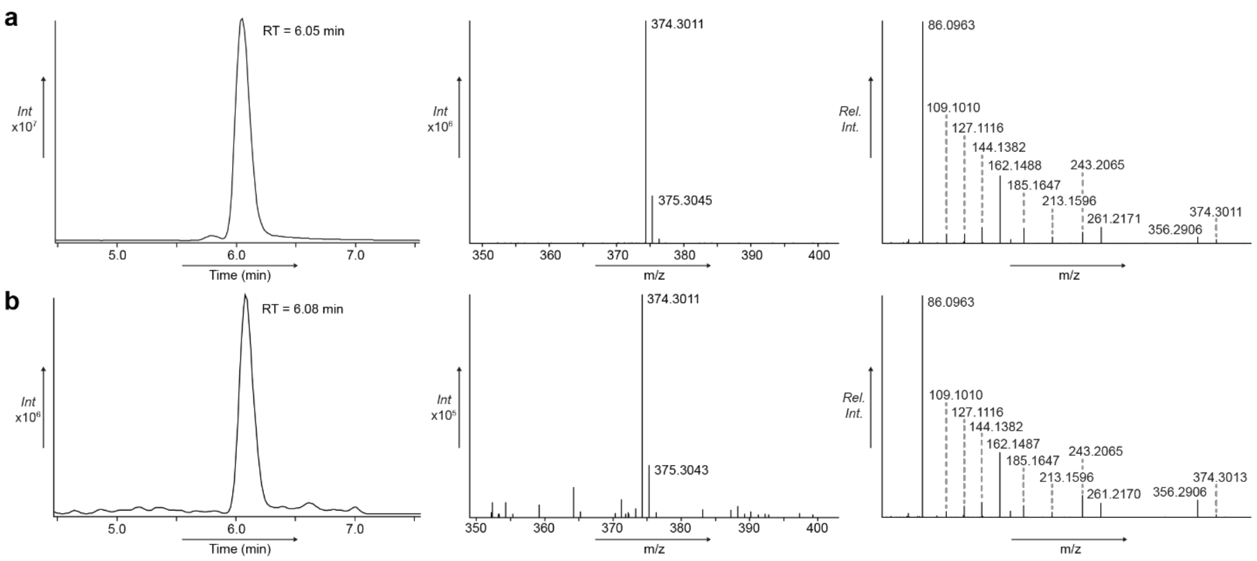


**Figure S5. Comparison of natural and synthetic gammanonin. a.** Extracted ion chromatogram (EIC) of the natural gammanonin (374.30 [M+H]), along with the parent ion, and MS/MS spectra. **b.** Extracted ion chromatogram (EIC) of synthetic gammanonin (374.30 [M+H]), along with the parent ion, and MS/MS spectra.

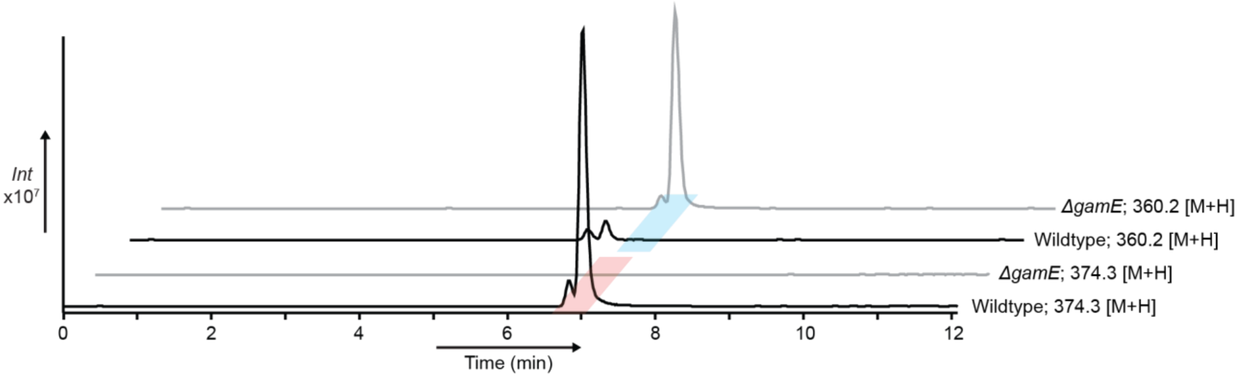


**Figure S6. Gammanonin products generated without the GamE N-methyltransferase.** Extracted cultures of *E. coli* carrying our expression plasmids with (wildtype) or without the gamE N-methyltransferase (Δ*gamE*) were analyzed by LCMS. Extracted ion chromatograms (EIC) are provided for gammanonin (374.3 [M+H]) and unmethylated gammanonin (360.2 [M+H]).

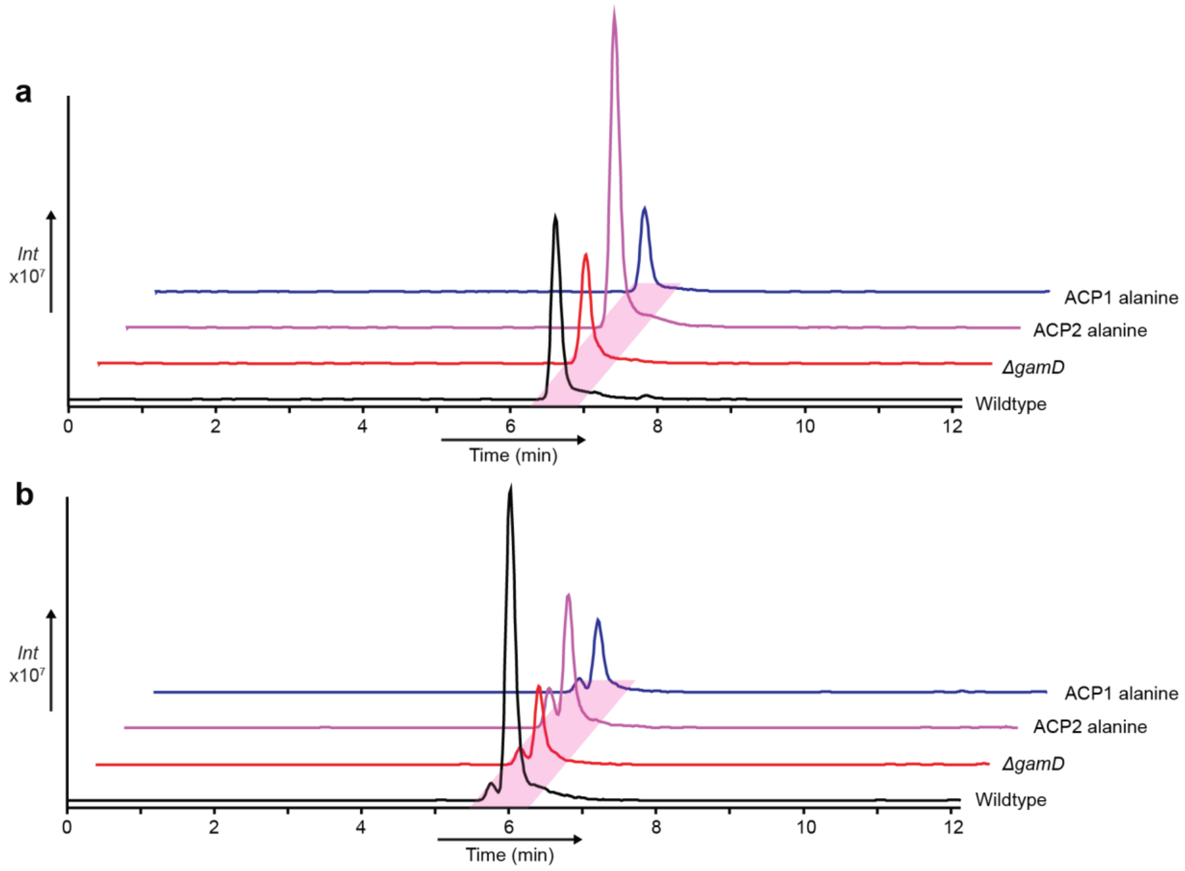


**Figure S7. Investigating the roles of *gamD* and dual PKS acyl carrier protein domains.** Shown above are LCMS traces of extracted cultures of *E. coli* carrying our expression plasmids with unmodified sequences (wildtype), without gamD (Δ*gamD*), or with the first (ACP1) or second (ACP2) PKS acyl carrier protein active serine mutated to alanine. **a.** Extracted ion chromatograms (EIC) for the gammanonin tripeptide (344.3 [M+H]). **b.** Extracted ion chromatograms (EIC) for gammanonin (374.3 [M+H]).

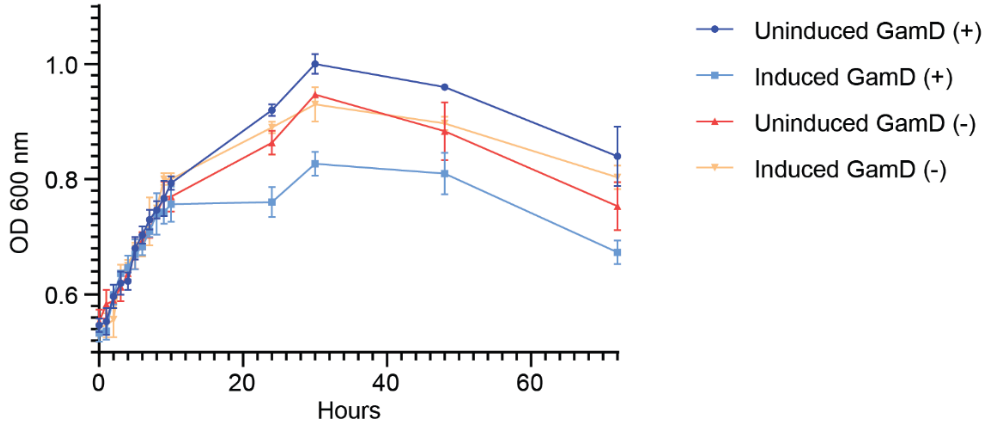


**Figure S8. Effect of GamD on *E. coli* carrying gammanonin expression plasmids.** *E. coli* BAP1 with gammanonin expression plasmids ±*gamD* were grown in M9 minimal media under production conditions, then divided and left to grow in the presence or absence of IPTG. Optical density (OD) was monitored for 72 hours. Three biological replicates were used for each condition.

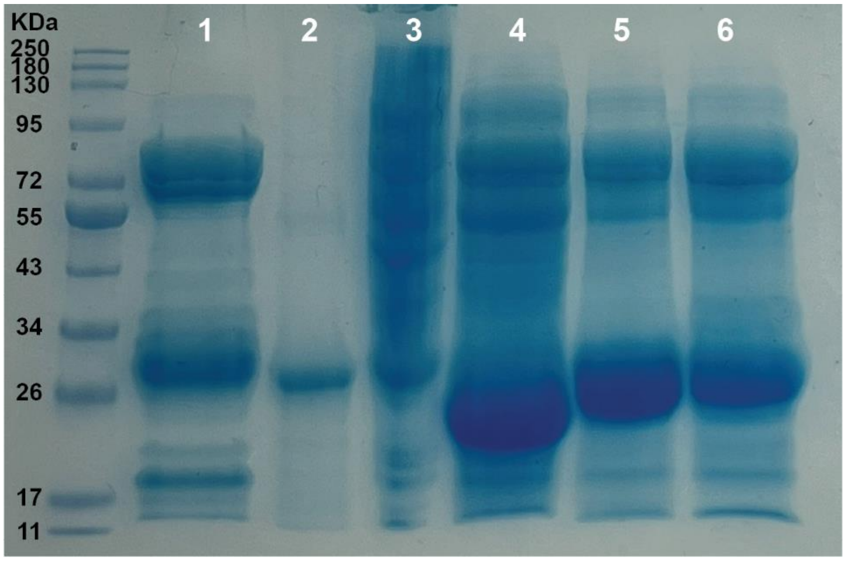


**Figure S9. Purified proteins used in this study.** SDS PAGE gel dyed with Coomassie, featuring **1.** GamE methyltransferase (27 kDa; N-terminal $His_6$ tag), **2.** *E. coli* peptide deformylase (21 kDa; N-terminal $His_6$ tag), **3.** *V. crassotrae* DSM 17220 peptide deformylase (19 kDa; N-terminal $His_6$ tag), **4.** GamD peptide deformylase (19 kDa; N-terminal $His_6$ tag), **5.** Ga0077872_1535 (*V. tubiashii* house-keeping PDF; 19 kDa; N-terminal $His_6$ tag), **6.** Ga0077872_1694 (*V. tubiashii* isolated PDF; 19 kDa; N-terminal $His_6$ tag).

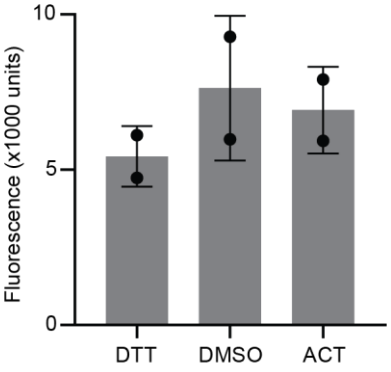


**Figure S10. The non-conserved *V. tubiashii* peptide deformylase Ga0077872_1694 has limited activity in vitro.** Peptide deformylase activity of purified Ga0077872_1694 was assessed according to our established methods, using a DTT-treated negative control, DMSO alone positive control, and 100 µM actinonin. Reactions were performed in duplicate for 30 minutes at 30°C prior to addition of fluorescamine to a final concentration of 60 µg $mL^{-1}$.

**Table S3. NMR spectroscopic data for Fmoc-leucine methyl ester (3)(600 MHz in DMSO-$d_6$)[a]**

| Position | $\delta_H$ mult. | $\delta_C$ | Position | $\delta_H$ mult. | $\delta_C$ |
|---|---|---|---|---|---|
| 1 | 7.78 (d, 8) | - | 10 | 4.32 (d, 7) | 66.0 |
| 2 | 4.06 (m) | 52.6 | 11 | 4.23 (t, 7) | 47.2 |
| 3 | - | 173.8 | 12 | - | 144.2 |
| 4 | 1.59, 1.48 (m) | 39.9 | 13 | 7.72 (d, 7.5) | 125.7 |
| 5 | 1.63 (m) | 24.6 | 14 | 7.33 (t, 7.5) | 127.5 |
| 6 | 0.85 (d, 6.5) | 21.6 | 15 | 7.42 (t, 7.5) | 128.0 |
| 7 | 0.89 (d, 6.5) | 23.3 | 16 | 7.90 (d, 7.5) | 120.5 |
| 8 | 3.62 (s) | 52.3 | 17 | - | 141.2 |
| 9 | - | 156.6 | | | |

[a]Chemical shift $\delta$ and (multiplicity, $J$ in Hz).

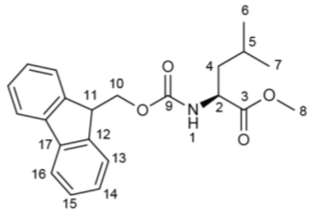


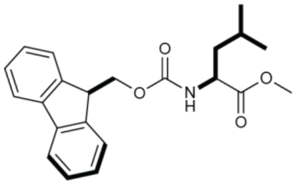


**$^1$H NMR spectrum of Fmoc-L-leucine methyl ester (3) in DMSO-$d_6$.**

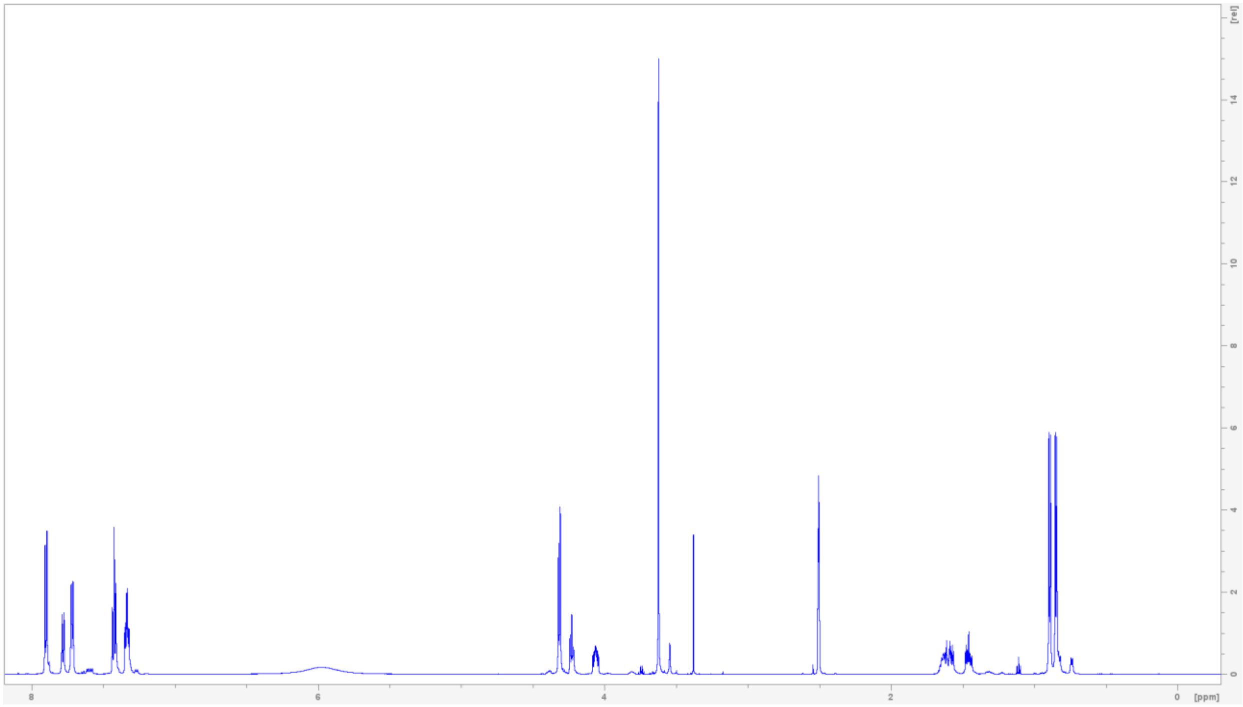

$^{13}C$ NMR spectrum of Fmoc-L-leucine methyl ester (3) in DMSO-$d_6$.

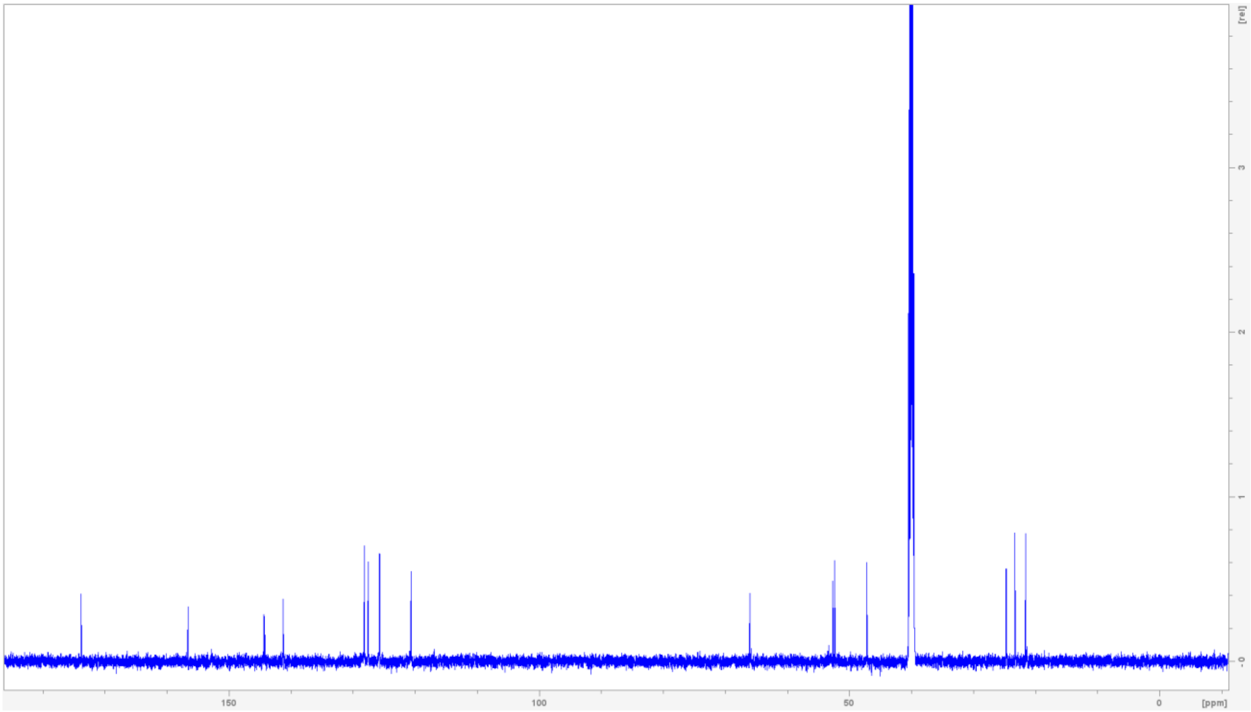


$^{1}H$-$^{1}H$ COSY NMR spectrum of Fmoc-L-leucine methyl ester (3) in DMSO-$d_6$.

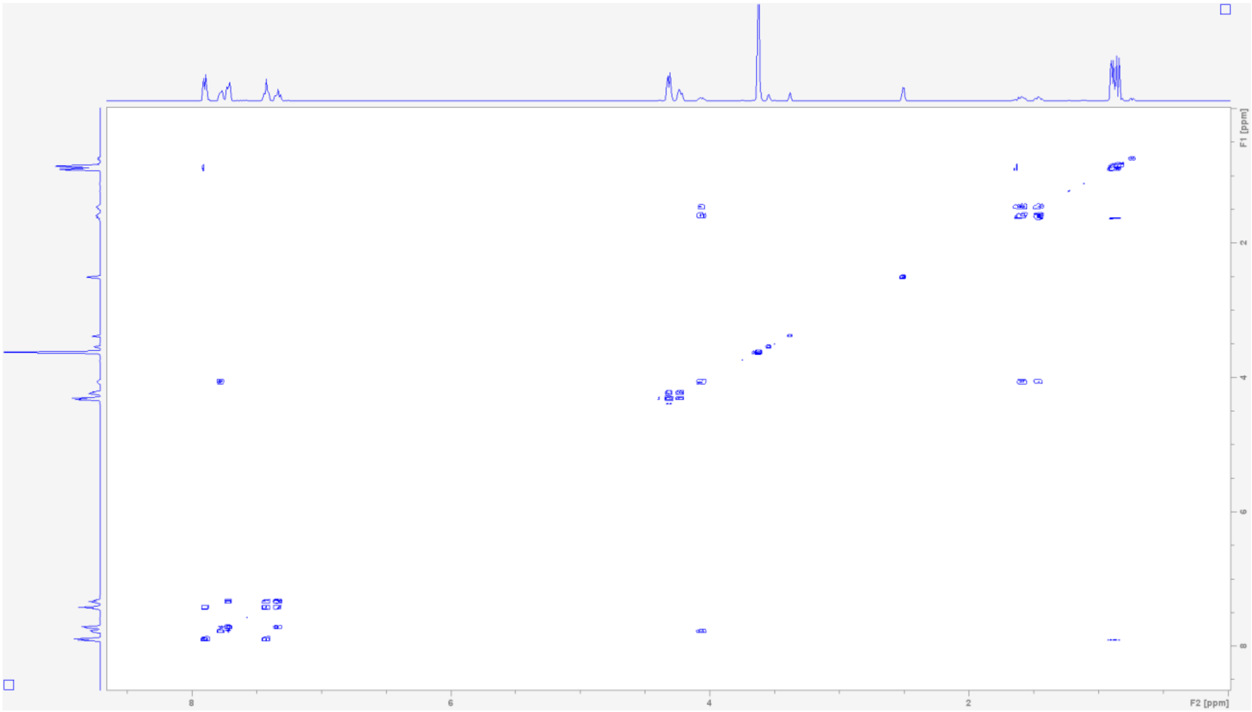

$^1H$-$^{13}C$ **HSQC NMR spectrum of Fmoc-L-leucine methyl ester (3) in DMSO-$d_6$.**

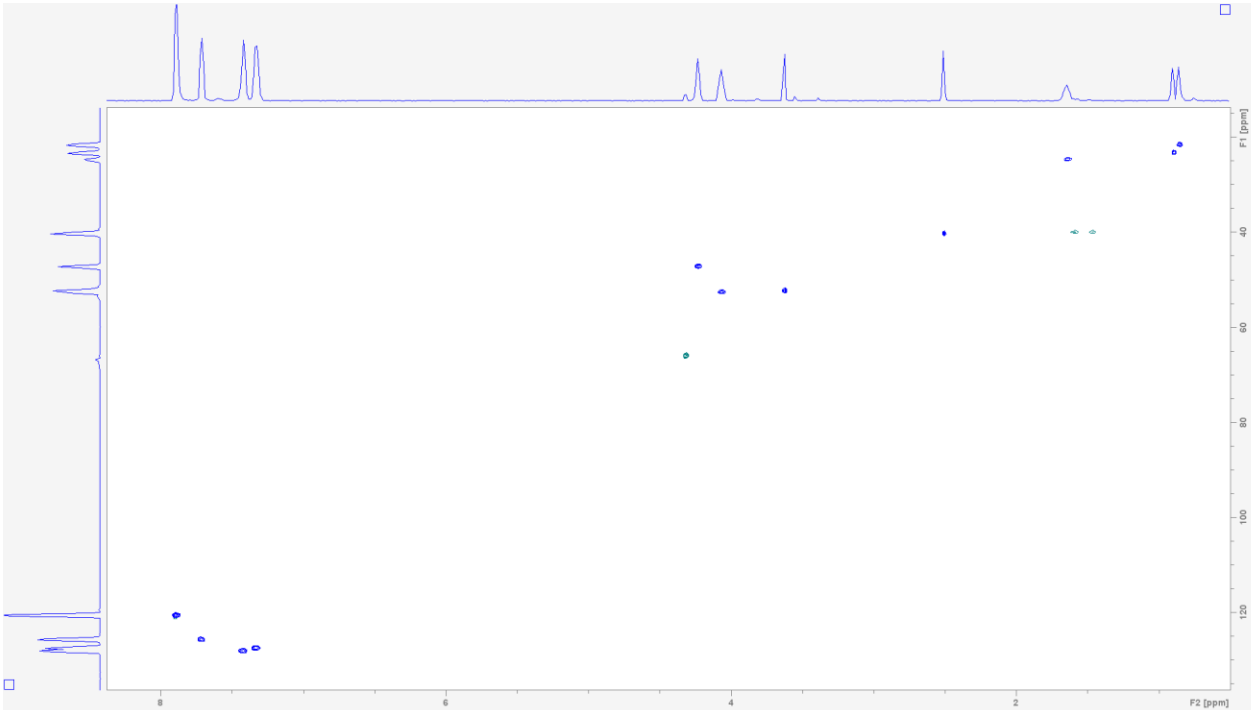


**Table S4. NMR spectroscopic data for Fmoc-leucine aldehyde (4)(600 MHz in DMSO-$d_6$)[a]**

| Position | $\delta_H$ mult. | $\delta_C$ | Position | $\delta_H$ mult. | $\delta_C$ |
|---|---|---|---|---|---|
| 1 | 7.73 (m) | - | 9 | 4.39 (d, 7) | 66.0 |
| 2 | 3.96 (m) | 58.6 | 10 | 4.23 (m) | 47.2 |
| 3 | 9.45 (s) | 202.3 | 11 | - | 144.2 |
| 4 | 1.51, 1.42 (m) | 36.8 | 12 | 7.71 (d, 7.5) | 125.6 |
| 5 | 1.57 (m) | 24.4 | 13 | 7.32 (m, 7.5) | 127.4 |
| 6 | 0.83 (m, 6.5) | 21.7 | 14 | 7.40 (m, 7.5) | 128.0 |
| 7 | 0.87 (m, 6.5) | 24.0 | 15 | 7.86 (d, 7.5) | 120.5 |
| 8 | - | 156.8 | 16 | - | 141.2 |

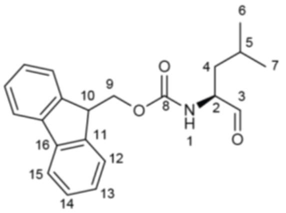


[a]Chemical shift $\delta$ and (multiplicity, $J$ in Hz).

$^{1}H$ NMR spectrum of Fmoc-leucine aldehyde (4) in DMSO-$d_6$.

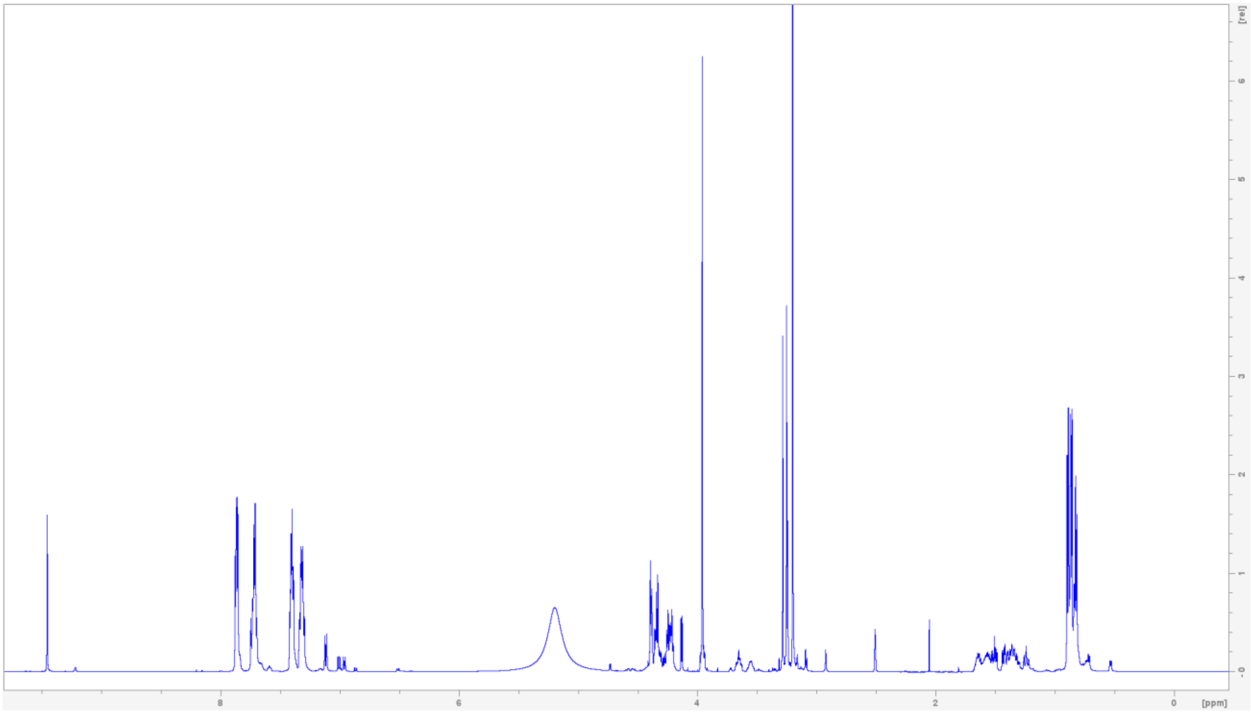


$^{13}C$ NMR spectrum of Fmoc-leucine aldehyde (4) in DMSO-$d_6$.

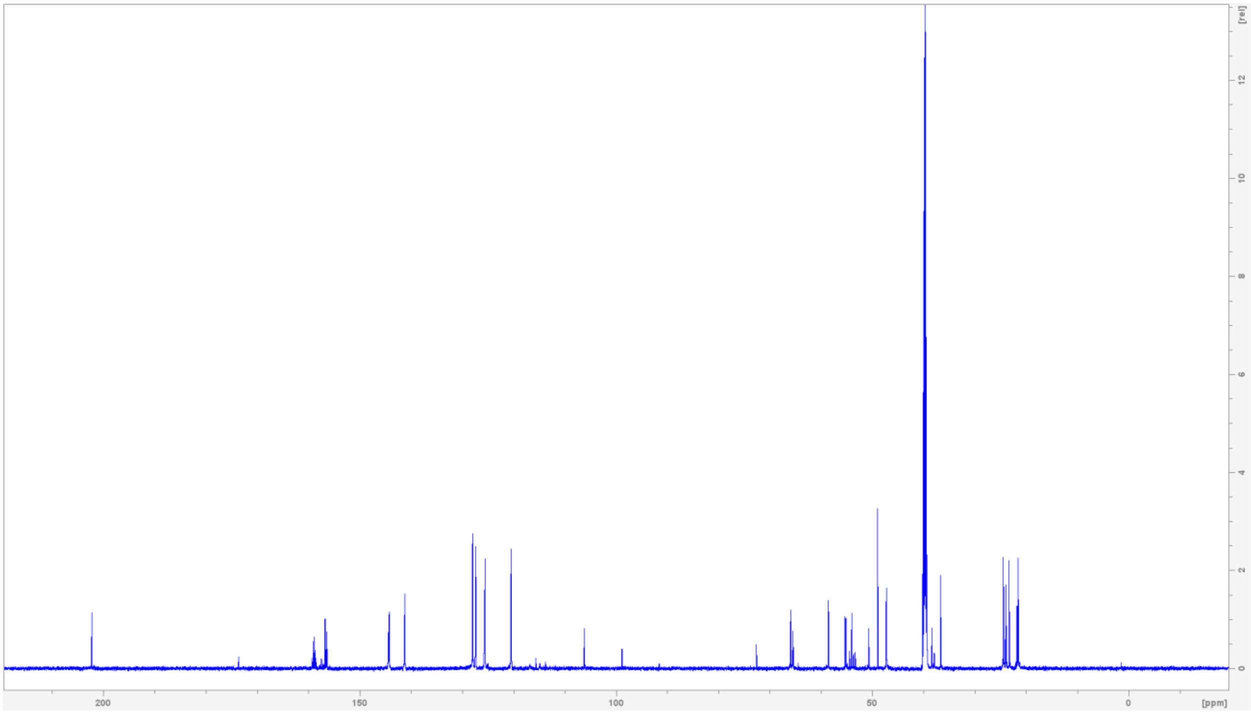

**$^{1}H$-$^{13}C$ HSQC NMR spectrum of Fmoc-leucine aldehyde (4) in DMSO-$d_6$.**

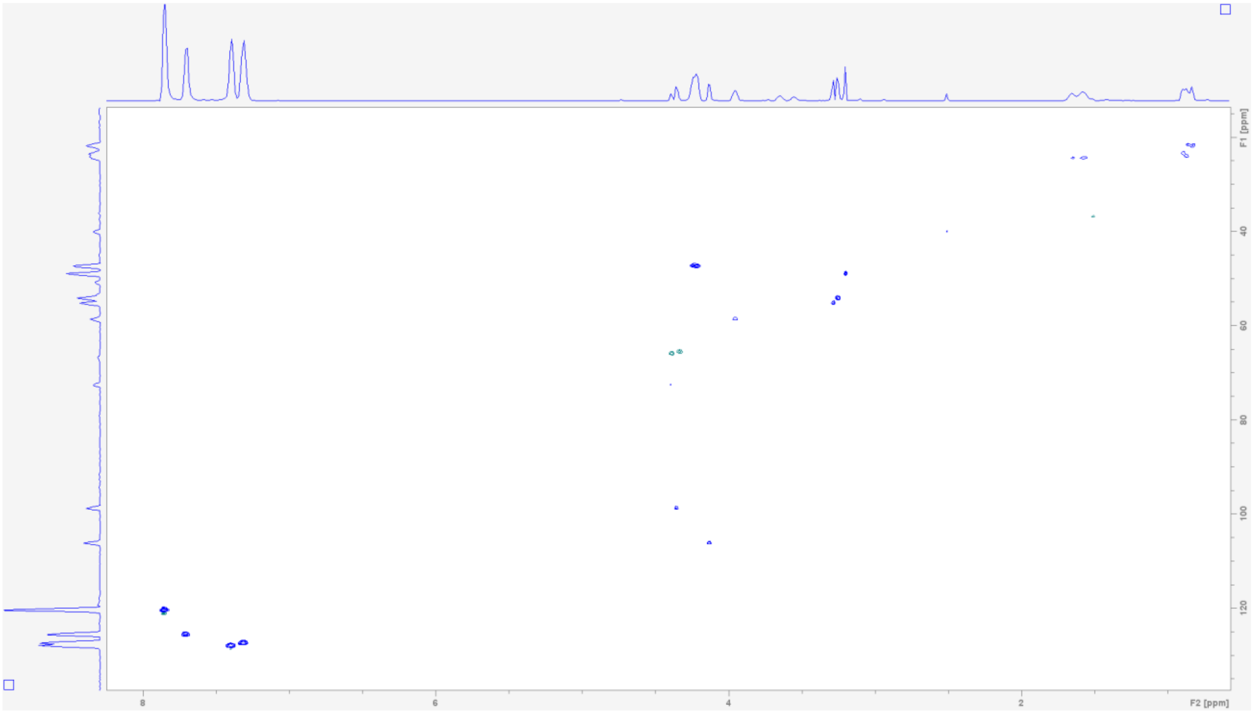


**Table S5. NMR spectroscopic data for Fmoc-(*S*)-statine ethyl ester (5)(600 MHz in DMSO-$d_6$)[a]**

| Position | $\delta_H$ mult. | $\delta_C$ | Position | $\delta_H$ mult. | $\delta_C$ |
|---|---|---|---|---|---|
| 1 | 6.96 (d, 9) | - | 12 | - | 156.6 |
| 2 | 3.57 (m) | 53.1 | 13 | 4.31 (d, 7) | 65.6 |
| 3 | 3.88 (m) | 69.5 | 14 | 4.22 (dd, 7) | 47.3 |
| 4 | 2.41, 2.20 (dd) | 39.0 | 15 | - | 144.3 |
| 5 | - | 172.0 | 16 | 7.72 (d, 7.5) | 125.7 |
| 6 | 1.36, 1.26 (b) | 38.7 | 17 | 7.32 (t, 7.5) | 127.5 |
| 7 | 1.55 (m) | 24.8 | 18 | 7.41 (t, 7.5) | 128.0 |
| 8 | 0.88 (d, 6.5) | 23.8 | 19 | 7.89 (d, 7.5) | 120.5 |
| 9 | 0.84 (d, 6.5) | 22.2 | 20 | - | 141.2 |
| 10 | 4.04 (q, 7) | 60.1 | 21 | 4.90 (d, 6) | - |
| 11 | 1.17 (t, 7) | 14.5 | | | |

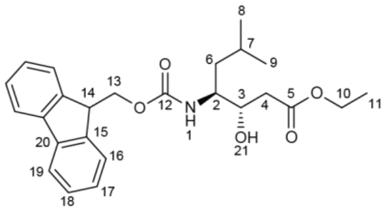


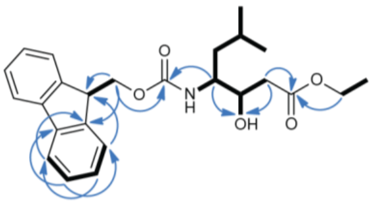


[a]Chemical shift $\delta$ and (multiplicity, $J$ in Hz).

**$^{1}$H NMR spectrum of Fmoc-(*S*)-statine ethyl ester (5) in DMSO-*d*$_6$.**

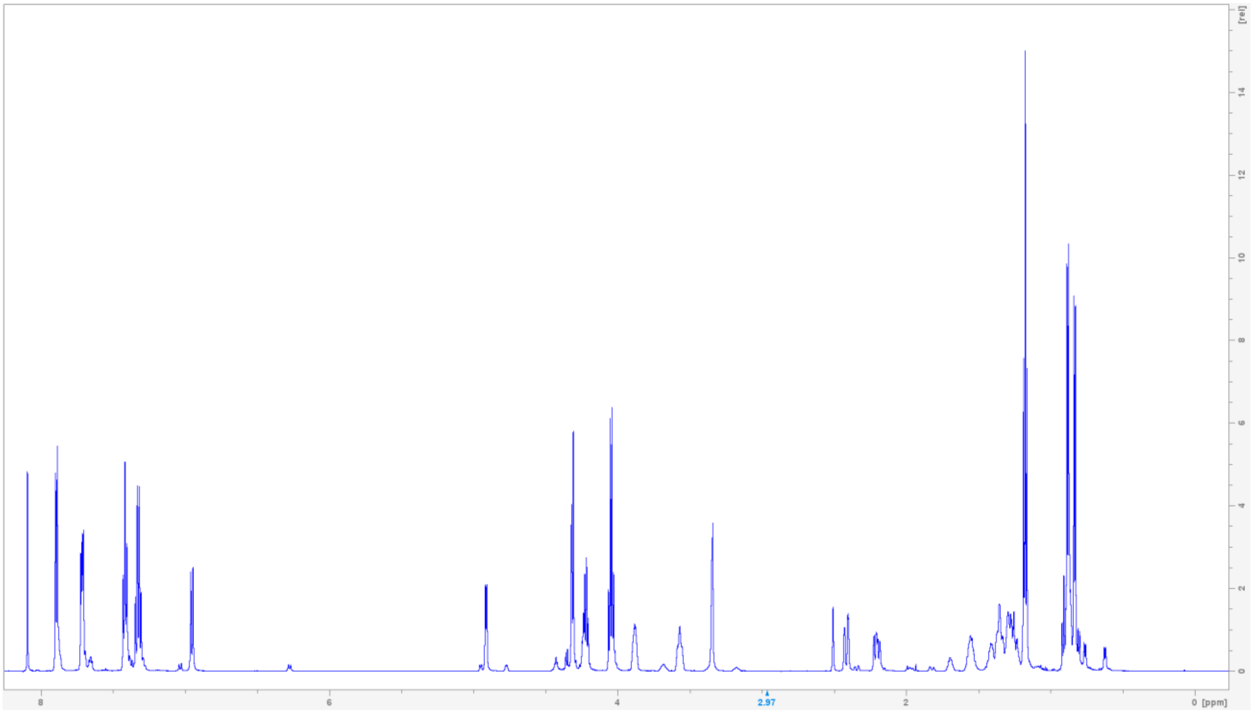

**$^{13}$C NMR spectrum of Fmoc-(*S*)-statine ethyl ester (5) in DMSO-*d*$_6$**

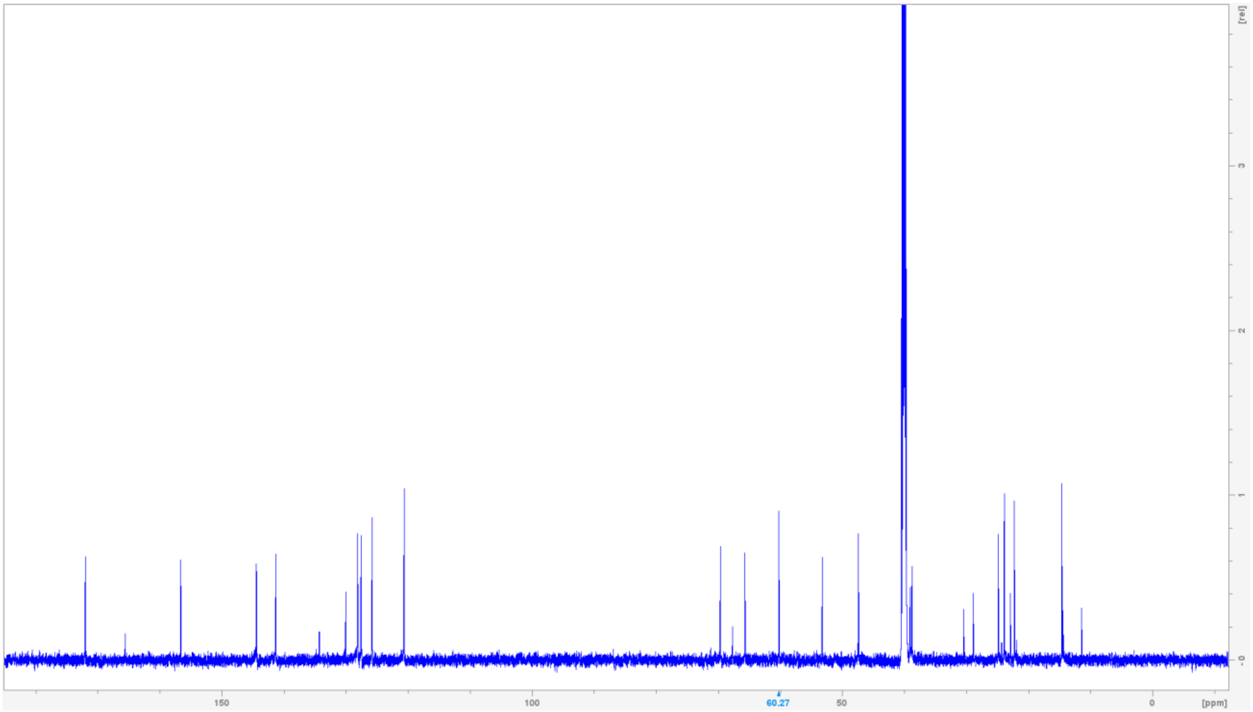

$^1H$-$^1H$ COSY NMR spectrum of Fmoc-(*S*)-statine ethyl ester (5) in DMSO-$d_6$.

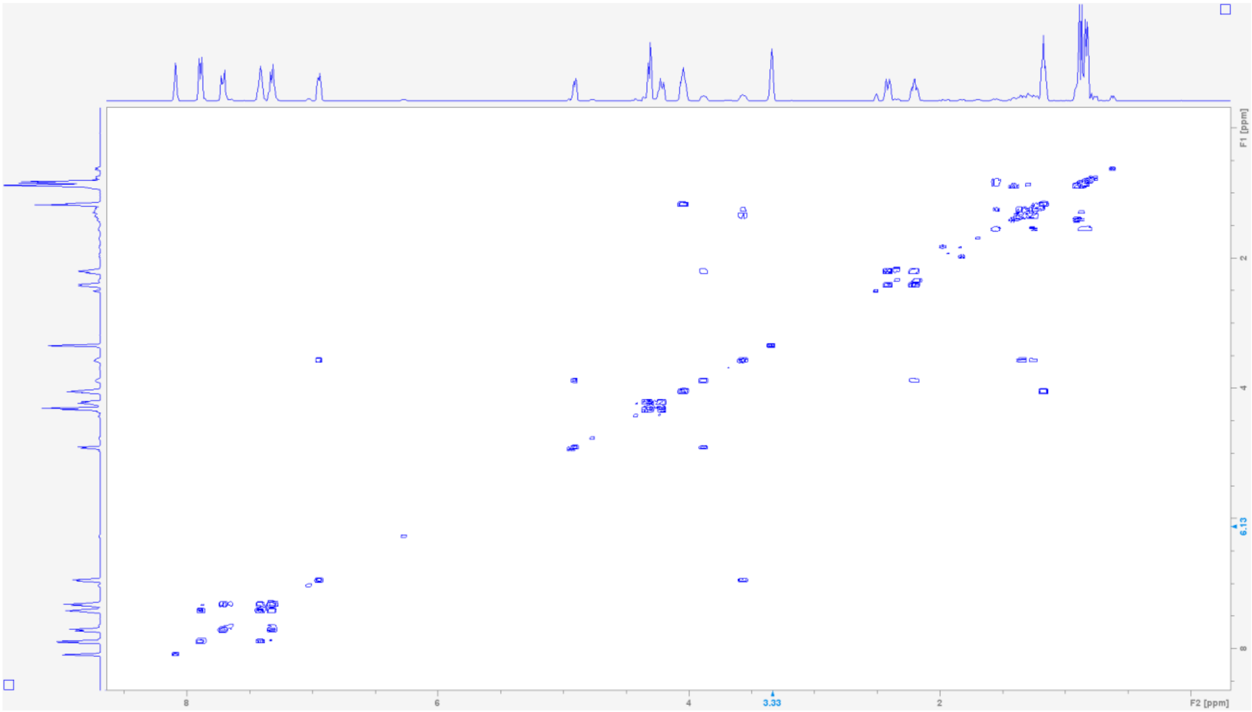

$^1H$-$^{13}C$ HSQC NMR spectrum of Fmoc-(*S*)-statine ethyl ester (5) in DMSO-$d_6$.

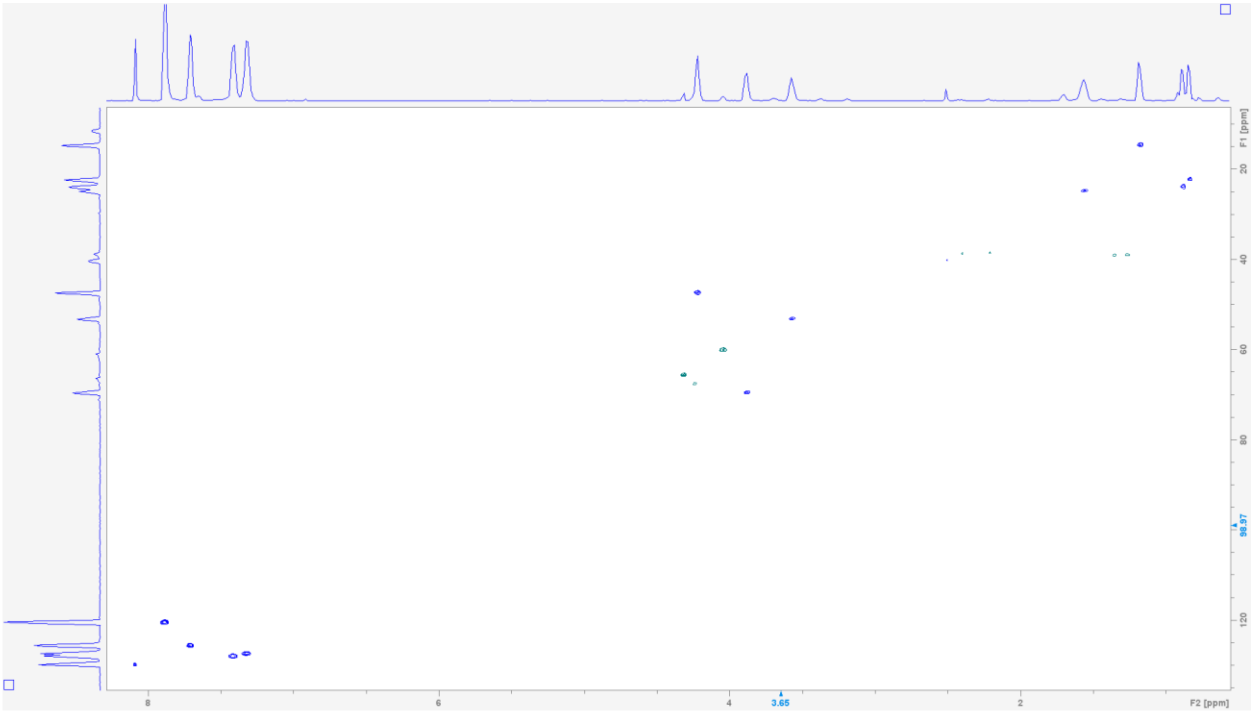

**$^{1}H$-$^{13}C$ HMBC NMR spectrum of Fmoc-(*S*)-statine ethyl ester (5) in DMSO-*d*$_6$.**

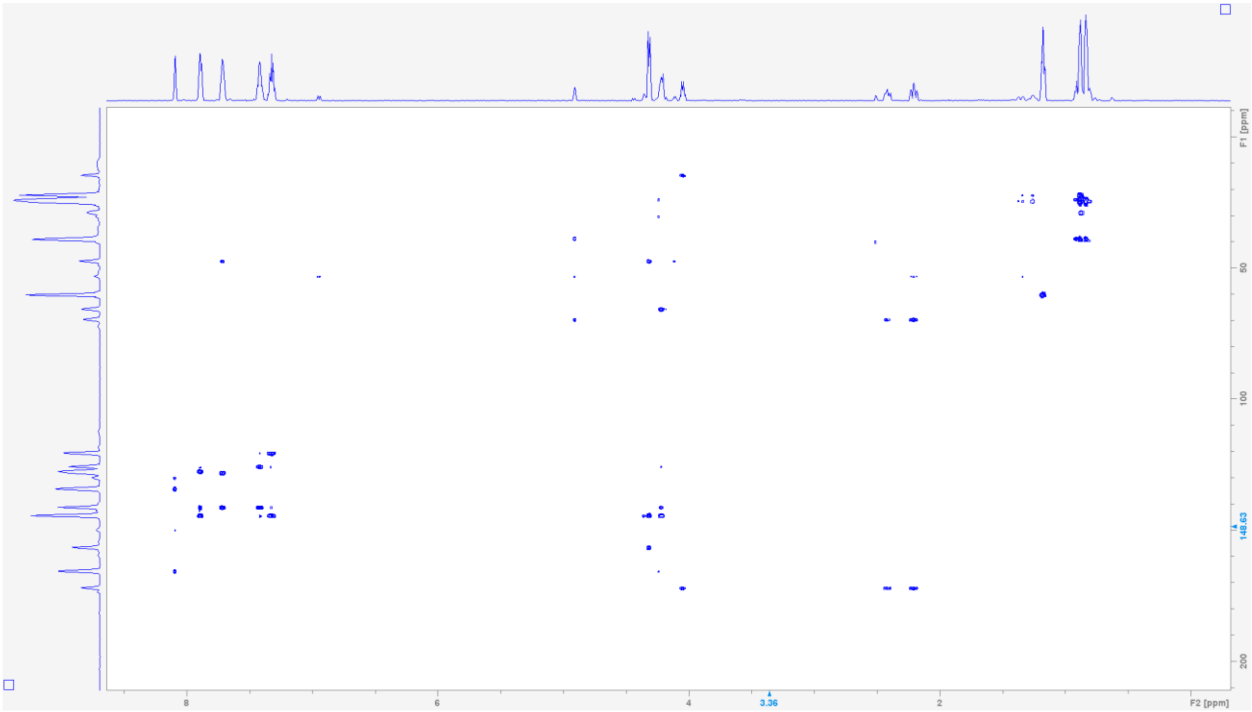


**Table S6. NMR spectroscopic data for Fmoc-L-(*S*)-statinol (7) (600 MHz in DMSO-*d*$_6$)[a]**

| Position | $\delta_H$ mult. | $\delta_C$ | Position | $\delta_H$ mult. | $\delta_C$ |
|---|---|---|---|---|---|
| 1 | 6.83 (d, 9) | - | 10 | - | 156.6 |
| 2 | 3.51 (ov) | 53.4 | 11 | 4.30, 4. 29 (m, 6) | 65.5 |
| 3 | 3.53 (ov) | 69.8 | 12 | 4.22 (t, 7.5) | 47.3 |
| 4 | 1.52, 1.40 (m) | 36.6 | 13 | - | 144.3 |
| 5 | 3.51, 3.47 (ov) | 58.7 | 14 | 7.72 (d, 7.5) | 125.7 |
| 6 | 1.40, 1.23 (m) | 39.8 | 15 | 7.32 (t, 7.5) | 127.5 |
| 7 | 1.56 (ov) | 24.8 | 16 | 7.41 (t, 7.5) | 128.0 |
| 8 | 0.88 (d, 6.5) | 24.0 | 17 | 7.89 (d, 7.5) | 120.4 |
| 9 | 0.84 (d, 6.5) | 22.3 | 18 | - | 141.2 |

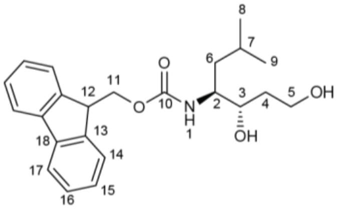


[a]Chemical shift $\delta$ and (multiplicity, *J* in Hz).

**$^1$H NMR spectrum of Fmoc-L-(*S*)-statinol (7) in DMSO-$d_6$.**

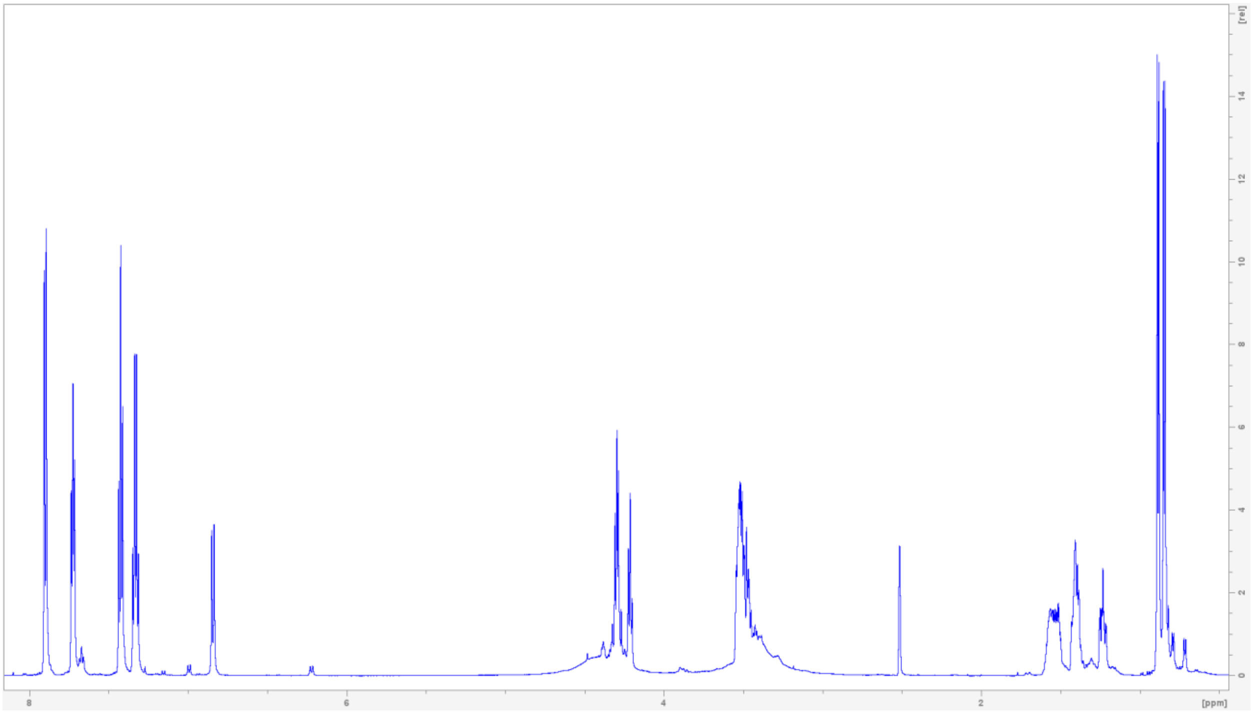

**$^{13}$C NMR spectrum of Fmoc-L-(*S*)-statinol (7) in DMSO-$d_6$.**

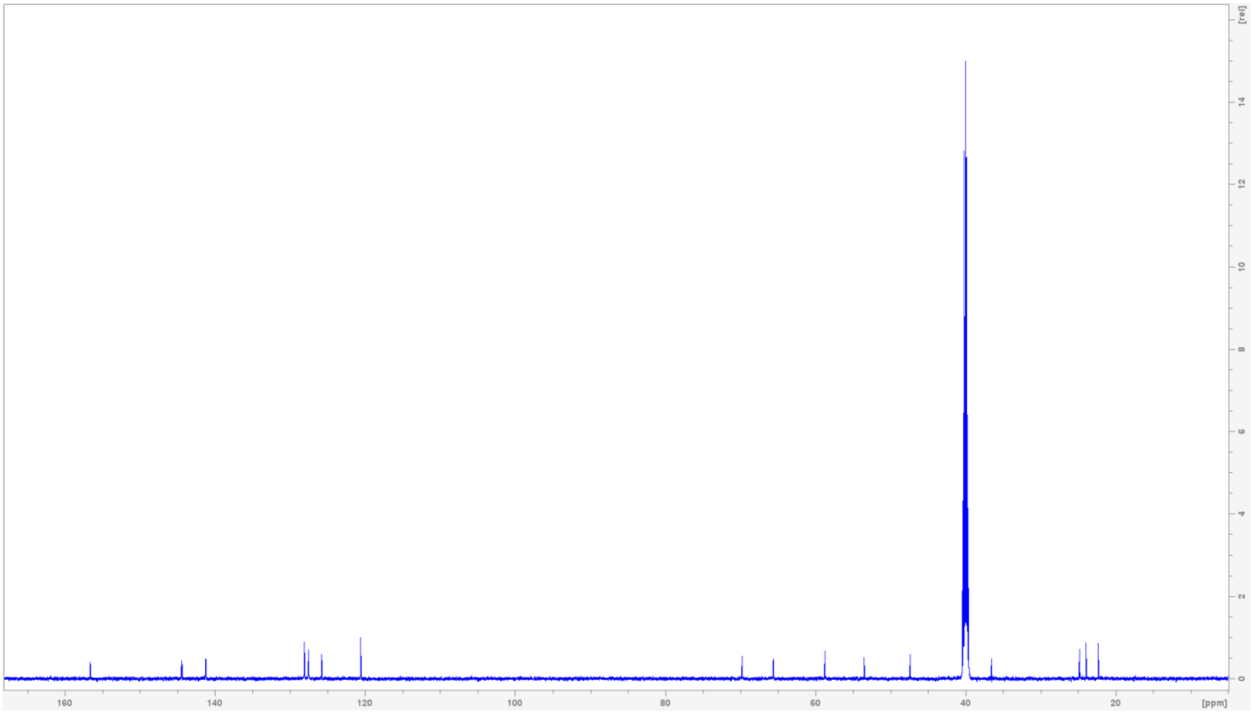

**$^1$H-$^1$H COSY NMR spectrum of Fmoc-L-(*S*)-statinol (7) in DMSO-$d_6$.**

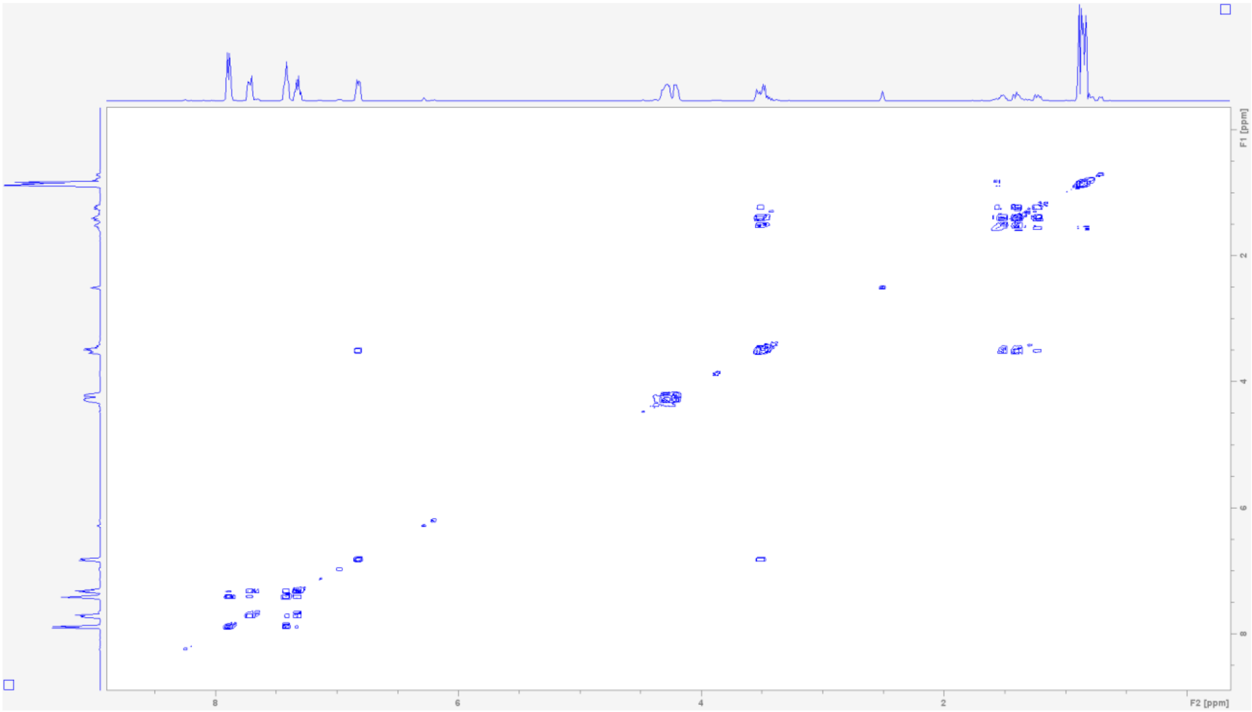


**$^1$H-$^{13}$C HSQC NMR spectrum of Fmoc-L-(*S*)-statinol (7) in DMSO-$d_6$.**

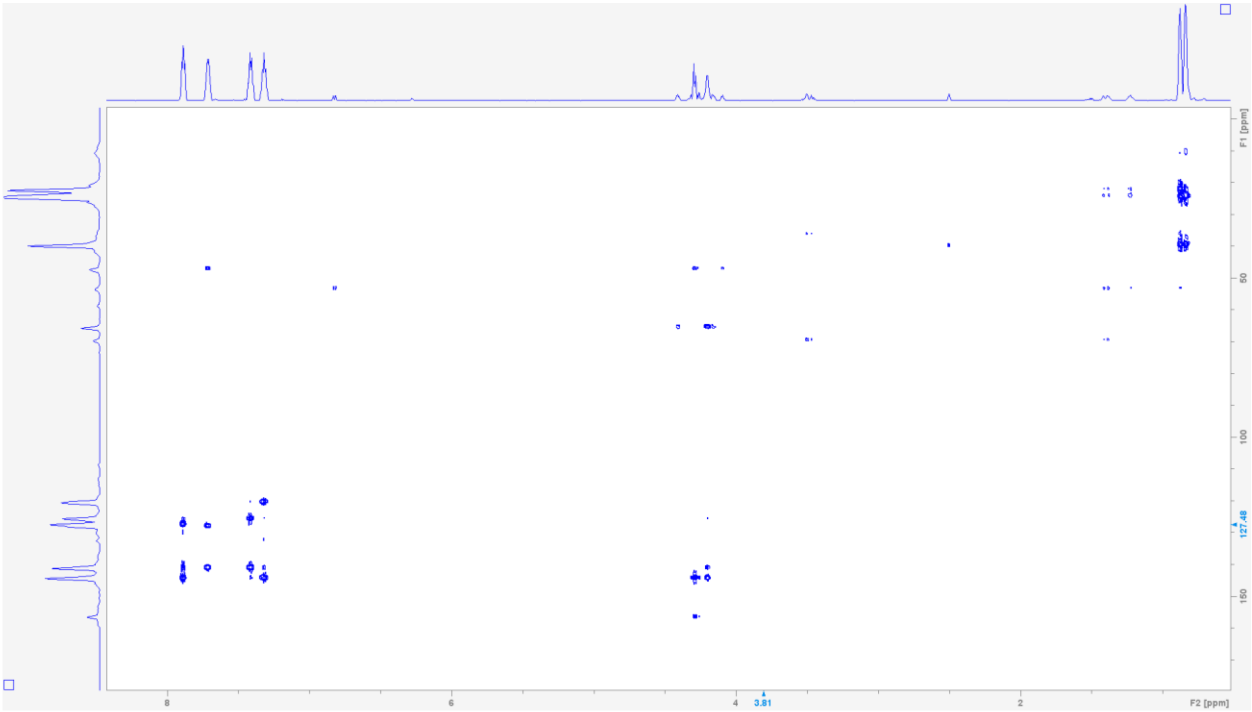

**$^{1}$H-$^{13}$C HMBC NMR spectrum of Fmoc-L-(*S*)-statinol (7) in DMSO-*d*$_6$.**

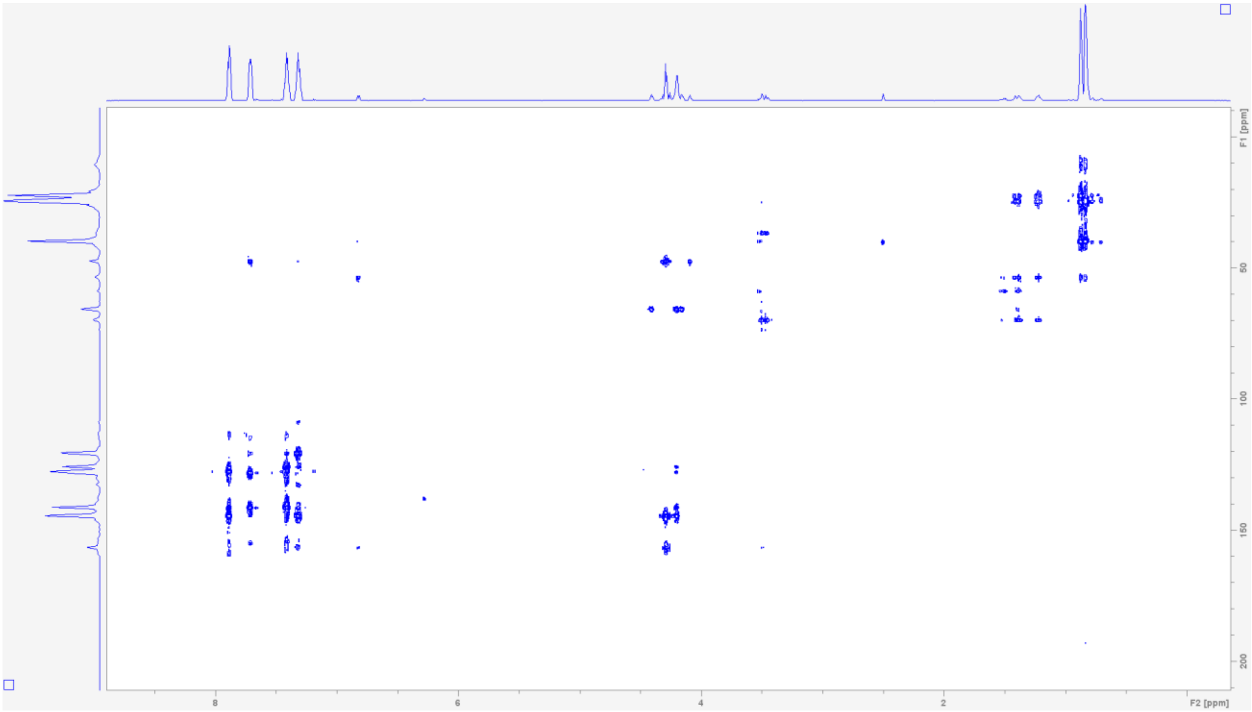


**Table S7. NMR spectroscopic data for (*S*)-statinol (8)(600 MHz in DMSO-*d*$_6$)[a]**

| Position | $\delta_H$ mult. | $\delta_C$ | Position | $\delta_H$ mult. | $\delta_C$ |
|---|---|---|---|---|---|
| 1 | - | - | 6 | 1.31, 1.27 (m, 6.5) | 41.4 |
| 2 | 2.75 (b) | 53.6 | 7 | 1.73 (m, 6.5) | 24.3 |
| 3 | 3.48 (m) | 69.2 | 8 | 0.86 (d, 6.5) | 22.4 |
| 4 | 1.61, 1.53 (m) | 37.1 | 9 | 0.89 (d, 6.5) | 23.8 |
| 5 | 3.53, 3.49 (m) | 58.3 | | | |

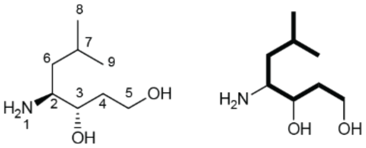


[a]Chemical shift $\delta$ and (multiplicity, *J* in Hz).

**$^1$H NMR spectrum of (*S*)-statinol (8) in DMSO-$d_6$.**

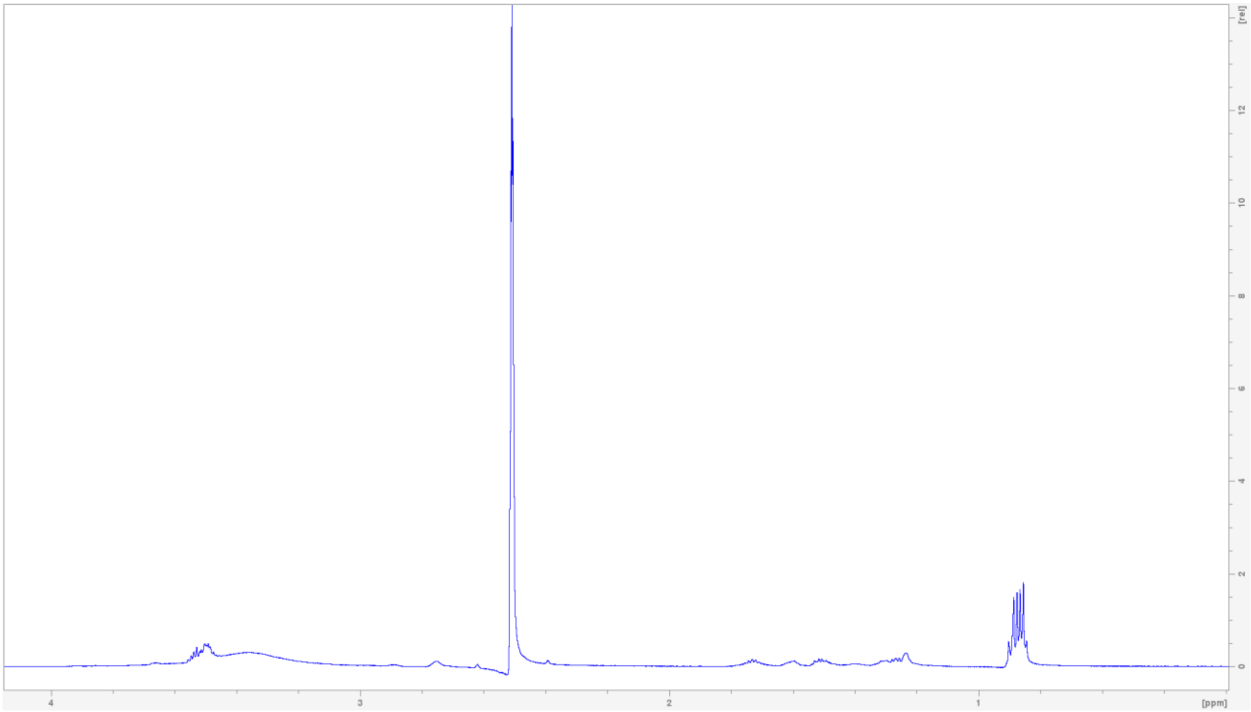



**$^1$H-$^1$H COSY NMR spectrum of (*S*)-statinol (8) in DMSO-$d_6$.**

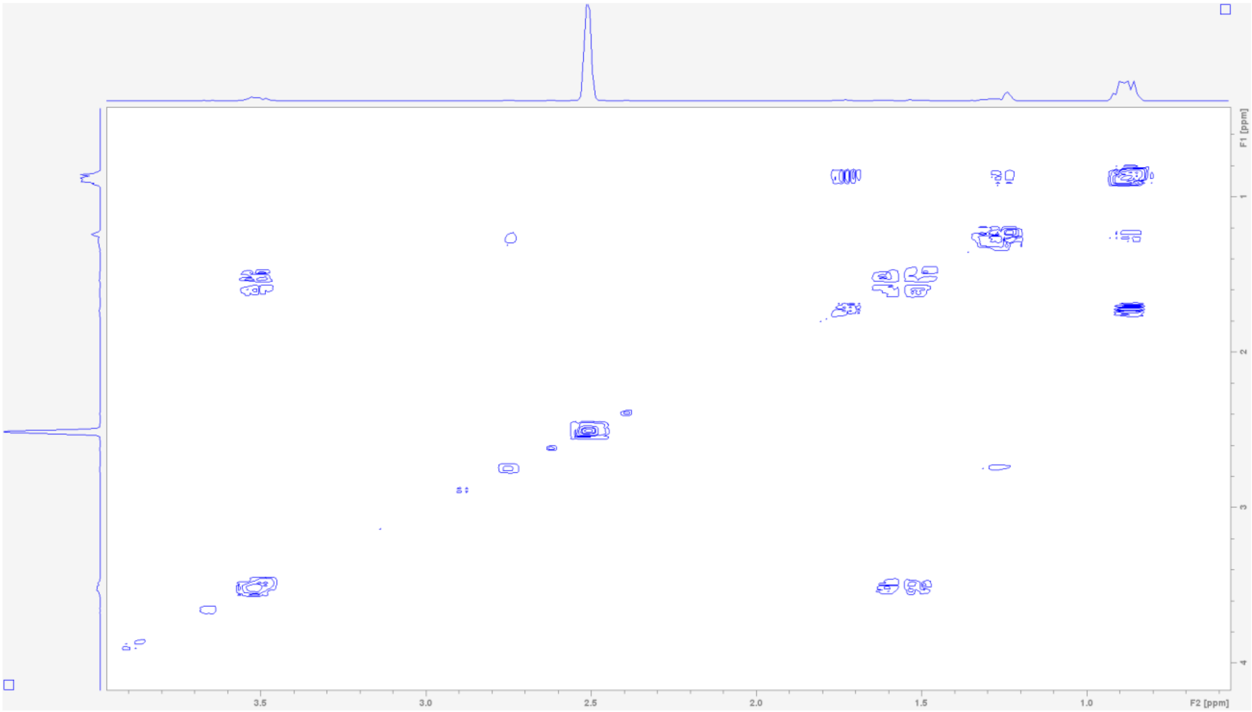

**$^1$H-$^{13}$C HSQC NMR spectrum of (*S*)-statinol (8) in DMSO-*d*$_6$.**

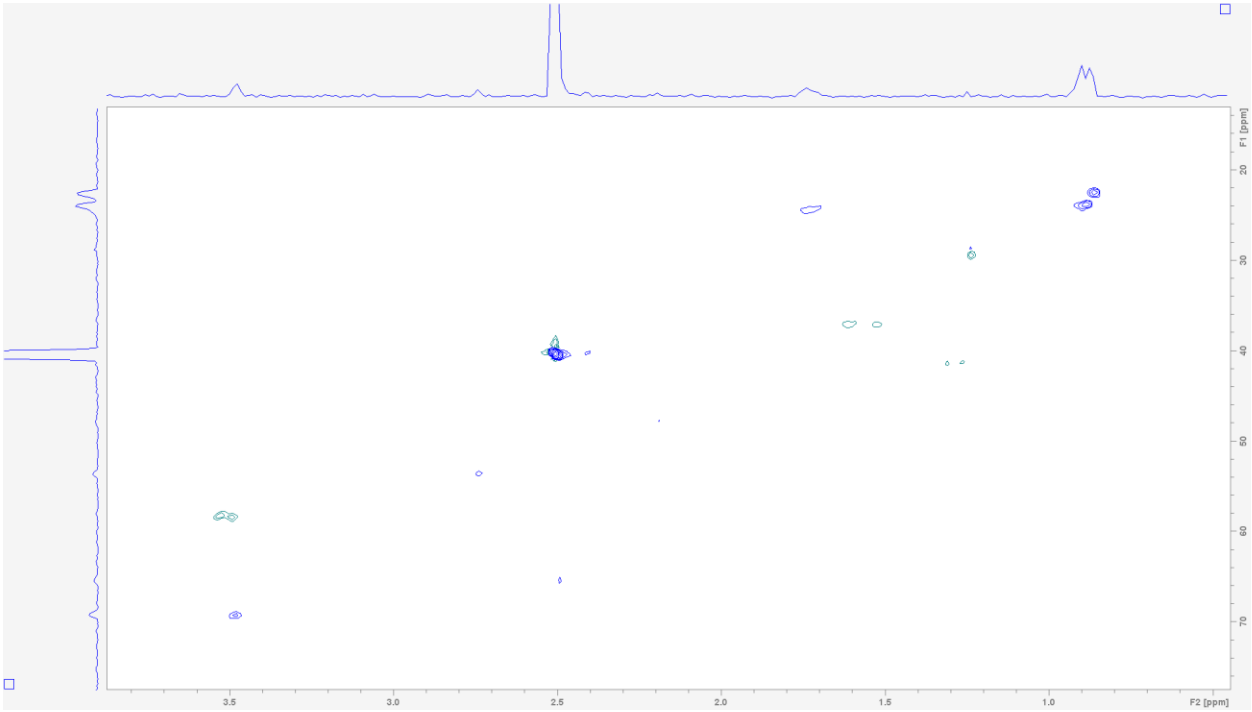


**Table S8. NMR spectroscopic data for Fmoc-L-*N*-methyl-valine-L-valine (9)(600 MHz in DMSO-*d*$_6$)[a]**

| Position | $\delta_H$ mult. | $\delta_C$ | Position | $\delta_H$ mult. | $\delta_C$ |
|---|---|---|---|---|---|
| 1 | 2.81 (s) | 29.9 | 12 | 1.03 (d, 7) | 18.0 |
| 2 | 4.31 (ov) | 63.6 | 13 | - | 156.5 |
| 3 | - | 170.8 | 14 | 4.42, 4.36 (ov) | 67.2 |
| 4 | 2.08 (ov) | 27.2 | 15 | 4.29 (ov) | 47.2 |
| 5 | 0.76 (ov) | 19.2 | 16 | - | 144.2 |
| 6 | 0.89 (ov) | 19.5 | 17 | 7.62 (d, 7.5) | 125.4 |
| 7 | 8.05 (d, 8) | - | 18 | 7.31 (t, 7.5) | 127.5 |
| 8 | 4.07 (t, 7) | 57.7 | 19 | 7.41 (t, 7.5) | 128.0 |
| 9 | - | 173.1 | 20 | 7.88 (d, 7.5) | 120.5 |
| 10 | 2.05 (m, 7) | 29.8 | 21 | - | 141.2 |
| 11 | 0.99 (d, 7) | 18.2 | | | |

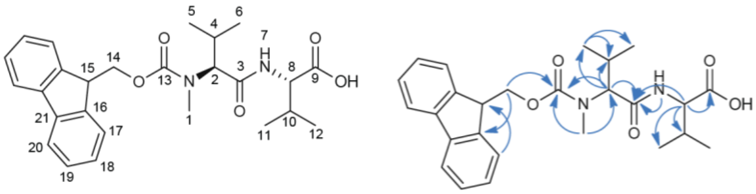

[a]Chemical shift $\delta$ and (multiplicity, *J* in Hz).

[1]H NMR spectrum of Fmoc-L-*N*-methyl-valine-L-valine (9) in DMSO-$d_6$.

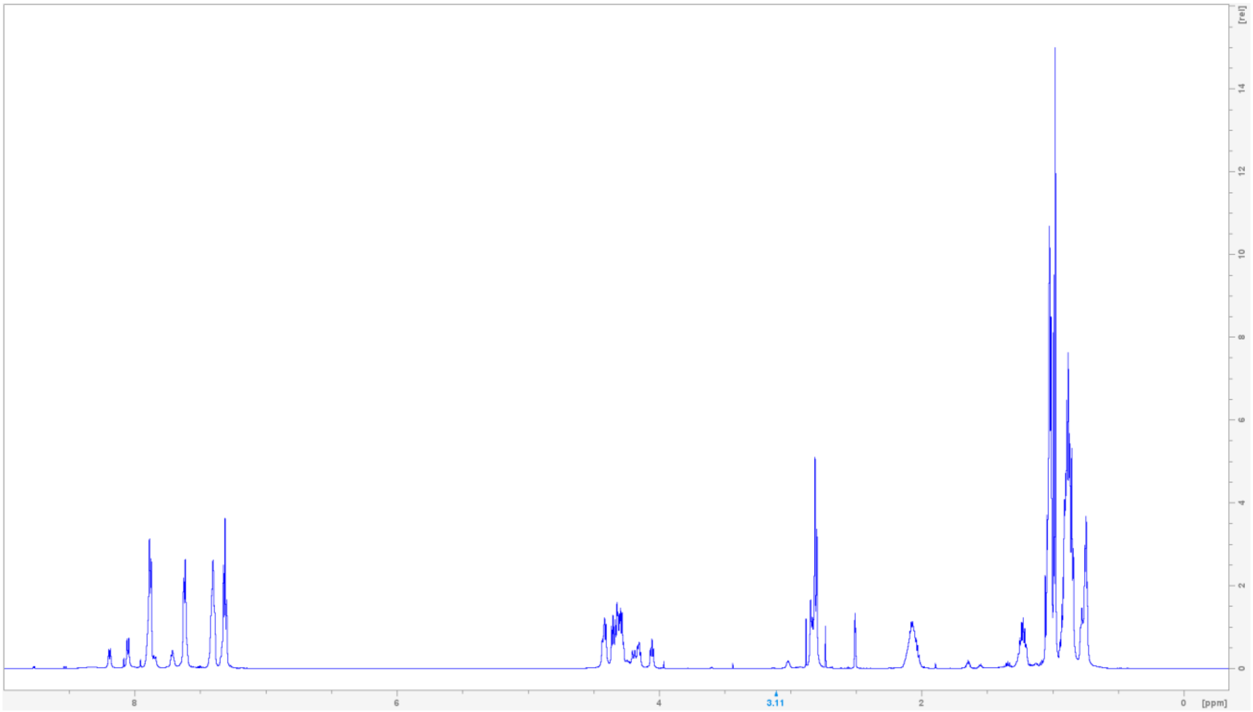

[13]C NMR spectrum of Fmoc-L-*N*-methyl-valine-L-valine (9) in DMSO-$d_6$.

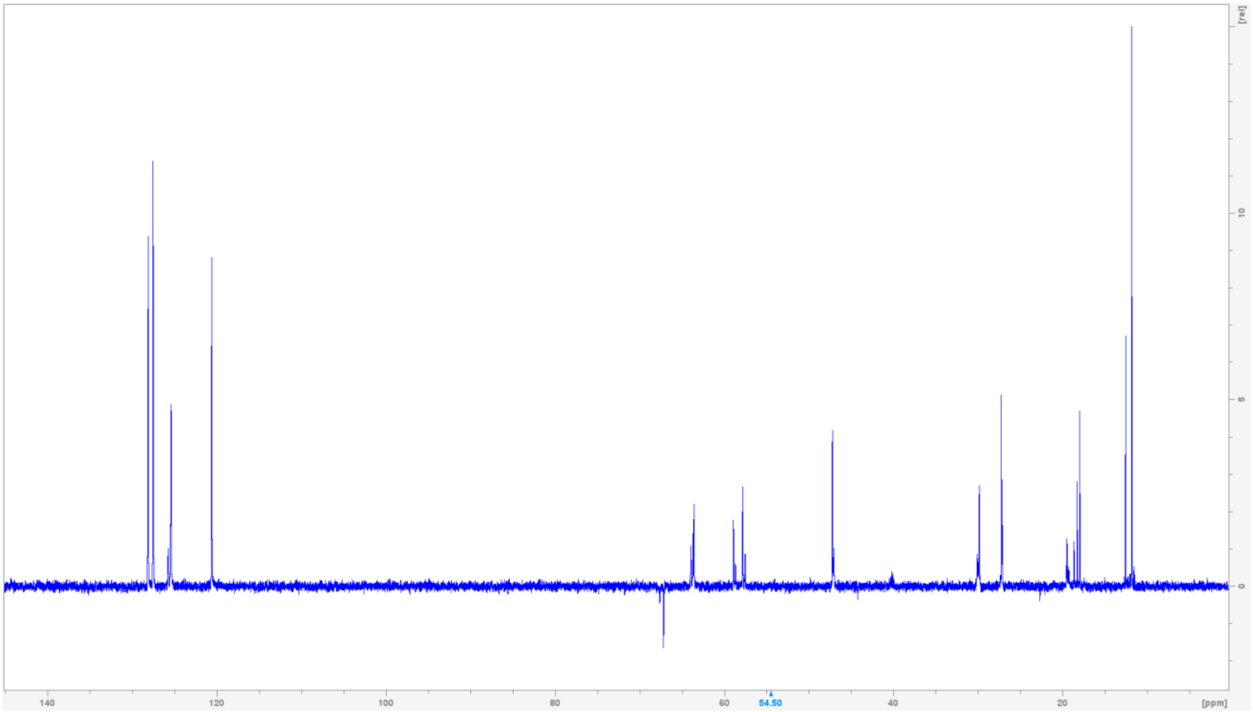

$^1$H-$^{13}$C HSQC NMR spectrum of Fmoc-L-*N*-methyl-valine-L-valine (9) in DMSO-$d_6$.

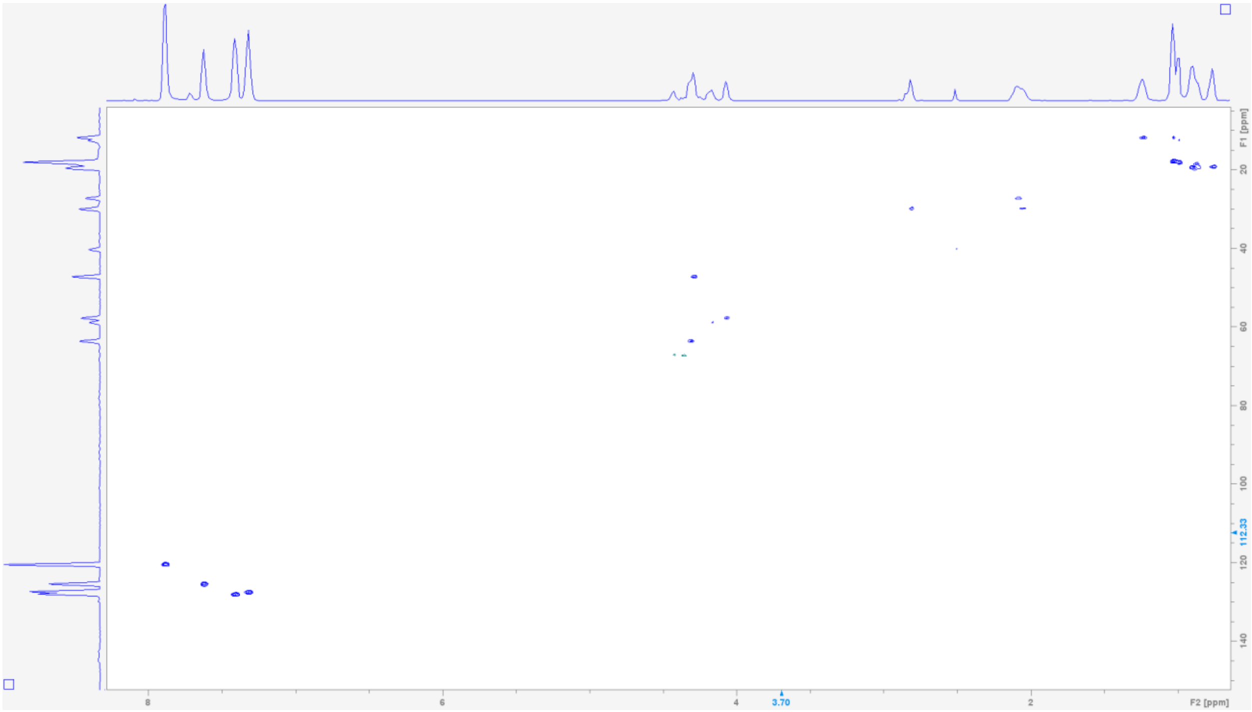


$^1$H-$^{13}$C HMBC NMR spectrum of Fmoc-L-*N*-methyl-valine-L-valine (9) in DMSO-$d_6$.

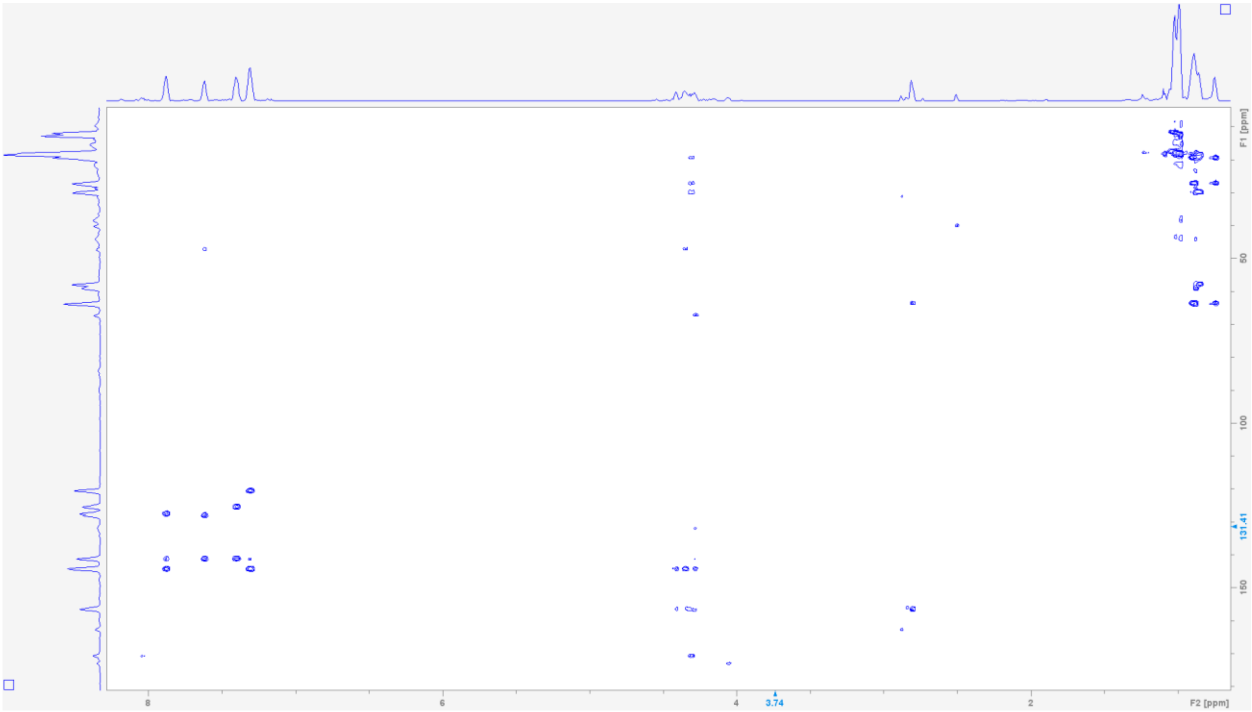

**Table S9. NMR spectroscopic data for (S) MTPA ester of Fmoc-(S)-statine ethyl ester (10) (600 MHz in DMSO-$d_6$)[a]**

| Position | $\delta_H$ mult. | Position | $\delta_H$ mult. |
|---|---|---|---|
| 1 | 7.28 (m) | 15 | - |
| 2 | 3.79 (br) | 16 | 7.68 (d, 7.5) |
| 3 | 5.40 (m) | 17 | 7.32 (t, 7.5) |
| 4 | 2.81, 2.56 (br) | 18 | 7.42 (t, 7.5) |
| 5 | - | 19 | 7.90 (d, 7.5) |
| 6 | 1.27 (ov) | 20 | - |
| 7 | - | 21 | - |
| 8 | 0.77 (d, 7) | 22 | - |
| 9 | 0.67 (d, 7) | 23 | - |
| 10 | 4.07 (q, 7) | 24 | - |
| 11 | 1.17 (t, 7) | 25 | - |
| 12 | - | 26 | - |
| 13 | 4.41, 4.33 (m) | 27 | - |
| 14 | 4.22 (m) | 28 | - |

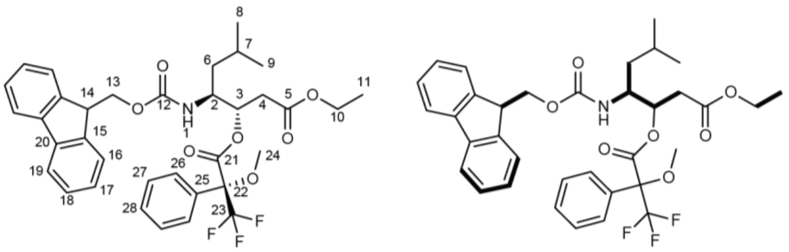

[a]Chemical shift $\delta$ and (multiplicity, $J$ in Hz)

**$^1$H NMR spectrum of (S) MTPA ester of Fmoc-(S)-statine ethyl ester (10) in DMSO-$d_6$.**

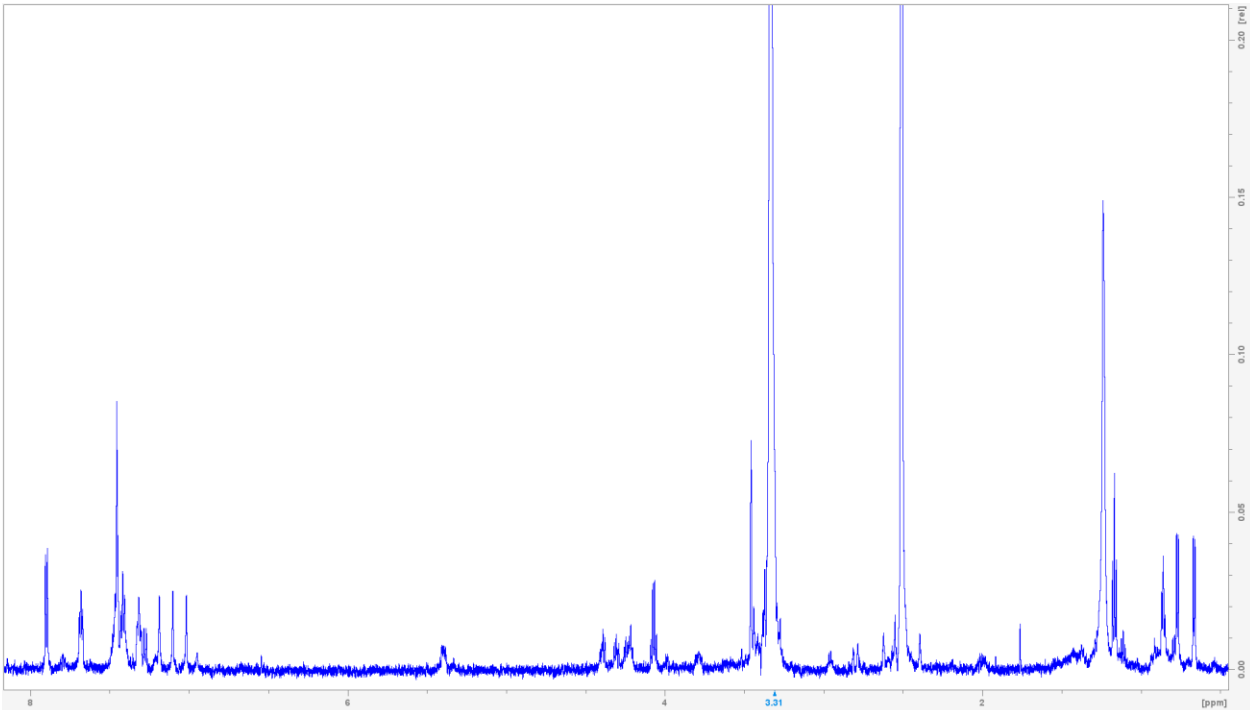

**$^1$H-$^1$H COSY NMR spectrum of (S) MTPA ester of Fmoc-(S)-statine ethyl ester (10) in DMSO-*d*$_6$.**

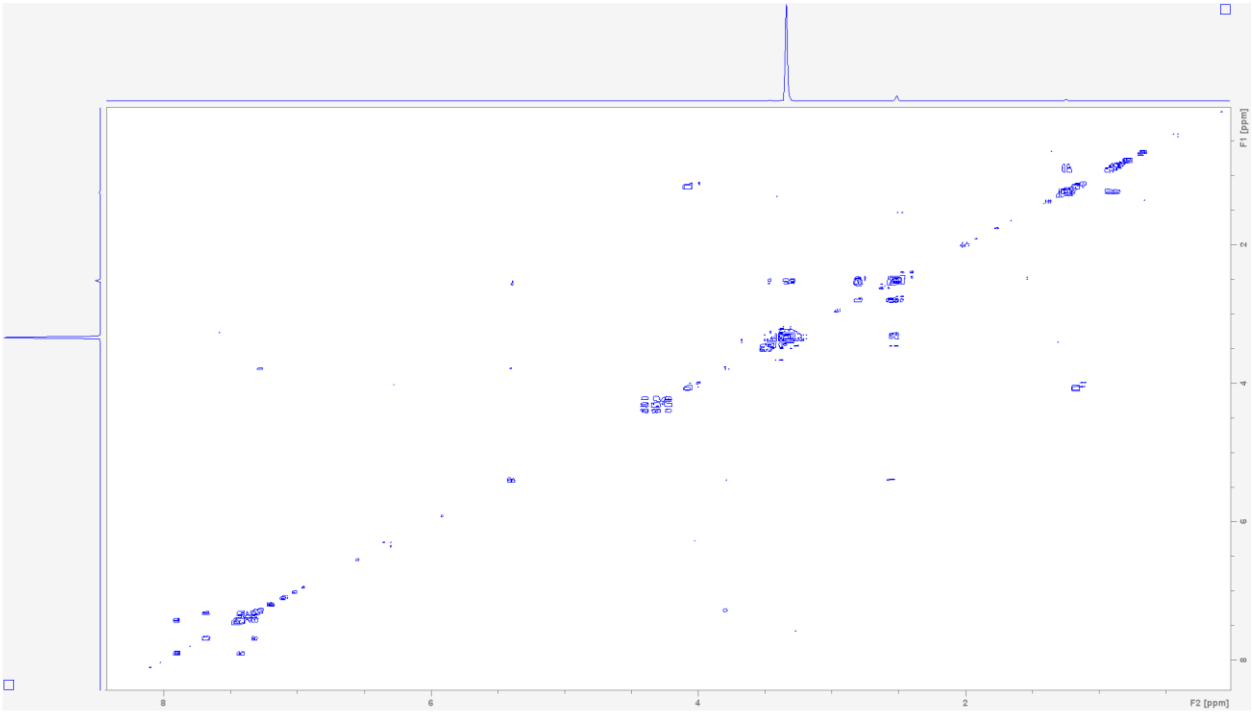

**Table S10. NMR spectroscopic data for (R) MTPA ester of Fmoc-(S)-statine ethyl ester (11) (600 MHz in DMSO-*d*$_6$)[a]**

| Position | $\delta_H$ mult. | Position | $\delta_H$ mult. |
|---|---|---|---|
| 1 | 7.36 (d, 9) | 15 | - |
| 2 | 3.87 (br) | 16 | 7.68 (d, 7.5) |
| 3 | 5.42 (m) | 17 | 7.32 (t, 7.5) |
| 4 | 2.76, 2.48 (m) | 18 | 7.42 (t, 7.5) |
| 5 | - | 19 | 7.89 (d, 7.5) |
| 6 | 1.41 (br) | 20 | - |
| 7 | 1.54 (br) | 21 | - |
| 8 | 0.87 (d, 7) | 22 | - |
| 9 | 0.80 (d, 7) | 23 | - |
| 10 | 4.00 (q, 7) | 24 | - |
| 11 | 1.12 (m) | 25 | - |
| 12 | - | 26 | - |
| 13 | 4.41, 4.33 (d, 7.5) | 27 | - |
| 14 | 4.22 (t, 7.5) | 28 | - |

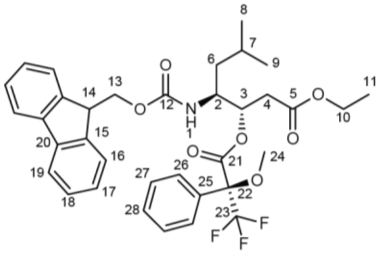

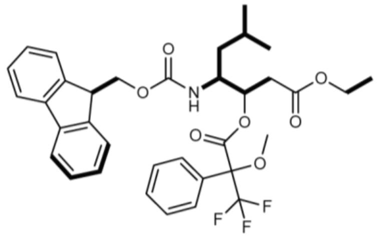

[a]Chemical shift $\delta$ and (multiplicity, *J* in Hz).

[1]H NMR spectrum of (R) MTPA ester of Fmoc-(S)-statine ethyl ester (11) in DMSO-$d_6$.

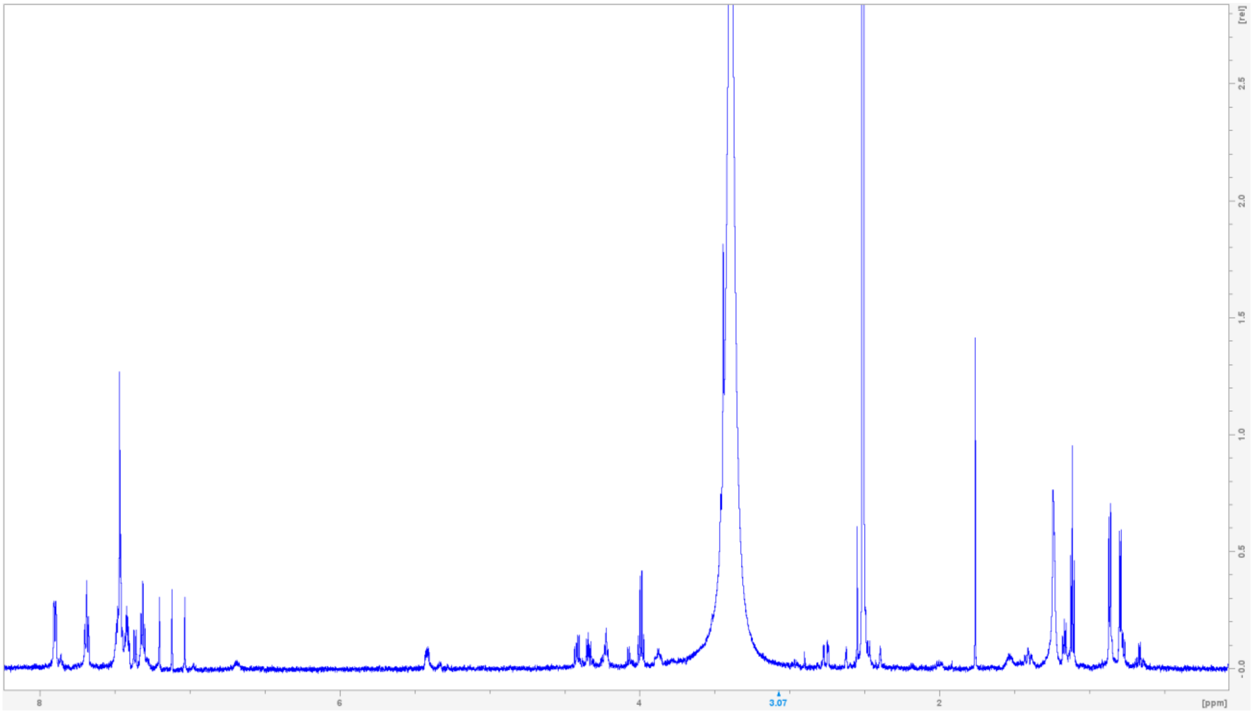

[1]H-[1]H COSY NMR spectrum of (R) MTPA ester of Fmoc-(S)-statine ethyl ester (11) in DMSO-$d_6$.

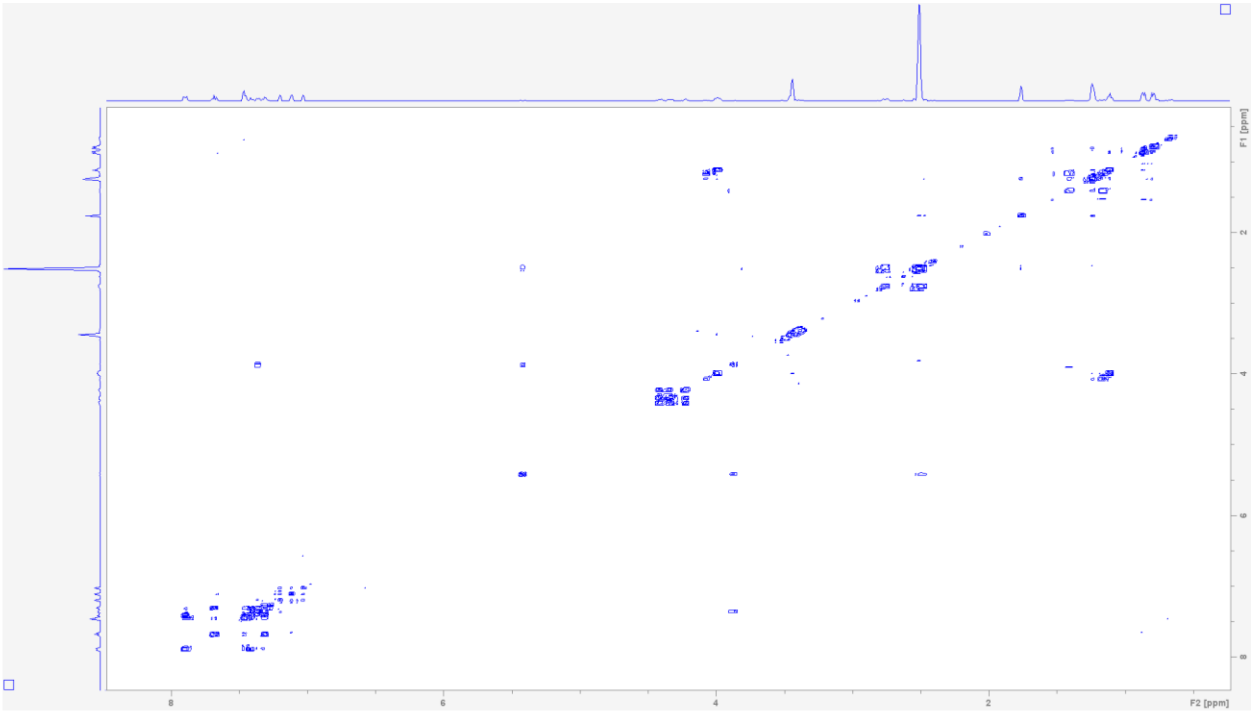

**Table S11. NMR spectroscopic data for natural L-*N*-methylvaline-L-valine-L-leucine (2) (800 MHz in DMSO-$d_6$)[a]**

| Position | $\delta_H$ mult. | $\delta_C$ | Position | $\delta_H$ mult. | $\delta_C$ |
|---|---|---|---|---|---|
| 1 | - | - | 11 | 1.96 (m, 7) | 30.5 |
| 2 | 2.66 (d, 3.5) | 70.0 | 12 | 0.83 (d, 7) | 18.1 |
| 3 | - | 173.0 | 13 | 0.86 (ov) | 19.4 |
| 4 | 1.75 (m, 7) | 31.0 | 14 | 8.17 (b) | - |
| 5 | 0.84 (d, 7) | 19.4 | 15 | 4.08 (m) | 51.0 |
| 6 | 0.86 (d, 7) | 18.8 | 16 | - | 174.2 |
| 7 | 2.17 (s) | 34.9 | 17 | 1.47, 1.45 (ov) | 40.8 |
| 8 | 7.90 (d, 9) | - | 18 | 1.62 (b) | 24.2 |
| 9 | 4.21 (m, 9) | 57.1 | 19 | 0.80 (d, 6.5) | 21.6 |
| 10 | - | 170.3 | 20 | 0.87 (d, 6.5) | 23.0 |

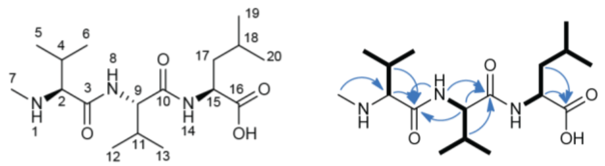

[a]Chemical shift $\delta$ and (multiplicity, *J* in Hz).

**$^1$H NMR spectrum of L-*N*-methylvaline-L-valine-L-leucine (2) in DMSO-$d_6$.**

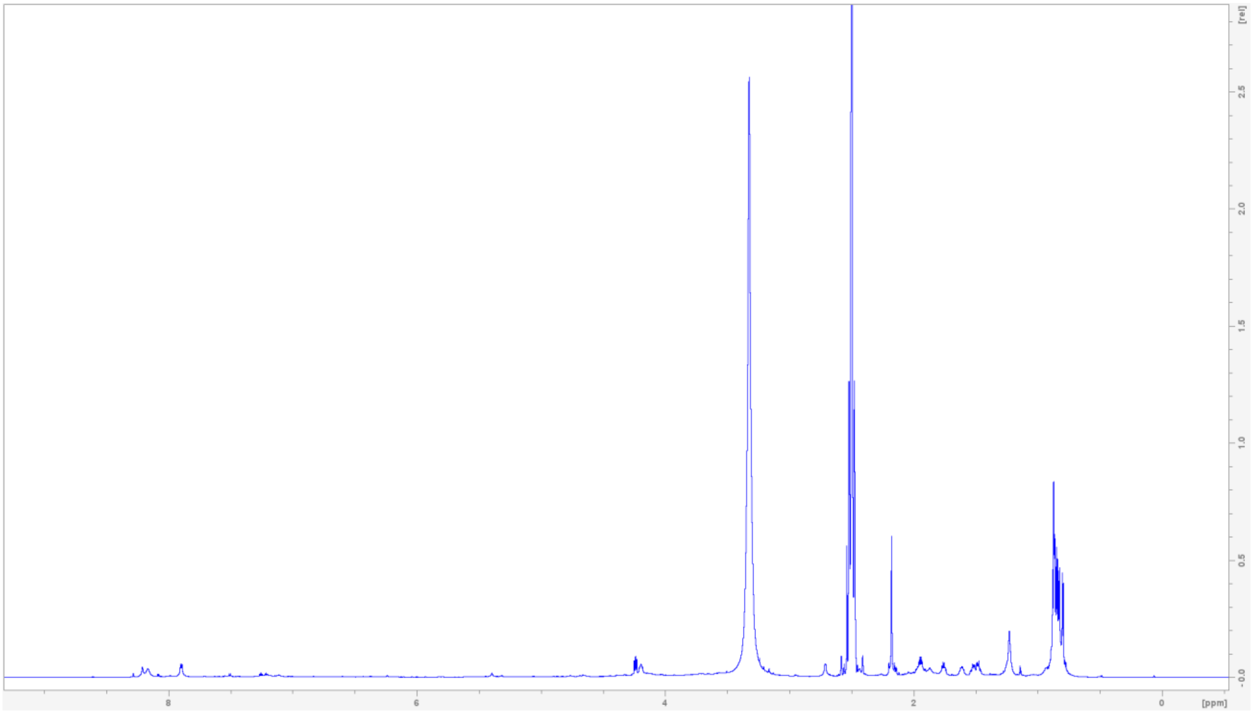

**$^{13}$C NMR spectrum of L-*N*-methylvaline-L-valine-L-leucine (2) in DMSO-*d*$_6$.**

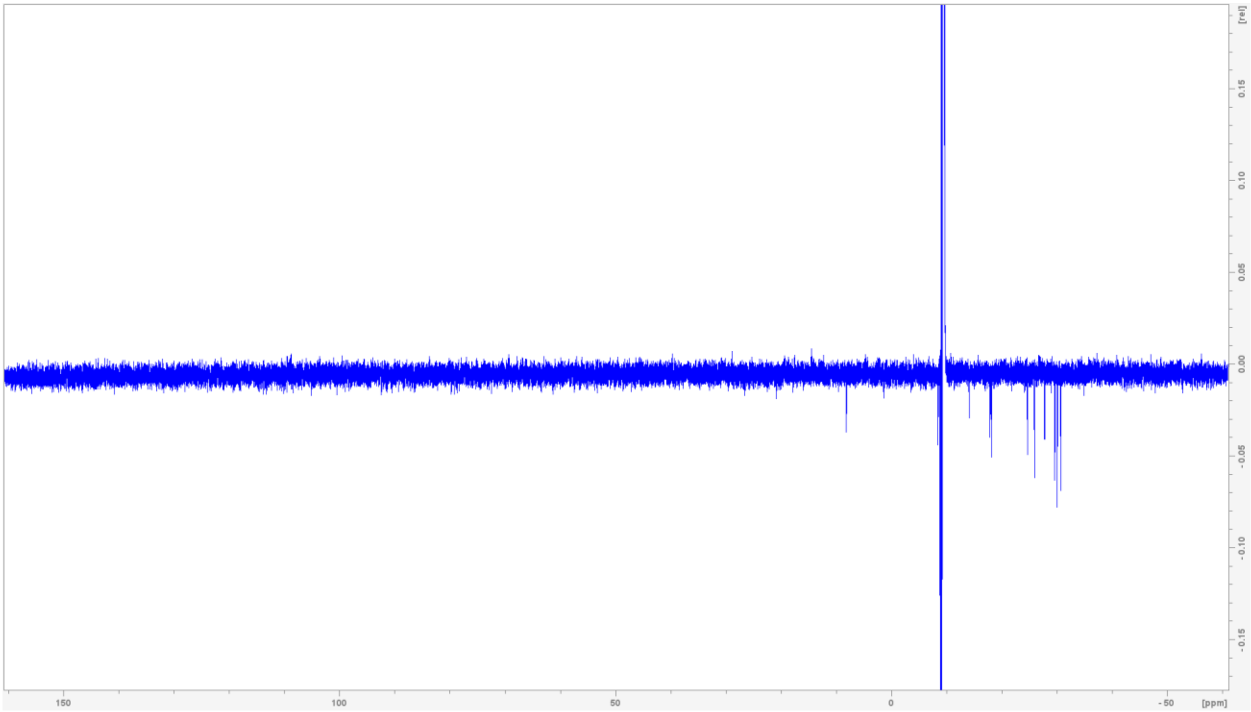


**$^1$H-$^1$H COSY NMR spectrum of L-*N*-methylvaline-L-valine-L-leucine (2) in DMSO-*d*$_6$**

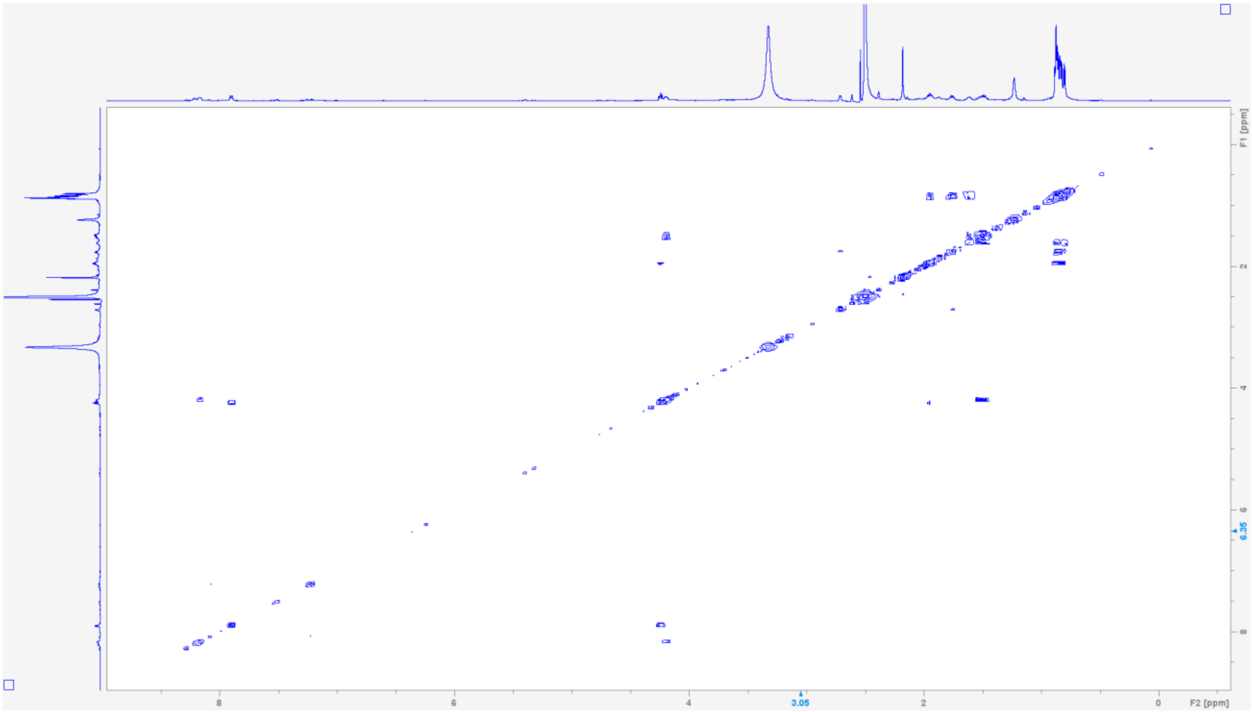

$^{1}H$-$^{13}C$ **HSQC NMR spectrum of L-*N*-methylvaline-L-valine-L-leucine (2) in DMSO-$d_6$.**

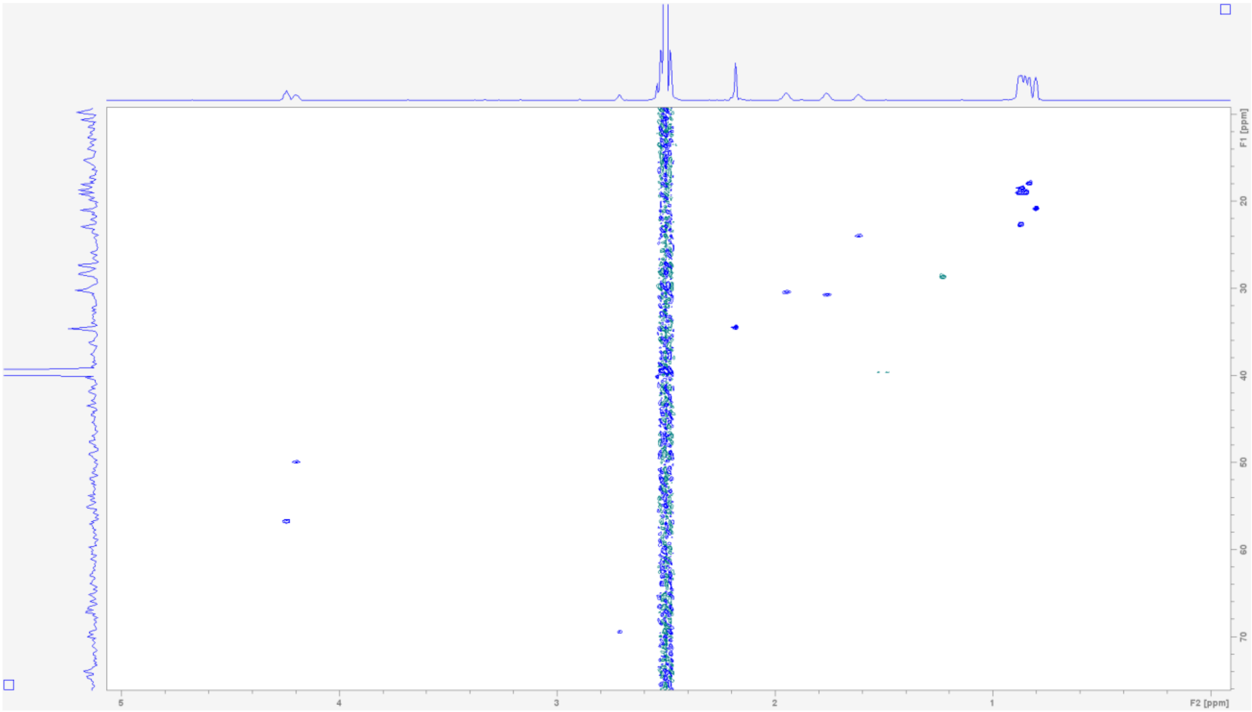


$^{1}H$-$^{13}C$ **HMBC NMR spectrum of L-*N*-methylvaline-L-valine-L-leucine (2) in DMSO-$d_6$.**

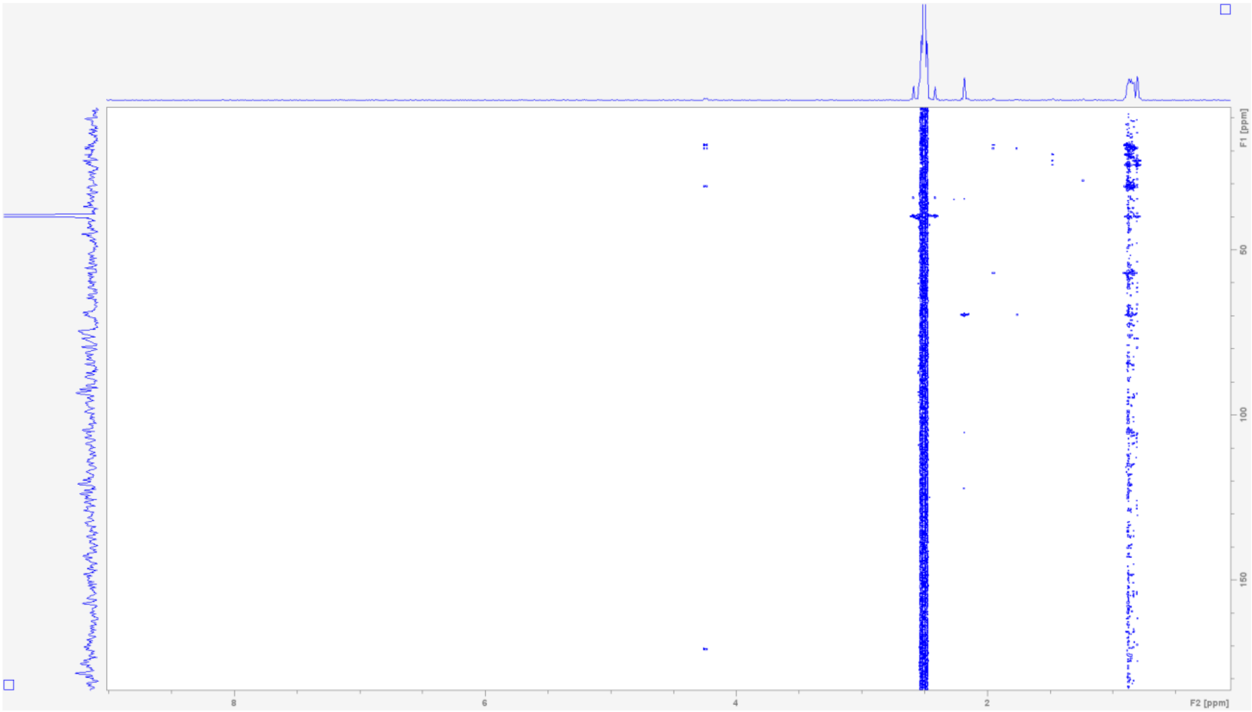

**Table S12. NMR spectroscopic data for natural gammanonin (1)(800 MHz in DMSO-$d_6$)[a]**

| Position | $\delta_H$ mult. | $\delta_C$ | Position | $\delta_H$ mult. | $\delta_C$ |
|---|---|---|---|---|---|
| 1 | - | - | 12 | 0.84 (ov) | 18.8 |
| 2 | 2.71 (d, 6.5) | 70.0 | 13 | 0.87 (ov) | 19.8 |
| 3 | - | 173.0 | 14 | 7.68 (d, 9) | - |
| 4 | 1.76 (m, 6.5) | 31.6 | 15 | 3.73 (m) | 51.9 |
| 5 | 0.87 (d, 6.5) | 19.3 | 16 | 3.39 (ov) | 71.0 |
| 6 | 0.88 (d, 6.5) | 19.9 | 17 | 1.56, 1.40 (ov) | 37.0 |
| 7 | 2.17 (s) | 35.3 | 18 | 3.50, 3.47 (ov) | 58.7 |
| 8 | 7.88 (d, 9) | - | 19 | 1.32, 1.30 (ov) | 39.3 |
| 9 | 4.20 (dd, 9) | 58.0 | 20 | 1.51 (m) | 24.5 |
| 10 | - | 171.0 | 21 | 0.84 (d, 6.5) | 24.4 |
| 11 | 1.93 (m, 7) | 31.1 | 22 | 0.78 (d, 6.5) | 21.8 |

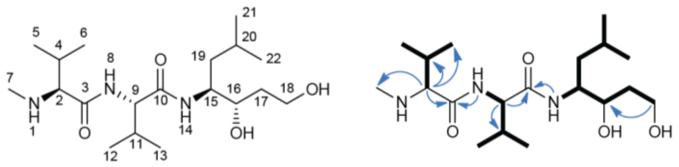

[a]Chemical shift $\delta$ and (multiplicity, $J$ in Hz).

**$^1$H NMR spectrum of gammanonin (1) in DMSO-$d_6$.**

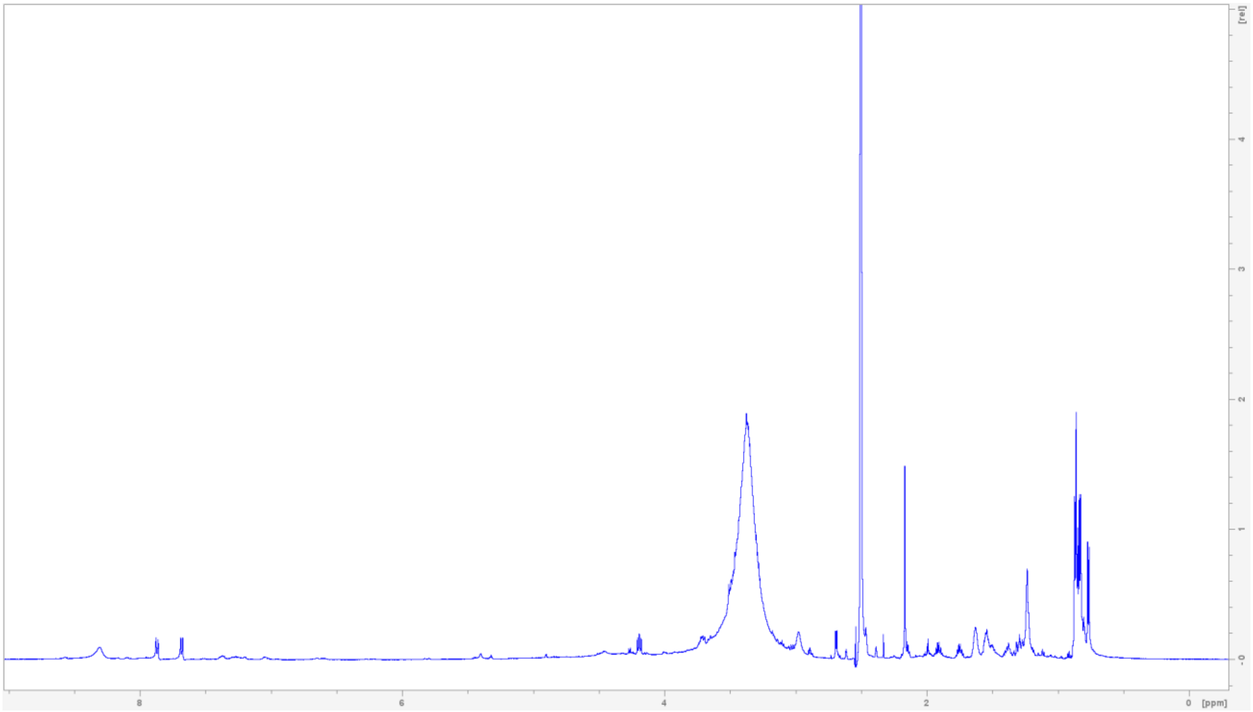

**$^{1}$H-$^{1}$H COSY NMR spectrum of gammanonin (1) in DMSO-$d_6$**

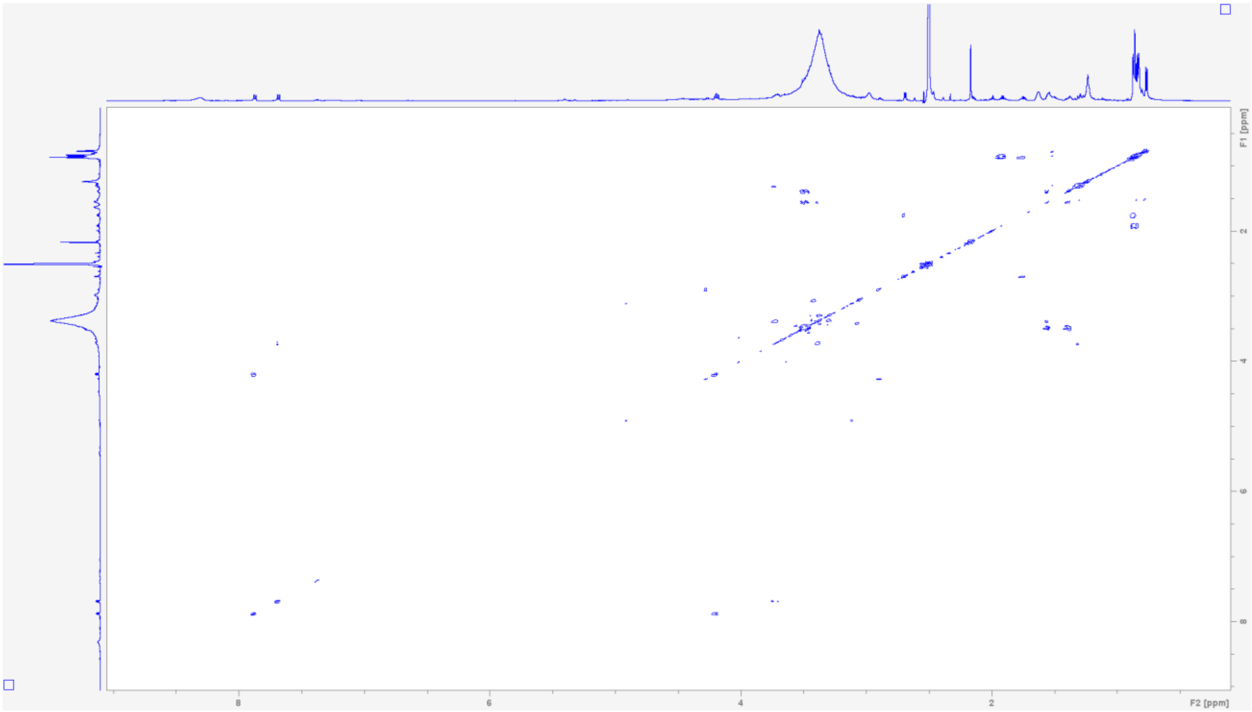


**$^{1}$H-$^{1}$H TOCSY NMR spectrum of gammanonin (1) in DMSO-$d_6$**

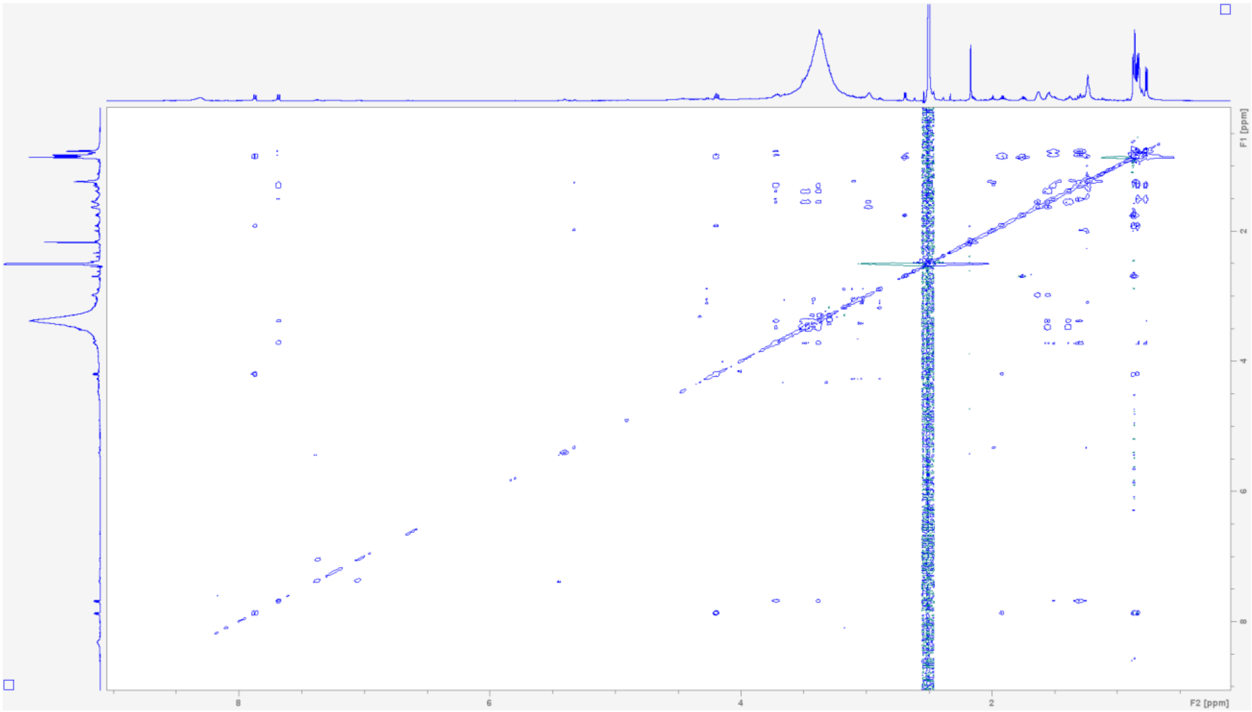

$^{1}H$-$^{13}C$ HSQC NMR spectrum of gammanonin (1) in DMSO-$d_6$.

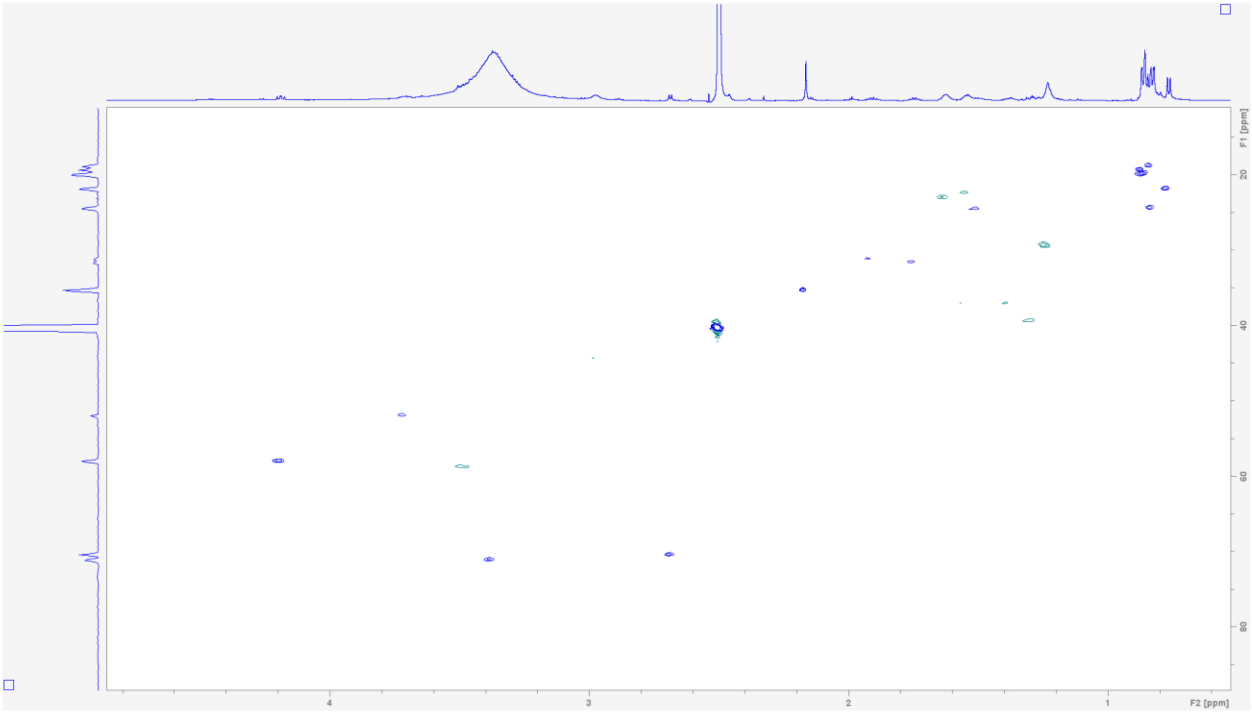


$^{1}H$-$^{13}C$ HMBC NMR spectrum of gammanonin (1) in DMSO-$d_6$.

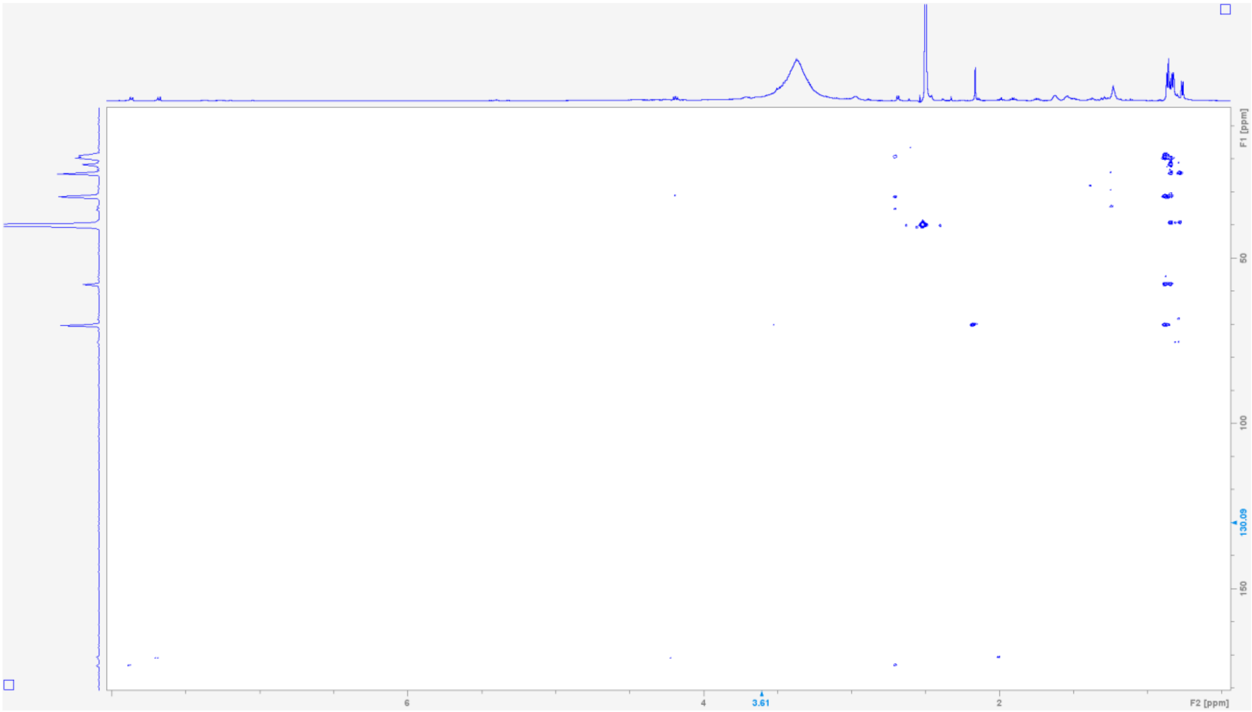

**Table S13. NMR spectroscopic data for L-valine-N-acetyl-cysteamine (12) (600 MHz in DMSO-$d_6$)[a]**

| Position | $\delta_H$ mult. | $\delta_C$ | Position | $\delta_H$ mult. | $\delta_C$ |
|---|---|---|---|---|---|
| 1 | - | - | 7 | 3.00 (m, 6.5) | 28.7 |
| 2 | 3.95 (d, 4.5) | 64.6 | 8 | 3.21 (m, 6.5) | 38.5 |
| 3 | - | 198.3 | 9 | 8.15 (t, 5.5) | - |
| 4 | 2.16 (m, 6.5) | 30.8 | 10 | - | 169.9 |
| 5 | 0.97 (d, 7) | 18.7 | 11 | 1.79 (s) | 23.1 |
| 6 | 0.93 (d, 7) | 17.7 | | | |

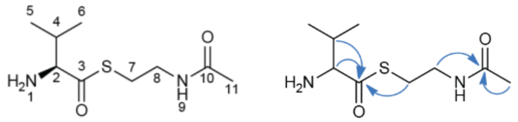

[a]Chemical shift $\delta$ and (multiplicity, $J$ in Hz).

**$^1$H NMR spectrum of L-valine-N-acetyl-cysteamine (12) in DMSO-$d_6$.**

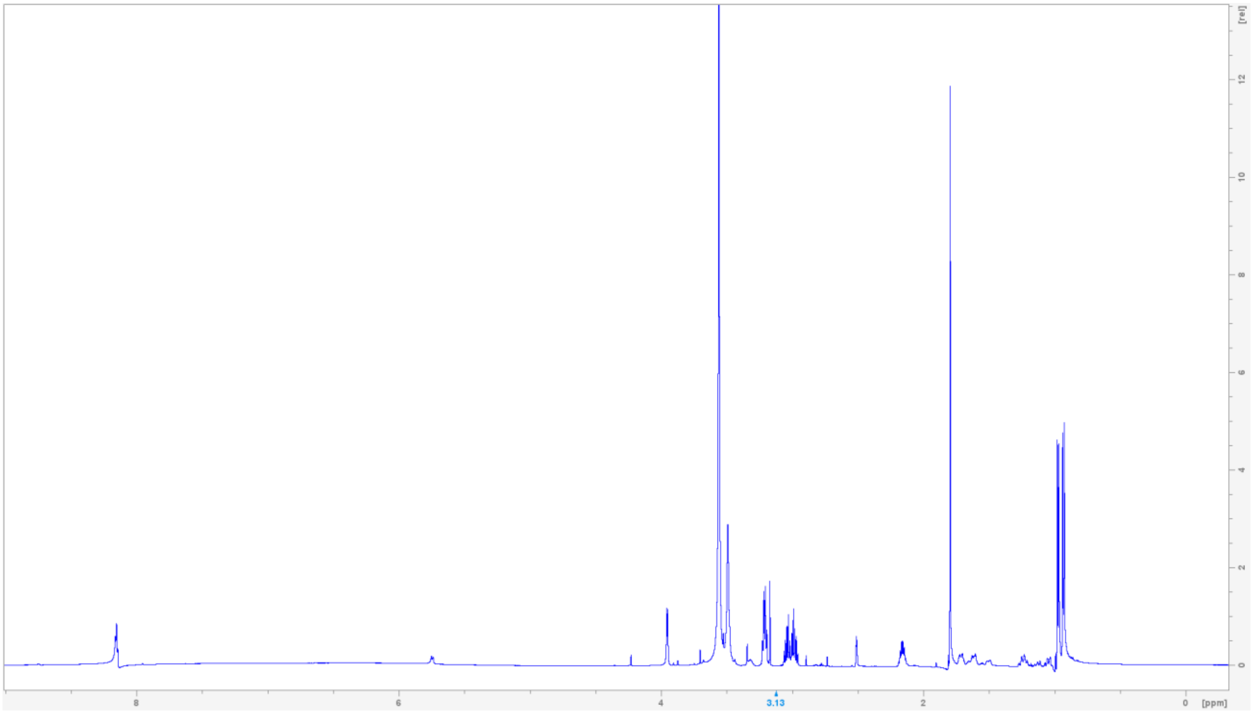

**$^{13}C$ NMR spectrum of L-valine-N-acetyl-cysteamine (12) in DMSO-$d_6$.**

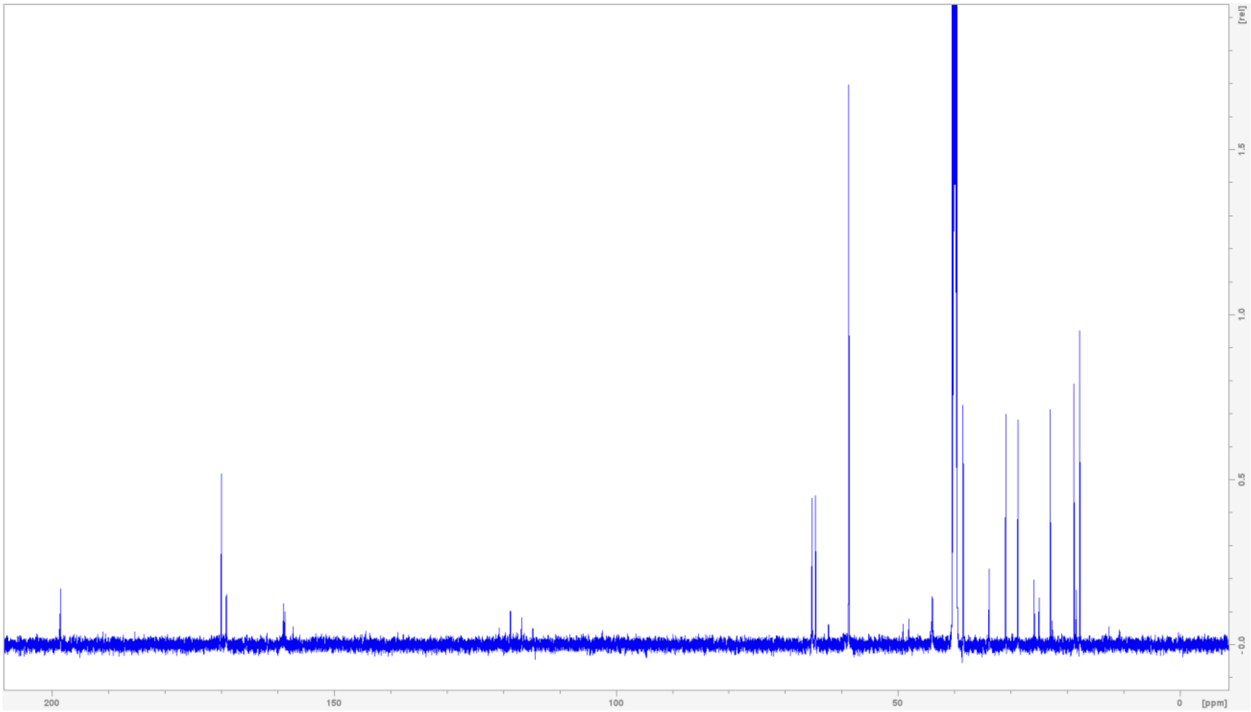


**$^{1}H$-$^{13}C$ HSQC NMR spectrum of L-valine-N-acetyl-cysteamine (12) in DMSO-$d_6$.**

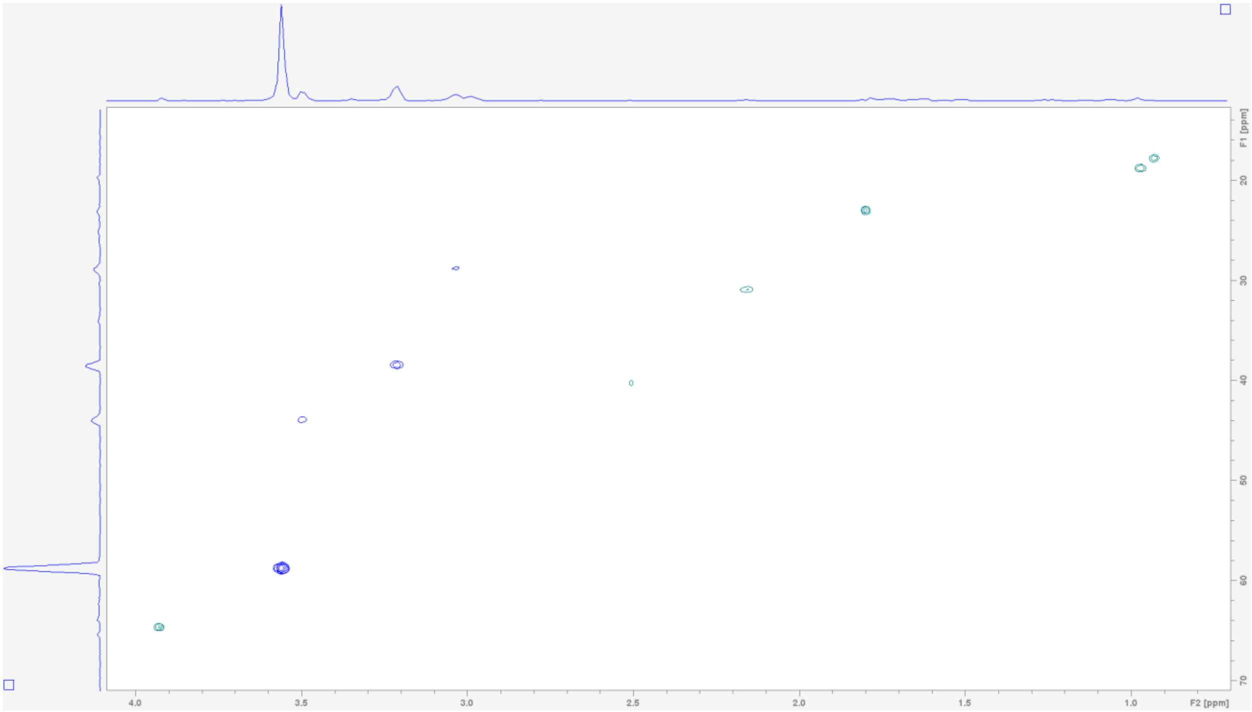

**$^{1}H$-$^{13}C$ HMBC NMR spectrum of L-valine-N-acetyl-cysteamine (12) in DMSO-$d_6$.**

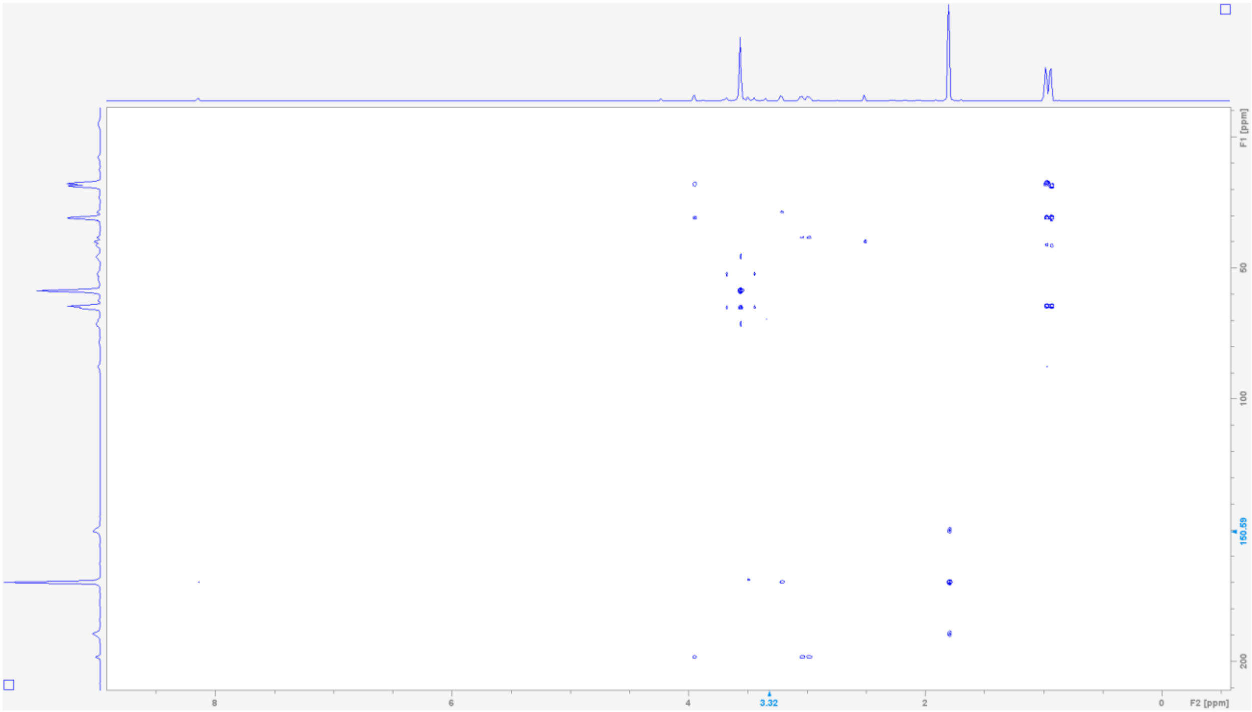

**Table S14. NMR spectroscopic data for L-valine-L-valine-N-acetyl-cysteamine (13) (600 MHz in DMSO-$d_6$)[a]**

| Position | $\delta_H$ mult. | $\delta_C$ | Position | $\delta_H$ mult. | $\delta_C$ |
|---|---|---|---|---|---|
| 1 | - | - | 10 | 2.17 (m, 6.5) | 30.5 |
| 2 | 3.82 (d, 4.5) | 57.5 | 11 | 0.92 (d, 7) | 19.5 |
| 3 | - | 168.6 | 12 | 1.01 (d, 7) | 19.1 |
| 4 | 2.22 (m, 7) | 30.2 | 13 | 2.92 (m, 6.5) | 28.3 |
| 5 | 0.91 (d, 7) | 18.2 | 14 | 3.14 (m, 7) | 38.6 |
| 6 | 0.94 (d, 7) | 17.5 | 15 | 8.05 (t, 5.5) | - |
| 7 | 8.81 (d, 8) | - | 16 | - | 169.7 |
| 8 | 4.35 (dd, 8) | 64.9 | 17 | 1.78 (s) | 22.3 |
| 9 | - | 199.8 | | | |

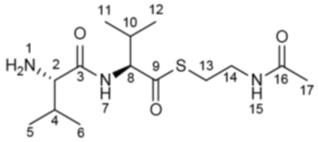

[a]Chemical shift $\delta$ and (multiplicity, $J$ in Hz).

$^{1}H$ NMR spectrum of L-valine-L-valine-N-acetyl-cysteamine (13) in DMSO-$d_6$.

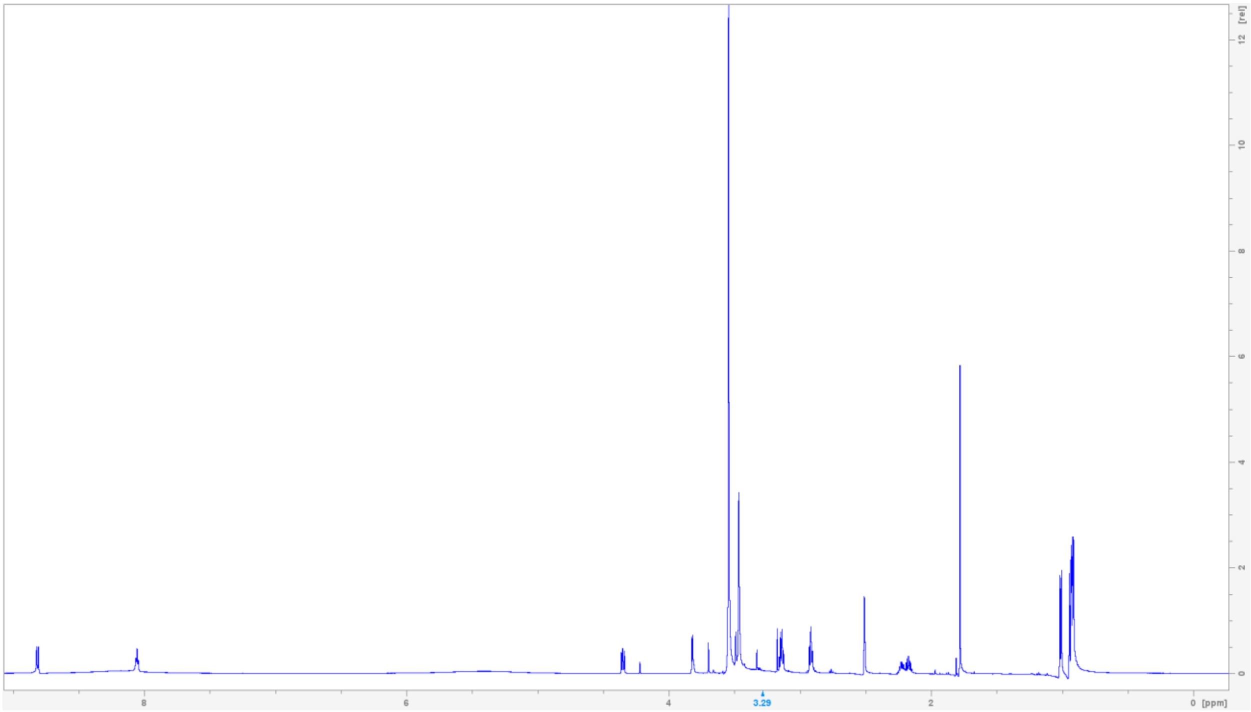


$^{13}C$ NMR spectrum of L-valine-L-valine-N-acetyl-cysteamine (13) in DMSO-$d_6$.

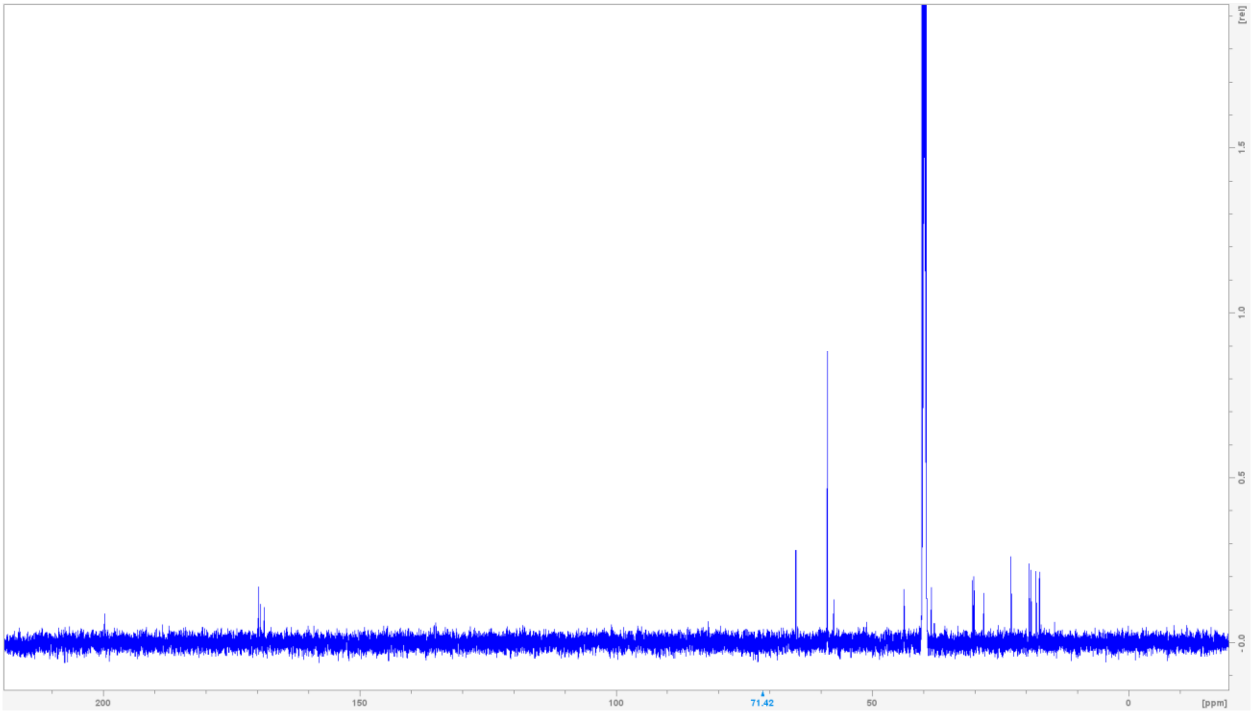

**$^{1}H$-$^{13}C$ HSQC NMR spectrum of L-valine-L-valine-N-acetyl-cysteamine (13) in DMSO-$d_6$.**

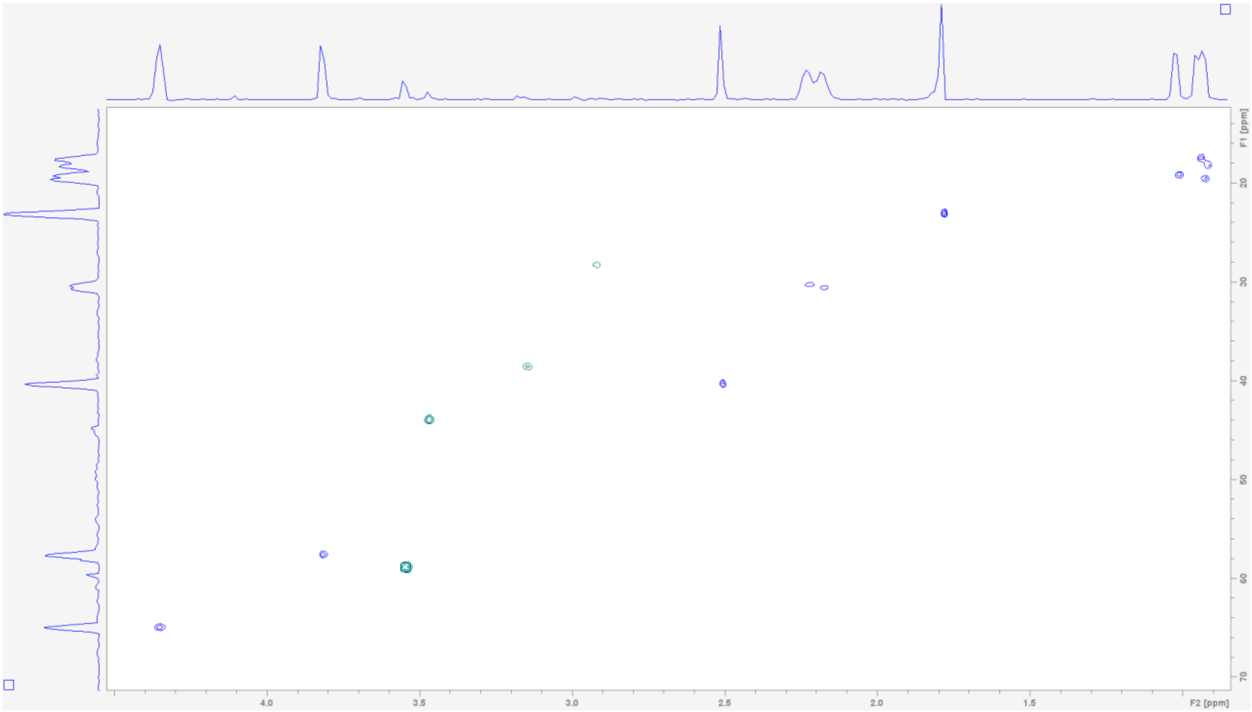

**Table S15. NMR spectroscopic data for L-valine-L-valine-L-leucine-N-acetyl-cysteamine (14) (600 MHz in DMSO-$d_6$)[a]**

| Position | $\delta_H$ mult. | $\delta_C$ | Position | $\delta_H$ mult. | $\delta_C$ |
|---|---|---|---|---|---|
| 1 | - | - | 13 | 8.44 (d, 8.5) | - |
| 2 | 3.73 (d, 5.5) | 57.4 | 14 | 4.41 (m, 4) | 57.9 |
| 3 | - | 168.3 | 15 | - | 201.5 |
| 4 | 2.03 (m, 7) | 30.5 | 16 | 1.57, 1.51 (ov) | 40.3 |
| 5 | 0.90 (d, 7) | 18.2 | 17 | 1.62 (ov) | 24.5 |
| 6 | 0.91 (d, 7) | 18.5 | 18 | 0.88 (d, 6.5) | 23.4 |
| 7 | 8.72 (d, 8) | - | 19 | 0.78 (d, 6.5) | 21.2 |
| 8 | 4.29 (dd, 7) | 58.4 | 20 | 2.86 (t, 7) | 28.1 |
| 9 | - | 171.6 | 21 | 3.12 (m, 7) | 38.5 |
| 10 | 2.07 (m, 7) | 30.9 | 22 | 8.05 (t, 5.5) | - |
| 11 | 0.97 (d, 7) | 19.6 | 23 | - | 169.7 |
| 12 | 0.92 (d, 7) | 18.9 | 24 | 1.78 (s) | 23.0 |

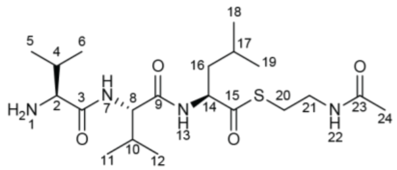

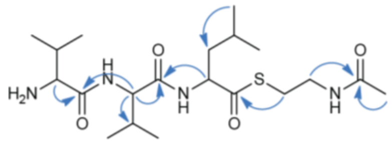

[a]Chemical shift $\delta$ and (multiplicity, $J$ in Hz).

**$^1$H NMR spectrum of L-valine-N-acetyl-cysteamine (14) in DMSO-$d_6$.**

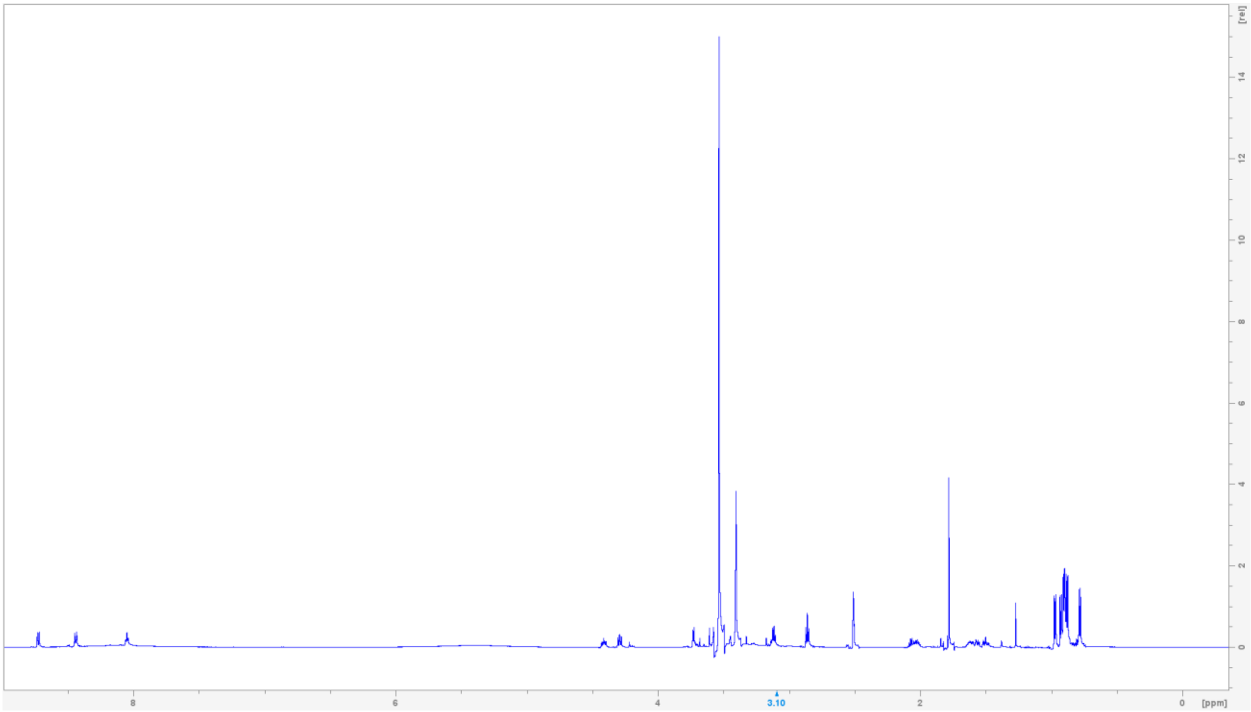

**$^{13}$C NMR spectrum of L-valine-N-acetyl-cysteamine (14) in DMSO-$d_6$.**

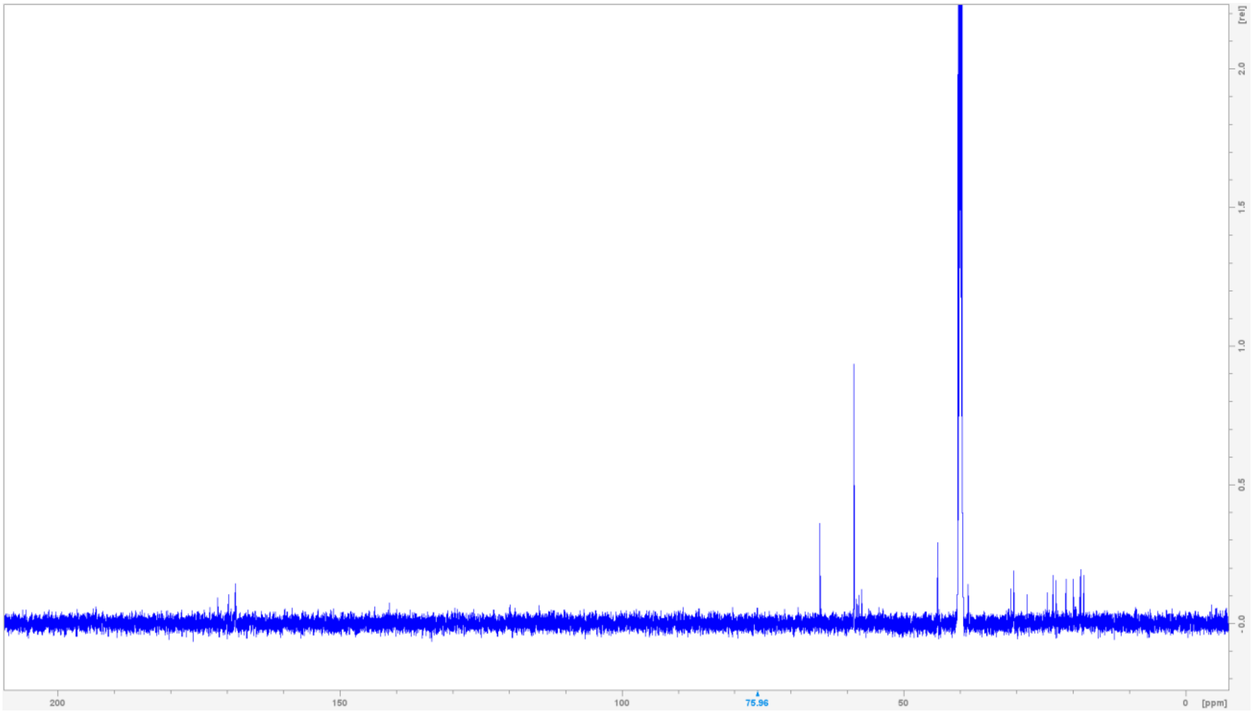

$^{1}H$-$^{13}C$ **HSQC NMR spectrum of L-valine-N-acetyl-cysteamine (14) in DMSO-*d*$_6$.**

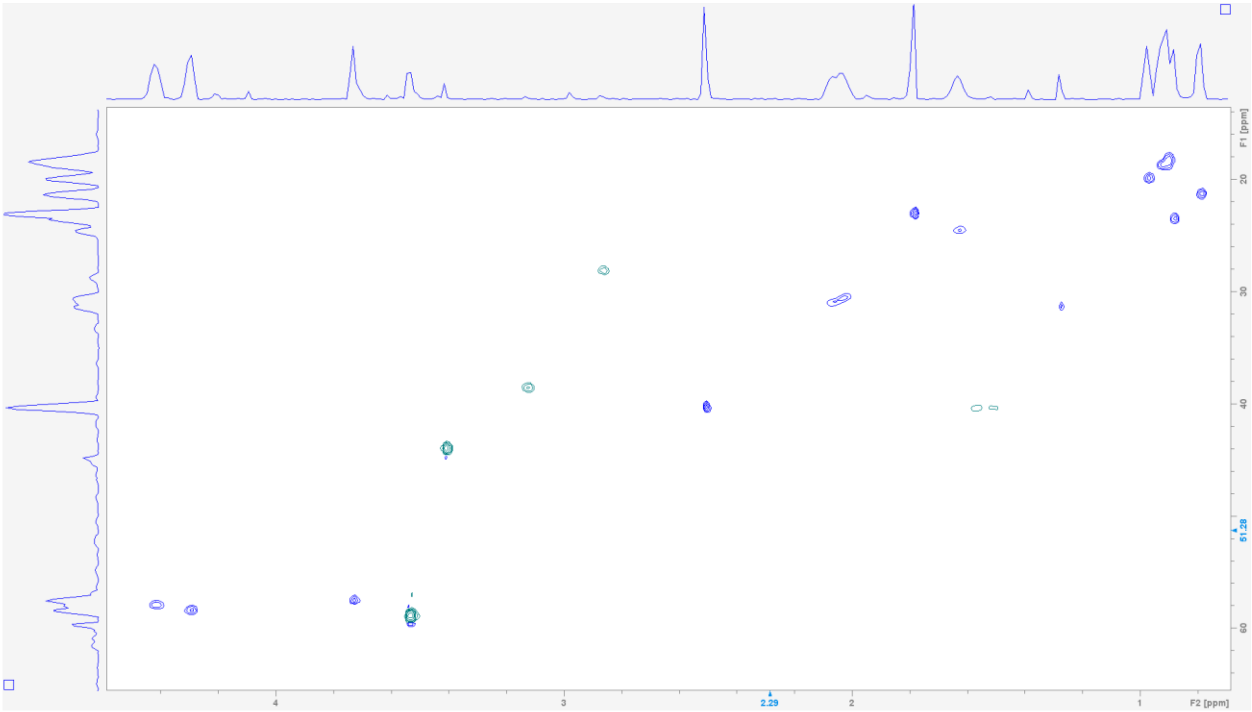

$^{1}H$-$^{13}C$ **HMBC NMR spectrum of L-valine-N-acetyl-cysteamine (14) in DMSO-*d*$_6$.**

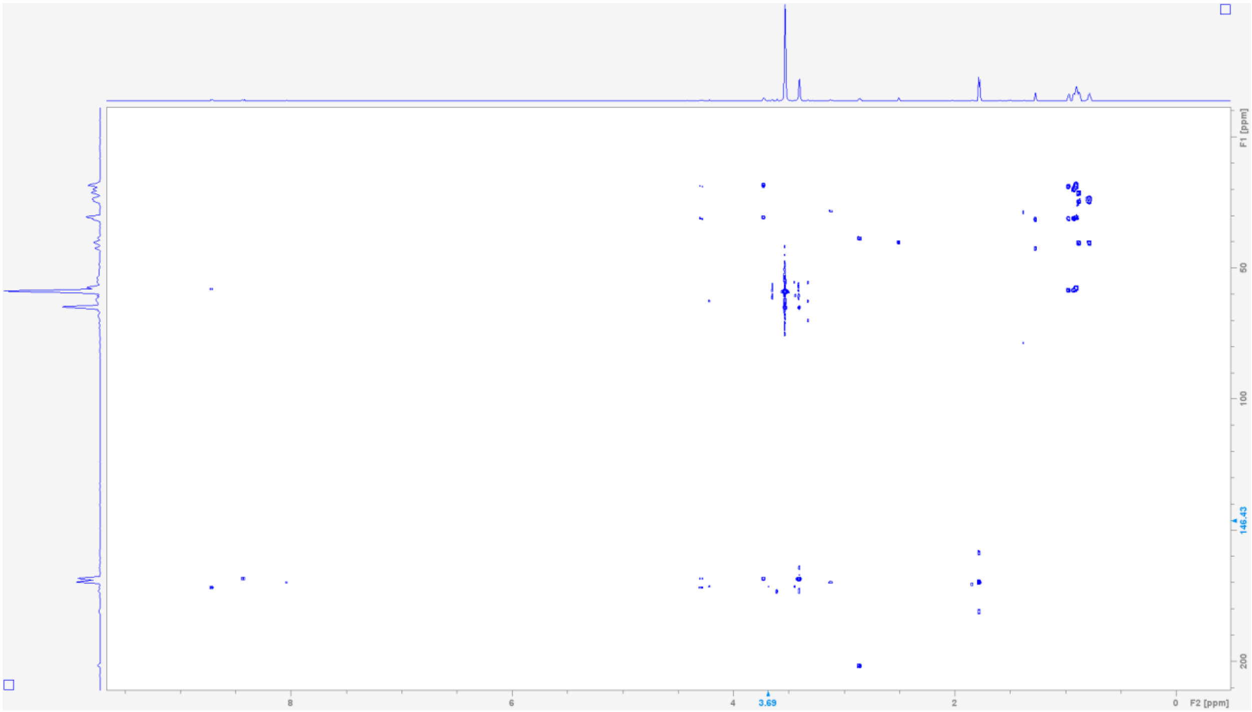

### References


(1) Johnston, C. W.; Badran, A. H.; Collins, J. J. Continuous Bioactivity-Dependent Evolution of an Antibiotic Biosynthetic Pathway. *Nat Commun* **2020**, *11* (1), 4202.

(2) Pfeifer, B. A.; Admiraal, S. J.; Gramajo, H.; Cane, D. E.; Khosla, C. Biosynthesis of Complex Polyketides in a Metabolically Engineered Strain of *E. Coli*. *Science* **2001**, *291* (5509), 1790–1792.

(3) Hadjithomas, M.; Chen, I.-M. A.; Chu, K.; Huang, J.; Ratner, A.; Palaniappan, K.; Andersen, E.; Markowitz, V.; Kyrpides, N. C.; Ivanova, N. N. IMG-ABC: New Features for Bacterial Secondary Metabolism Analysis and Targeted Biosynthetic Gene Cluster Discovery in Thousands of Microbial Genomes. *Nucleic Acids Res* **2017**, *45* (D1), D560–D565.

(4) Palaniappan, K.; Chen, I.-M. A.; Chu, K.; Ratner, A.; Seshadri, R.; Kyrpides, N. C.; Ivanova, N. N.; Mouncey, N. J. IMG-ABC v.5.0: An Update to the IMG/Atlas of Biosynthetic Gene Clusters Knowledgebase. *Nucleic Acids Research* **2020**, *48* (D1), D422–D430.

(5) Blin, K.; Shaw, S.; Kloosterman, A. M.; Charlop-Powers, Z.; van Wezel, G. P.; Medema, M. H.; Weber, T. antiSMASH 6.0: Improving Cluster Detection and Comparison Capabilities. *Nucleic Acids Research* **2021**, *49* (W1), W29–W35.

(6) Navarro-Muñoz, J. C.; Selem-Mojica, N.; Mullowney, M. W.; Kautsar, S. A.; Tryon, J. H.; Parkinson, E. I.; De Los Santos, E. L. C.; Yeong, M.; Cruz-Morales, P.; Abubucker, S.; Roeters, A.; Lokhorst, W.; Fernandez-Guerra, A.; Cappelini, L. T. D.; Goering, A. W.; Thomson, R. J.; Metcalf, W. W.; Kelleher, N. L.; Barona-Gomez, F.; Medema, M. H. A Computational Framework to Explore Large-Scale Biosynthetic Diversity. *Nat Chem Biol* **2020**, *16* (1), 60–68.

(7) Cordero, O. X.; Wildschutte, H.; Kirkup, B.; Proehl, S.; Ngo, L.; Hussain, F.; Le Roux, F.; Mincer, T.; Polz, M. F. Ecological Populations of Bacteria Act as Socially Cohesive Units of Antibiotic Production and Resistance. *Science* **2012**, *337* (6099), 1228–1231.

(8) Aron, A. T.; Gentry, E. C.; McPhail, K. L.; Nothias, L.-F.; Nothias-Esposito, M.; Bouslimani, A.; Petras, D.; Gauglitz, J. M.; Sikora, N.; Vargas, F.; Van Der Hooft, J. J. J.; Ernst, M.; Kang, K. B.; Aceves, C. M.; Caraballo-Rodríguez, A. M.; Koester, I.; Weldon, K. C.; Bertrand, S.; Roullier, C.; Sun, K.; Tehan, R. M.; Boya P., C. A.; Christian, M. H.; Gutiérrez, M.; Ulloa, A. M.; Tejeda Mora, J. A.; Mojica-Flores, R.; Lakey-Beitia, J.; Vásquez-Chaves, V.; Zhang, Y.; Calderón, A. I.; Tayler, N.; Keyzers, R. A.; Tugizimana, F.; Ndlovu, N.; Aksenov, A. A.; Jarmusch, A. K.; Schmid, R.; Truman, A. W.; Bandeira, N.; Wang, M.; Dorrestein, P. C. Reproducible Molecular Networking of Untargeted Mass Spectrometry Data Using GNPS. *Nat Protoc* **2020**, *15* (6), 1954–1991.

(9) Adusumilli, R.; Mallick, P. Data Conversion with ProteoWizard msConvert. In *Proteomics: Methods and Protocols*; Comai, L., Katz, J. E., Mallick, P., Eds.; Springer New York: New York, NY, 2017; pp 339–368.

(10) Shannon, P.; Markiel, A.; Ozier, O.; Baliga, N. S.; Wang, J. T.; Ramage, D.; Amin, N.; Schwikowski, B.; Ideker, T. Cytoscape: A Software Environment for Integrated Models of Biomolecular Interaction Networks. *Genome Res.* **2003**, *13* (11), 2498–2504.

(11) Yang, J.; Wencewicz, T. A. *In Vitro* Reconstitution of Fimsbactin Biosynthesis from *Acinetobacter Baumannii*. *ACS Chem. Biol.* **2022**, *17* (10), 2923–2935.

(12) Xu, F.; Butler, R.; May, K.; Rexhepaj, M.; Yu, D.; Zi, J.; Chen, Y.; Liang, Y.; Zeng, J.; Hevel, J.; Zhan, J. Modified Substrate Specificity of a Methyltransferase Domain by Protein Insertion into an Adenylation Domain of the Bassianolide Synthetase. *J Biol Eng* **2019**, *13* (1), 65.

(13) Ehmann, D. E.; Trauger, J. W.; Stachelhaus, T.; Walsh, C. T. Aminoacyl-SNACs as Small-Molecule Substrates for the Condensation Domains of Nonribosomal Peptide Synthetases. *Chemistry & Biology* **2000**, *7* (10), 765–772.

(14) Harnden, K. A.; Roy, A.; Hosseinzadeh, P. Overview of Methods for Purification and Characterization of Metalloproteins. *Current Protocols* **2021**, *1* (8), e234.

(15) Reddi, B.; Kishor, C.; Jangam, A.; Bala, S.; Rajeswari Batchu, U.; Gundla, R.; Addlagatta, A. Regioselectivity in Inhibition of Peptide Deformylase from Haemophilus Influenzae by 4- vs 5-Azaindole Hydroxamic Acid Derivatives: Biochemical, Structural and Antimicrobial Studies. *Bioorganic Chemistry* **2022**, *128*, 106095.

(16) Yang, N.; Sun, C. The Inhibition and Resistance Mechanisms of Actinonin, Isolated from Marine Streptomyces Sp. NHF165, against Vibrio Anguillarum. *Front. Microbiol.* **2016**, *7*.

(17) Du, Y. E.; Cui, J.; Cho, E.; Hwang, S.; Jang, Y.-J.; Oh, K.-B.; Nam, S.-J.; Oh, D.-C. Serratiomycins D1–D3, Antibacterial Cyclic Peptides from a *Serratia* Sp. and Structure Revision of Serratiomycin. *J. Nat. Prod.* **2024**, *87* (5), 1330–1337.

(18) Rittle, K. E.; Homnick, C. F.; Ponticello, G. S.; Evans, B. E. A Synthesis of Statine Utilizing an Oxidative Route to Chiral .Alpha.-Amino Aldehydes. *J. Org. Chem.* **1982**, *47* (15), 3016–3018.

(19) Yang, S.; Zhang, W.; Ding, N.; Lo, J.; Liu, Y.; Clare-Salzler, M. J.; Luesch, H.; Li, Y. Total Synthesis of Grassystatin A, a Probe for Cathepsin E Function. *Bioorganic & Medicinal Chemistry* **2012**, *20* (15), 4774–4780.

(20) Skinnider, M. A.; Johnston, C. W.; Gunabalasingam, M.; Merwin, N. J.; Kieliszek, A. M.; MacLellan, R. J.; Li, H.; Ranieri, M. R. M.; Webster, A. L. H.; Cao, M. P. T.; Pfeifle, A.; Spencer, N.; To, Q. H.; Wallace, D. P.; Dejong, C. A.; Magarvey, N. A. Comprehensive Prediction of Secondary Metabolite Structure and Biological Activity from Microbial Genome Sequences. *Nat Commun* **2020**, *11* (1), 6058.
